# A Pan-cancer Blueprint of the Heterogeneous Tumour Microenvironment Revealed by Single-Cell Profiling

**DOI:** 10.1101/2020.04.01.019646

**Authors:** Junbin Qian, Siel Olbrecht, Bram Boeckx, Hanne Vos, Damya Laoui, Emre Etlioglu, Els Wauters, Valentina Pomella, Sara Verbandt, Pieter Busschaert, Ayse Bassez, Amelie Franken, Marlies Vanden Bempt, Jieyi Xiong, Birgit Weynand, Yannick van Herck, Asier Antoranz, Francesca Maria Bosisio, Bernard Thienpont, Giuseppe Floris, Ignace Vergote, Ann Smeets, Sabine Tejpar, Diether Lambrechts

**Affiliations:** VIB Center for Cancer Biology, Leuven, Belgium; Laboratory for Translational Genetics, Department of Human Genetics, KU Leuven, Leuven, Belgium; Department of Obstetrics and Gynaecology, University Hospitals Leuven, Leuven, Belgium; Department of Oncology, KU Leuven, Surgical Oncology, University Hospitals Leuven, Leuven, Belgium; Laboratory of Cellular and Molecular Immunology, Vrije Universiteit Brussel, Brussels, Belgium; Laboratory of Myeloid Cell Immunology, VIB Center for Inflammation Research, Brussels, Belgium; Laboratory of Molecular Digestive Oncology, Department of Oncology, KU Leuven, Leuven, Belgium; Respiratory Oncology Unit (Pneumology) and Leuven Lung Cancer Group, University Hospital KU Leuven, Leuven, Belgium; Laboratory of Pneumology, Department of Chronic Diseases, Metabolism and Ageing, KU Leuven, Leuven, Belgium; Laboratory of Translational Cell & Tissue Research, Department of Imaging and Pathology, University Hospitals Leuven, KU Leuven, Leuven, Belgium; Laboratory of Experimental Oncology, KU Leuven, Leuven, Belgium; Laboratory for Functional Epigenetics, Department of Human Genetics, KU Leuven, Leuven, Belgium

**Keywords:** Tumour microenvironment, stromal cell heterogeneity, single-cell RNA-seq, CITE- seq, therapeutic target, clinical response

## Abstract

The stromal compartment of the tumour microenvironment consists of a heterogeneous set of tissue-resident and tumour-infiltrating cells, which are profoundly moulded by cancer cells. An outstanding question is to what extent this heterogeneity is similar between cancers affecting different organs. Here, we profile 233,591 single cells from patients with lung, colorectal, ovary and breast cancer (n=36) and construct a pan-cancer blueprint of stromal cell heterogeneity using different single-cell RNA and protein-based technologies. We identify 68 stromal cell populations, of which 46 are shared between cancer types and 22 are unique. We also characterise each population phenotypically by highlighting its marker genes, transcription factors, metabolic activities and tissue-specific expression differences. Resident cell types are characterised by substantial tissue specificity, while tumour-infiltrating cell types are largely shared across cancer types. Finally, by applying the blueprint to melanoma tumours treated with checkpoint immunotherapy and identifying a naïve CD4^+^ T-cell phenotype predictive of response to checkpoint immunotherapy, we illustrate how it can serve as a guide to interpret scRNA-seq data. In conclusion, by providing a comprehensive blueprint through an interactive web server, we generate a first panoramic view on the shared complexity of stromal cells in different cancers.

## Introduction

In recent years, single-cell RNA sequencing (scRNA-seq) studies have provided an unprecedented view on how stromal cells consist of heterogeneous and phenotypically diverse populations of cells. Indeed, by now, the tumour microenvironment (TME) of several cancer types has been profiled, including melanoma^1^, lung cancer^2^, head and neck cancer^3^, hepatocellular carcinoma^4^, glioma^5^, medulloblastoma^6^, pancreatic cancer^7^, *etc.* However, while there is still an unmet need to chart TME heterogeneity in additional tumours and cancer types, the higher-level question relates to the similarities between these microenvironments.

Indeed, it remains unexplored whether the same stromal cell phenotypes are present in different cancer types. Also, it is not clear to what extent these phenotypes are reminiscent of the normal tissue from which they originate and are thus characterised by tissue-specific expression. Such knowledge is highly desirable, because it not only facilitates comparison between different scRNA-seq studies, but also contributes to our insights in cancer type-specific gene expression patterns and treatment vulnerabilities.

Furthermore, this knowledge would allow us to assess at single-cell level the underlying mechanisms of action of novel cancer therapies. Indeed, most innovative cancer therapies are given to cancer patients with advanced disease, in which tissue biopsies often can only be collected from metastasized organs. It is difficult, however, to systematically identify stromal phenotypes in biopsies taken from different organs, as their expression is determined by the metastasized tissue. Another challenge is that rare stromal cell phenotypes often cluster together with other more common phenotypes, and can therefore only be detected when several 10,000s of cells derived from multiple patient biopsies are profiled together. Many of these rare phenotypes are critical in determining response to cancer treatment and therefore need to be assessed as a separate population of cells. For instance, scRNA-seq of melanoma T-cells exposed to anti-PD1 identified TCF7^+^ CD8^+^ memory-precursor T-cells as the population underlying treatment response. These cells are rare, as they represent only ∼15% of CD8^+^ T-cells, which by themselves represent only ∼2.5% of cells in these tumours^8^. In order not to miss these rare phenotypes, a blueprint of the different cell populations present in each cancer type would be of considerable benefit.

We therefore generated a comprehensive blueprint of stromal cell heterogeneity across cancer types and provide a detailed view on the shared complexity and heterogeneity of stromal cells in these cancers. We illustrate how this blueprint can serve as a guide to interpret scRNA-seq data at individual patient level, even when comparing tumours collected from different tissues or profiled using different scRNA-seq technologies. Our single-cell blueprint can be visualised, analysed and downloaded from an interactive web server (http://blueprint.lambrechtslab.org).

## Results

### scRNA-seq and cell typing of tumour and normal tissue

First, we performed scRNA-seq on tumours from 3 different organs (or cancer types): colorectal cancer (CRC, n=7), lung cancer (LC, n=8) and ovarian cancer (OvC, n=5). Whenever possible, we retrieved both malignant (tumour) and matched non-malignant (normal) tissue during surgical resection with curative intent. All tumours were treatment-naïve and reflected different disease stages (e.g. stage I-IV CRC) or histopathologies (e.g. adenocarcinoma versus squamous LC), and whenever possible tissues were collected from different anatomic sites (e.g. primary tumour from the ovary and omentum in OvC, or from core *versus* border regions in CRC). Overall, 50 tumour tissues and 17 normal tissues were profiled (**Fig. 1a**). Clinical and tumour mutation data are summarised in **Tables S1-3**.

**Fig. 1.**
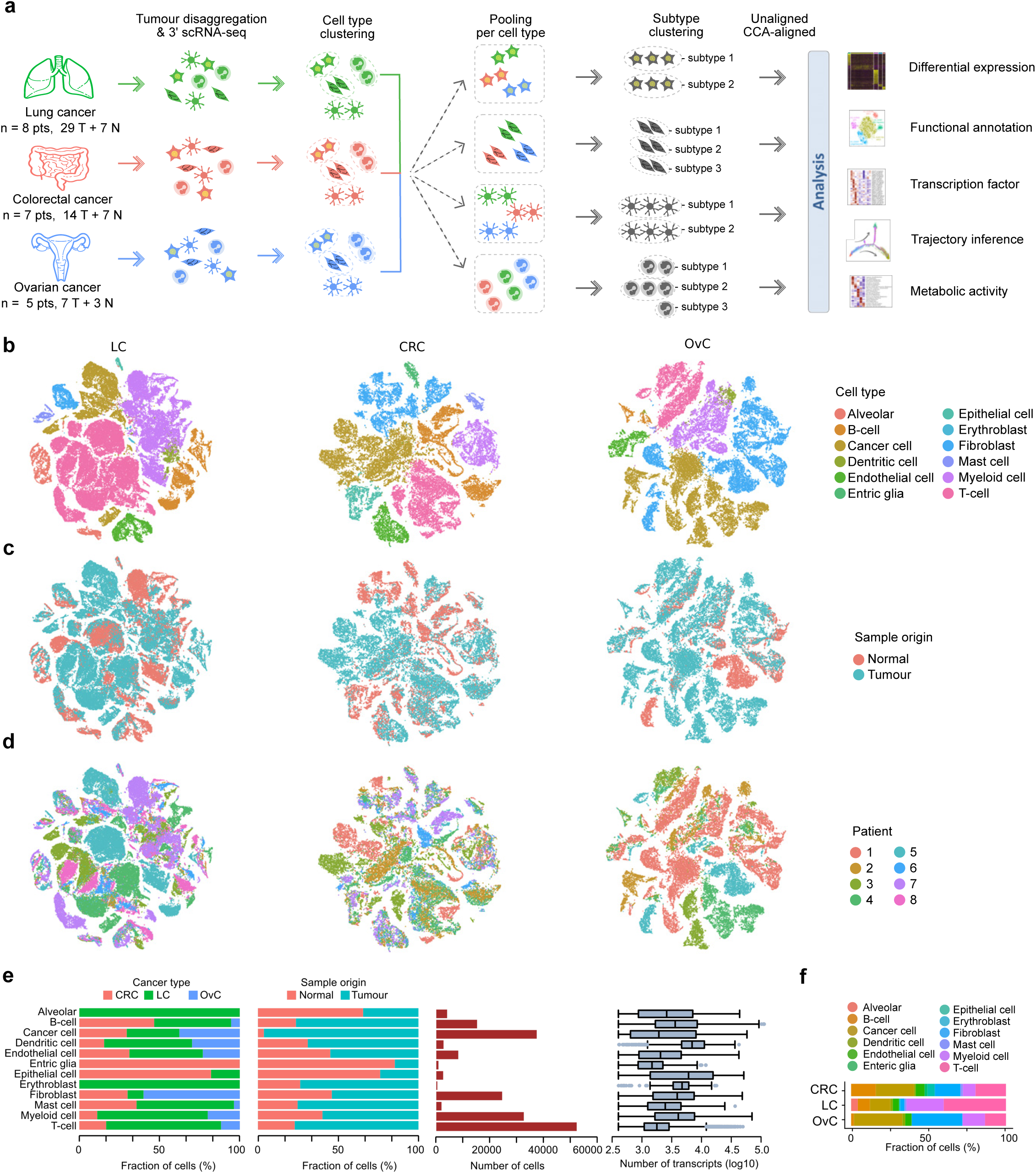
Experimental design and cell typing. **a** Analysis workflow of tumour and matched normal samples from 3 cancer types. **b-d** t-SNE representation for LC (n=93,576 cells), CRC (n=44,685) and OvC (45,115) colour-coded for cell type (**b**), sample origin (**c**) and patient (**d**). **e** Bar plots representing per cell type from left to right: the fraction of cells per tissue and per origin, the number of cells, the total number of transcripts. Dendritic cells were transcriptionally most active (*p* < 1.6×10^−10^). **f** Fraction of cells for major cell types per cancer type. T-cells were most frequent in LC (*p* < 0.0047).

Following resection, tissues were rapidly digested to a single-cell suspension and unbiasedly subjected to 3’-scRNA-seq. After quality filtering (Methods), we obtained ∼1 billion unique transcripts from 183,373 cells with >200 genes detected. Of these, 71.7% of cells originated from malignant tissue. Principle component analysis (PCA) using variably expressed genes was used to generate t-SNEs at different resolutions (Supplementary information, **Fig. S1a,b**). Marker genes were used to identify cell types (Supplementary information, **Fig. S1c**). At low resolution, cells clustered based on cancer type, whereas at high resolution they clustered based on patient identity (Supplementary information, **Fig. S1d**). Also, when assessing how cell types previously identified in LC now clustered^2^, obvious differences were noted, with similar phenotypic cells now belonging to distinct clusters.

### Sub-phenotyping of cell types

We therefore used a different strategy. First, we clustered cells for each cancer type separately and assigned cell type identities to each cell (**Fig. 1a**). This revealed that cells mostly clustered based on cell type (**Fig. 1b** and Supplementary information, **Fig. S1e**), allowing us to assess the relative contribution of tumour *versus* normal tissue or individual patients to each cell type (**Fig. 1c-e**, and Supplementary information, **Fig. S1f**). We observed that dendritic cells were transcriptionally most active, while T-cells were the most frequent cell type across cancer types (**Fig. 1e-f**), especially in LC (as observed in other datasets; Supplementary information, **Fig. S1g**). We also identified cell types specific for one cancer type, including lung alveolar, epithelial and enteric glial cells.

Next, we pooled cells from different cancer types based on cell type identity and performed PCA-based unaligned clustering, generating t-SNEs displaying the phenotypic heterogeneity for each cell type (**Fig. 1a**). For alveolar, epithelial and enteric glial cells this generated 15 tissue-specific subclusters (LC: 5 alveolar clusters and 1 epithelial cluster; CRC: 8 epithelial clusters and 1 enteric glial cluster), most of which have been described previously^9,10^ (Supplementary information, **Fig. S1h-p**). Additionally, 7 tissue-specific subclusters were identified amongst the fibroblasts and macrophages (see below). Separately, we performed canonical correlation analysis (CCA) for each cancer type followed by graph-based clustering to generate a t-SNE per cell type (**Fig. 1a**)^11^. To avoid that CCA would erroneously assign cells unique for a cancer type, we did not include any of the 22 tissue-specific subclusters. Thus, while unaligned clustering revealed patient or cancer type-specific clusters, CCA aligned common sources of variation between cancer types. Two measures to calculate sample bias (i.e., ‘Shannon index’ and ‘mixing metrics’, see Methods) confirmed that after CCA bias decreased in all clusters (Supplementary information, **Fig. S1q,r**).

Overall, we identified 68 stromal subclusters or phenotypes, of which 46 were shared across cancer types. The number of phenotypes varied between cell types, ranging between 5 to 11 for dendritic cells and fibroblasts, respectively. Our approach was less successful for cancer cells, which due to underlying genetic heterogeneity continued to cluster patient-specifically (Supplementary information, **Fig. S1s-u**). The number of cancer cells varied substantially between tumours, while also T-cells, myeloid cells and B-cells varied considerably (Supplementary information, **Fig. S1v,w**).

Below, we describe each stromal phenotype in more detail, highlighting the number of cells, read counts and transcripts across all cancer types and for each cancer type separately, both in tumour *versus* normal tissue (**Table S4**). Additionally, marker genes and functional characteristics of each phenotype are highlighted (**Table S5**). The enrichment or depletion of these phenotypes in a cancer type (LC, CRC and OvC) or tissue (tumour *versus* normal) are evaluated (**Table S6**), while gene set enrichment analysis for biological and disease pathways (REACTOME and Gene Ontology) is also performed (see http://blueprint.lambrechtslab.org).

### Endothelial cells, tissue-specificity confined to normal tissue

Clustering the transcriptomes of 8,223 endothelial cells (ECs) using unaligned and CCA-aligned approaches identified, respectively, 13 and 9 clusters, each with corresponding marker genes (**Fig. 2a-c** and Supplementary information, **Fig. S2a-c**). Five CCA-aligned clusters were shared between cancer types (**Fig. 2d,e**), including, based on marker gene expression, C1_ESM1 tip cells (*ESM1, NID2*), C2_ACKR1 high endothelial venules (HEVs) and venous ECs (*ACKR1, SELP*), C3_CA4 capillary (*CA4, CD36*), C4_FBLN5 arterial (*FBLN5, GJA5*) and C5_PROX1 lymphatic (*PROX1, PDPN*) ECs. Three other clusters displayed T-cell (C6_CD3D), pericyte (C7_RGS5) and myeloid-specific (C8_AIF1) marker genes and consisted of doublet cells, while one cluster consisted of low-quality ECs (C9; Supplementary information, **Fig. S2d,e**). Tip ECs only resided in malignant tissue and were most prevalent in CRC, while also HEVs were enriched in tumours. In contrast, capillary ECs (cECs) were enriched in normal tissue (**Fig. 2d-f**; Supplementary information, **Fig. S2f**). We identified several genes differentially expressed between tumour and normal tissue (Supplementary information, **Fig. S2g** and **Table S7**). For instance, the pro-angiogenic factor perlecan (or *HSPG2*) was highly expressed in tumour *versus* normal cECs.

**Fig. 2.**
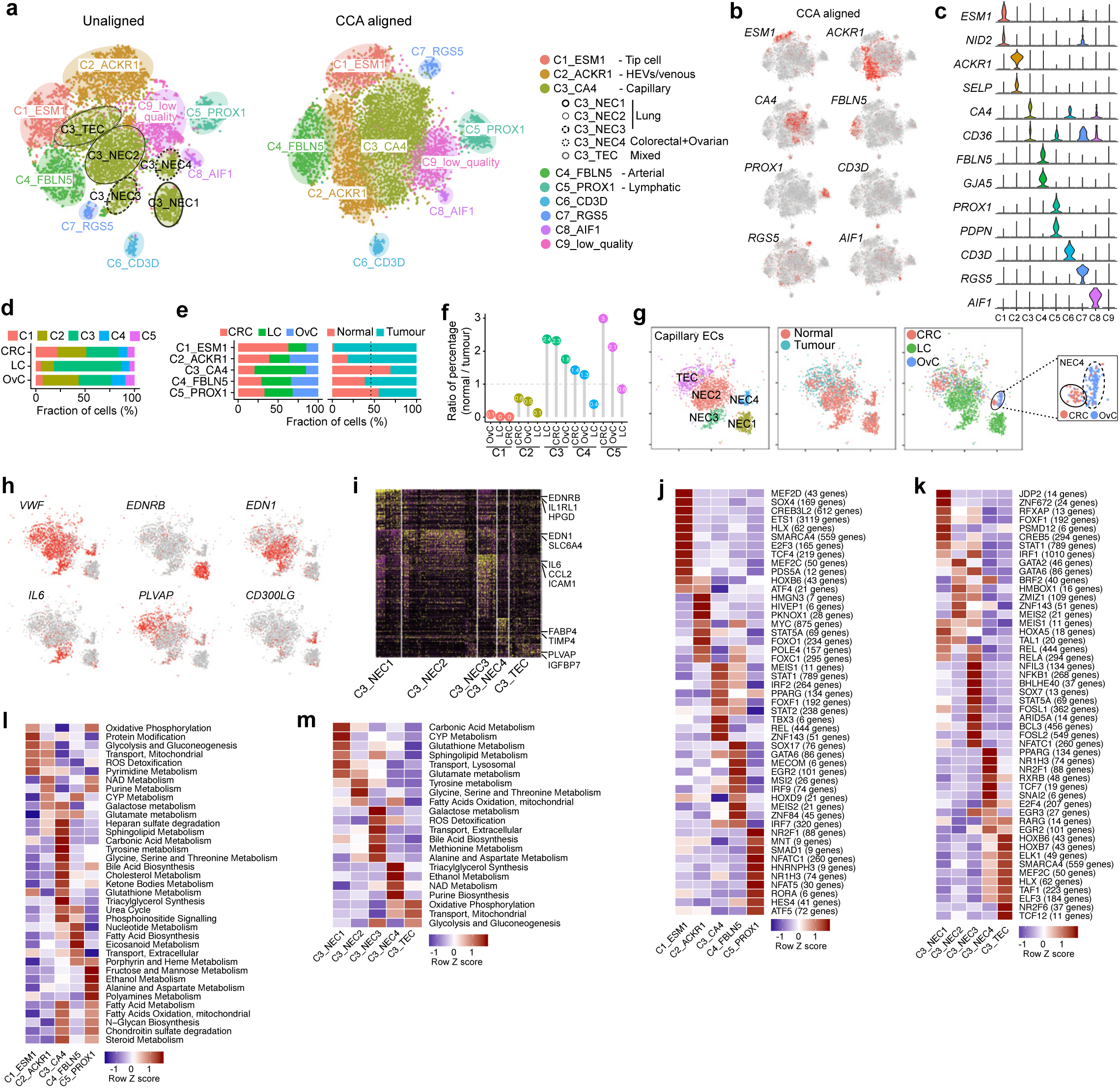
Clustering 8,223 ECs. **a** t-SNEs colour-coded for annotated ECs by unaligned and CCA aligned clustering. **b** t-SNEs with EC marker gene expression for CCA clusters. **c** Marker gene expression per EC cluster. **d** Fraction of cells in each cancer type per EC cluster. **e** Fraction of EC clusters per cancer type (left) and sample origin (right). **f** Normal/tumour ratio of relative % of EC clusters, <1 indicates tumour enrichment. Tip ECs (FDR=1.4×10^−141^) and HEVs (FDR=2.3×10^−60^) were enriched in tumour. **g** t-SNEs of cEC clusters by unaligned clustering, colour-coded by cluster, sample origin and cancer type, including a zoom-in of the NEC4 cluster (right). **h** t-SNE of marker gene expression in cEC clusters. **i-k** Heatmap of differentially expressed genes in cEC clusters (**i**), of TF activity by SCENIC for EC (**j**) or cEC clusters (**k**). **l,m** Heatmap showing metabolic activity for EC (**l**) or cEC clusters (**m**).

There were 5 unaligned cEC clusters, which clustered together (in C3_CA4) after CCA. Among these, 4 were derived from normal tissue (NEC1-4; **Fig. 2g**). Moreover, NEC1-3s were all from lung, suggesting that most cEC heterogeneity is ascribable to normal lung. C3_NEC1s represented alveolar cECs based on the absence of *VWF*, while C3_NEC2s and C3_NEC3s represented extra-alveolar cECs (**Fig. 2g-i**)^12,13^. C3_NEC1s expressed *EDNRB*, an oxygen-sensitive regulator mediating vasodilation^14^, but also IL33-receptor *IL1RL1* (ST2). This is surprising as major IL-33 effector cell types are thus far only immune cells, including basophils and innate lymphocytes^10^. Both extra-alveolar cNEC clusters expressed *EDN1*, which is a potent vasoconstrictor. C3_NEC3s additionally expressed cytokines, chemotactic and immune cell homing molecules (e.g. *IL6, CCL2, ICAM1*) (Supplementary information, **Fig. S2h)**. In contrast, C3_NEC4s were exclusively composed of ovary and colon-derived cells, suggesting similarities between NECs from both tissues. A polarized distribution of ovary and colon-derived ECs within the C3_NEC4 cluster (**Fig. 2g**) suggests, however, that there are also differences between both tissues. In contrast, tumour cECs (C3_TECs) were derived from all 3 cancer types and lacked tissue specificity on the t-SNE. Indeed, C3_TECs were all characterised by tumour EC markers *PLVAP* and *IGFBP7*^15–17^ (Supplementary information, **Fig. S2h, Table S5**), and only few genes were differentially expressed between cancer types in TECs (Supplementary information, **Fig. S2i)**.

SCENIC^18^ identified different transcription factors (TFs) underlying each EC phenotype (**Fig. 2j-k** and Supplementary information, **Table S8**). For instance, activation of NF-κB (NFKB1) and HOXB pathways was confined to C3_NEC3s and C3_TECs, respectively. Metabolic pathway analysis revealed distinct metabolic signatures among EC phenotypes (**Fig. 2l,m**): glycolysis and oxidative phosphorylation, which promote vessel sprouting^19^, were upregulated in tip cells, while fatty acid oxidation, essential for lymphangiogenesis was increased in lymphatic ECs^19^. Metabolic activities within cECs also differed: carbonic acid metabolism was most active in C3_NEC1, confirming these are alveolar cECs, which actively convert carbonic acid into CO2 during respiration. However, carbonic acid metabolism was reduced in C3_TECs, which instead deployed glycolysis and oxidative phosphorylation (Supplementary information, **Fig. S2j**). Similar characteristics were observed when assessing activation of cancer hallmark pathways (Supplementary information, **Fig. S2k,l**).

### Fibroblasts show the highest cancer type specificity

Fibroblasts are highly versatile cell types endowed with extensive heterogeneity^20^. Indeed, unaligned clustering of 24,622 fibroblasts resulted in 17 clusters (**Fig. 3a,b**), which were often tissue-specific (Supplementary information, **Fig. S3a-d**). Particularly, C1-C3 represented colon-specific clusters derived from normal tissue, while C4-C6 represented stroma (C4, C5) and mesothelium-derived cells (C6) specific for the ovary. C1-C6 fibroblasts were excluded from CCA, because they have a tissue-specific identity, localization and function that are unlikely to have counterparts in other tissues (see below). All other fibroblasts clustered into 5 clusters shared across cancer types and patients (C7-C11; **Fig. 3c-e** and Supplementary information, **Fig. S3e**). Three other CCA clusters represented a low-quality (C12) or doublet cluster (C13_CD3D, C14_AIF1) (Supplementary information, **S3f,g**). Fibroblasts therefore consist of 11 cellular phenotypes: tissue-specific clusters C1-C6 identified by unaligned clustering and shared clusters C7-C11 identified by CCA (**Fig. 3f,g** for marker genes and functional gene sets).

**Fig. 3.**
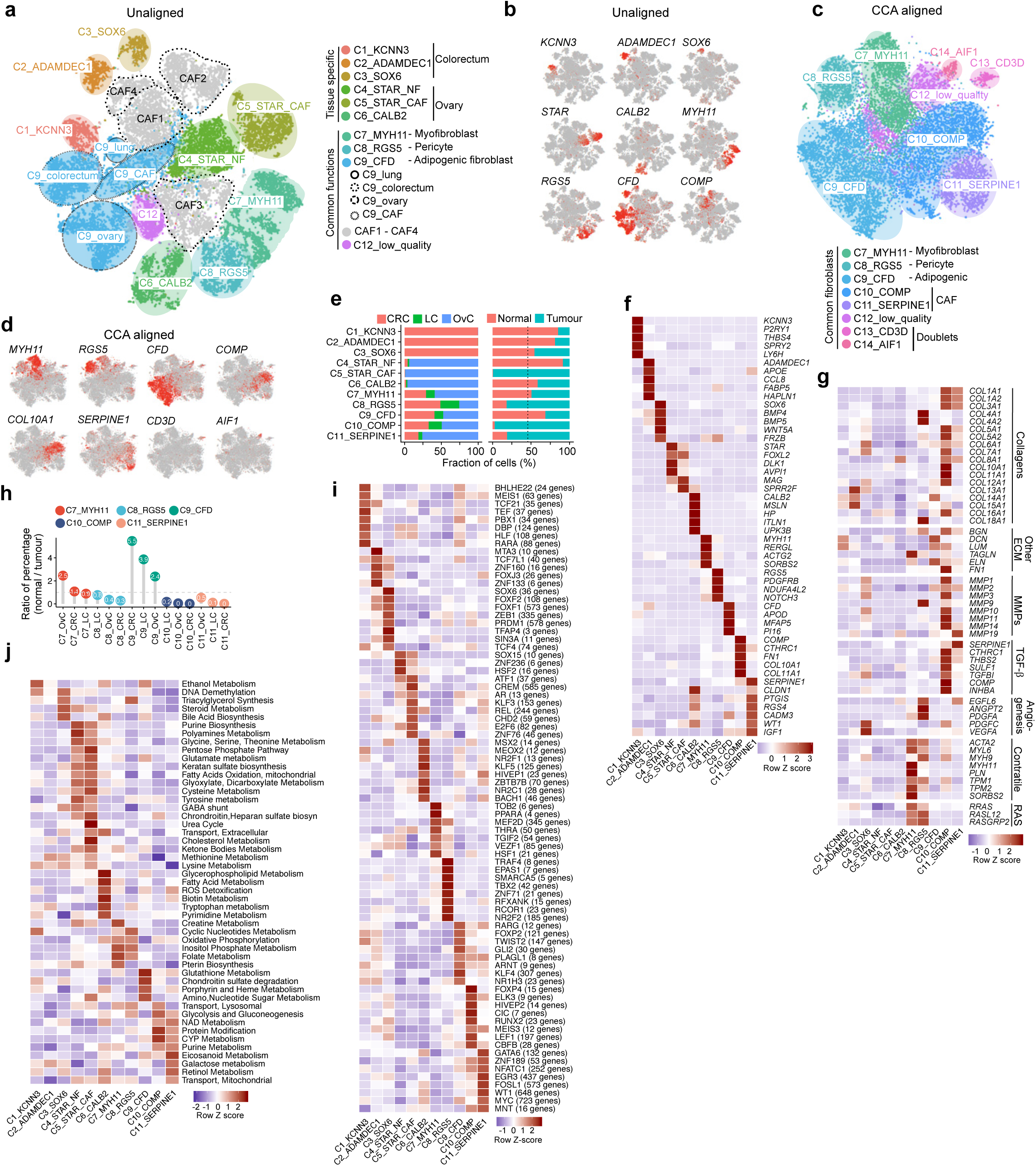
Characterization of 24,622 fibroblasts. **a** t-SNE colour-coded for annotated fibroblasts by unaligned clustering. **b** t-SNEs with marker gene expression in unaligned clusters. **c** t-SNE colour-coded for annotated fibroblasts by CCA. **d** t-SNE with marker gene expression in CCA clusters. **e** Fraction of fibroblast clusters per cancer type (left) and sample origin (right). C7-C11s are shared by CRC, LC and OvC. **f,g** Heatmap of marker gene expression (**f**) and functional gene sets (**g**). **h** Normal/tumour ratio of relative % of fibroblast clusters, <1 indicates tumour enrichment. Pericytes were enriched in tumour (FDR=7.8×10^−10^). **i,j** Heatmap of TF activity (**i**) or metabolic activity (**j**) in fibroblast clusters.

Colon-specific C1-C3s mostly resided in normal tissue (**Fig. 3e**). C1_KCNN3 fibroblasts co-expressed *KCNN3* and *P2RY1* (**Fig. 3f**), a potassium calcium-activated channel (SK3-type) and purine receptor (P2Y1), respectively. Their co-expression defines a novel excitable cell that co-localizes with motor neurons in the gastrointestinal tract and regulates their purinergic inhibitory response to smooth muscle function in the colon^21,22^. C1_KCNN3s also expressed *LY6H*, a neuron-specific regulator of nicotine-induced glutamatergic signalling^23^, suggesting these cells to regulate multiple neuromuscular transmission processes. C2_ADAMDEC1s represented mesenchymal cells of the colon lamina propria^24^, characterised by *ADAMDEC1* and *APOE*. C3_SOX6s were marked by *SOX6* expression, as well as *BMP4, BMP5, WNT5A* and *FRZB* expression (**Fig. 3f**). They are located in close proximity to the epithelial stem cell niche and promote stem cell maintenance in the colon^24^. C4-C5 ovarian stroma cells were marked by *STAR* and *FOXL2*^25,26^, which promote folliculogenesis^27^. Both clusters also expressed *DLK1*, which is typical for embryonic fibroblasts. C4_STARs were derived from normal tissue, while C5_STARs were exclusive to tumour tissue, suggesting that C4_STARs give rise to C5_STARs^25^. Based on calrectinin (*CALB2*) and mesothelin (*MSLN*) expression, C6_CALB2s were likely to represent mesothelium-derived cells^28^. These cells were especially enriched in omentum (Supplementary information, **Fig. S3h**), known to contain numerous mesothelial cells.

C7_MYH11 corresponded to myofibroblasts and were characterised by high expression of smooth muscle-related contractile genes, including *MYH11, PLN* and *ACTG2* (**Fig. 3f**). C8_RGS5 represented pericytes (*RGS5, PDGFRB*), which similar as myofibroblasts expressed contractile genes, but also showed pronounced expression of RAS superfamily members (*RRAS, RASL12*). Additionally, pericytes expressed a distinct subset of collagens (*COL4A1, COL4A2, COL18A1*), genes involved in angiogenesis (*EGFL6, ANGPT2;* **Fig. 3g**) and vessel maturation (*NID1, LAMA4, NOTCH3*; Supplementary information, **Fig. S3i**). Pericytes were enriched in malignant tissue (**Fig. 3e,h** and Supplementary information, **Fig. S3j**). When comparing pericytes from malignant *versus* normal tissue, the former exhibited increased expression of collagens and angiogenic factors (*PDGFA, VEGFA*; Supplementary information, **Fig. S3k**), but reduced expression of the vascular stabilization factor *TIMP3*^29^. These differences may contribute to a leaky tumour vasculature. C9_CFDs expressed adipocyte markers adipsin (*CFD*) and apolipoprotein D (*APOD*), suggesting these are adipogenic fibroblasts. They are positively associated with aging in the dermis^30^, but their role in malignancy has not been established. Notably, in the unaligned clusters, C9s separated into 3 tissue-specific clusters and a single cancer-associated fibroblasts (CAF) cluster (**Fig. 3a**), suggesting that C9 fibroblasts (similar as cECs) lose tissue-specificity in the TME.

C10-C11 represented CAFs showing strong activation of cancer hallmark pathways, including glycolysis, hypoxia, and epithelial-to-mesenchymal transition (Supplementary information, **Fig. S3l**). C10_COMPs typically expressed metalloproteinases (MMPs), TGFß-signalling molecules and extracellular matrix (ECM) genes, including collagens (**Fig. 3g**). They also expressed the TGF-ß co-activator *COMP*, which is activated during chondrocyte differentiation, and activin (*INHBA*), which synergizes with TGF-ß signalling^31,32^. Accordingly, chondrocyte-specific TGF-ß targets (*COL10A1, COL11A1*) were strongly upregulated. C11_SERPINE1s exhibited increased expression of *SERPINE1, IGF1, WT1* and *CLDN1*, which all promote cell migration and/or wound healing via various mechanisms^33–36^. They also expressed collagens, albeit to a lesser extent as C10_COMPs. Additionally, high expression of the pro-angiogenic *EGFL6* suggests these cells to exert paracrine functions^37,38^. Interestingly, the number of C10-C11 CAFs correlated positively with the presence of cancer cells (Supplementary information, **Fig. S3m**), confirming the role of CAFs in promoting tumour growth^20^.

Using SCENIC, we identified TFs unique to each fibroblast cluster (**Fig. 3i**). For instance, MYC and EGR3 underpinned C11_CAFs, while pericytes were characterised by EPAS1, TBX2 and NR2F2 activity. Interestingly, MYC activation of CAFs promote aggressive features of cancers through upregulation of unshielded RNA in exosome^39^. At the metabolic level, we observed that creatine and cyclic nucleotide metabolism, which are essential for smooth muscle function, were upregulated in myofibroblasts (C7), while glycolysis was most prominent in C10-11 CAFs (**Fig. 3j**). Indeed, highly proliferative CAFs rely on aerobic glycolysis, and their glycolytic adaptation promote a reciprocal metabolic symbiosis between CAFs and cancer cells ^20^.

### Dendritic cells, novel markers of cDC maturation revealed

Clustering the transcriptomes of 2,722 DCs identified 5 different DC phenotypes using unaligned and CCA-aligned approaches (**Fig. 4a)**. 92% of cells clustered similarly with both approaches, suggesting DCs in line with their non-resident nature to have limited cancer type specificity. C1_CLEC9As corresponded to conventional DCs type 1 (cDC1; *CLEC9A, XCR1*)^40,41^, C2_CLEC10As to cDCs type 2 (cDC2; *CD1C, CLEC10A, SIRPA*), while C3_CCR7s represented migratory cDCs (*CCR7, CCL17, CCL19*; **Fig. 4b,c** and Supplementary information, **Fig. S4a,b**). Further, C4_LILRA4s represented plasmacytoid DCs (pDCs; *LILRA4, CXCR3, IRF7*), while C5_CD207s were related to cDC2s based on *CD1C* expression. C5_CD207s additionally expressed Langerhans cell-specific markers: *CD207* (langerin) and *CD1A*, but not the epithelial markers *CDH1* and *EPCAM*, typically expressed in Langerhans cells^42^. These cells therefore likely represent a subset of cDC2s with a similar expression as Langerhans cells. Notably, Langerhans-like and migratory DCs were not previously characterised by scRNA-seq, possibly because these studies focused on blood-derived DCs^40^.

**Fig. 4.**
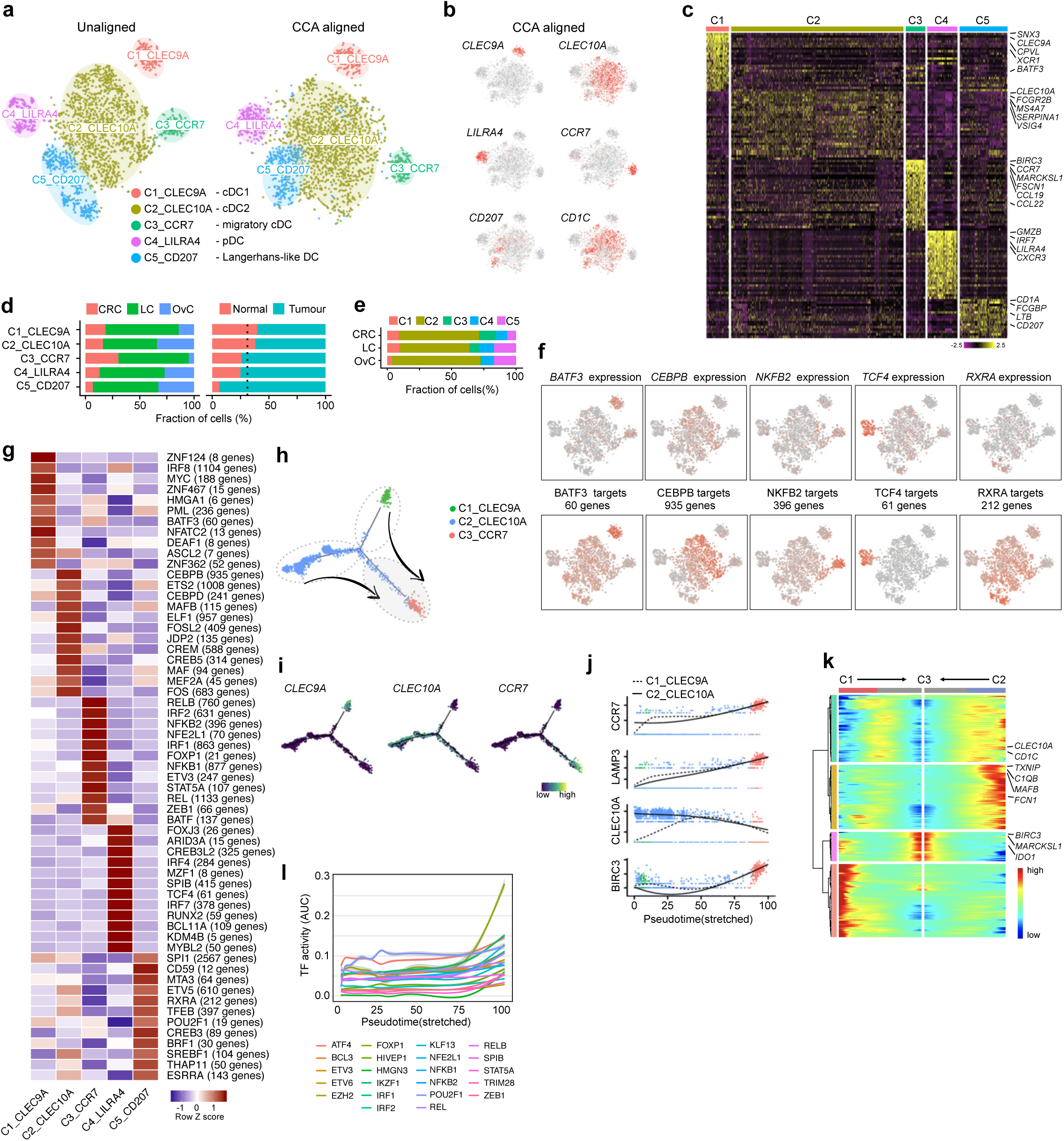
Clustering 2,722 DCs. **a** t-SNEs colour-coded for annotated DCs by unaligned and CCA aligned clustering. **b** t-SNEs with DC marker gene expression in CCA aligned clusters. **c** Heatmap for differential gene expression in unaligned clusters. **d** Fraction of DC clusters per cancer type (left) and sample origin (right). Migratory cDCs were depleted in OvC (FDR=0.017). **e** Fraction of cells in each cancer type per cluster. **f** t-SNEs with gene expression (upper) and corresponding TF activity (lower). **g** Heatmap showing TF activity in CCA aligned clusters. **h** Trajectory inference analysis of cDC-related subclusters. **i** Marker gene expression along the cDC trajectory. **j,k** Marker gene expression (**j**) and expression dynamics (**k**) during cDC maturation. (**l**) TF activation dynamics of cDC2 to migratory cDC differentiation.

Overall, C2_CLEC10As were most abundant, while the number of other DCs varied per cancer type. For instance, C3 was rare in OvC, and C5 enriched in malignant tissue (**Fig. 4d,e** and Supplementary information, **Fig. S4c,d**). SCENIC confirmed known TFs to underlie each DC phenotype, including *BATF3* for cDC1s, *CEBPB* for cDC2s, *NFKB2* for migratory cDCs and *TCF4* for pDCs (**Fig. 4f,g**). We also identified novel TFs (Supplementary information, **Table S8**). For instance, *SPI1*, a master regulator of Langerhans cell differentiation^43^, and *RXRA*, required for cell survival and antigen presentation in Langerhans cells^44^, were both expressed in C5. Cancer hallmark pathway analysis revealed activation of interferon-α and -γ signalling in migratory cDCs, while metabolic pathway analysis confirmed a critical role for folate metabolism (Supplementary information, **Fig. S4e,f**)^45^.

By leveraging trajectory inference analyses (using 3 different pipelines; Methods), we recapitulated the cDC maturation process and observed that cDC2s are enriched in the migrating branch (**Fig. 4h,i**), suggesting that migratory cDCs originated from cDC2s but not cDC1s, at least in tumours. Consistent herewith, some migratory cDC-related genes, i.e. *CCL17* and *CCL22*, were already upregulated in a subset of cDC2s (Supplementary information, **Fig. S4g**), highlighting that cDC2s are in a transitional state. In contrast, cDC maturation markers *CCR7* and *LAMP3* were only upregulated at a later stage of the trajectory (**Fig. 4j**, Supplementary information, **Fig. S4h**)^46^. Interestingly, in OvC, cDC2s got stuck early in the differentiation lineage compared to CRC and LC (Supplementary information, **Fig. S4i**). By modelling expression along the branches, we retrieved 4 clusters with distinct temporal expression (**Fig. 4k**), in which we identified 30 and 210 genes up- or down-regulated (Supplementary information, **Table S9**). For example, *CLEC10A* was gradually lost during cDC2 maturation, while *BIRC3* was upregulated, suggesting they represent novel markers of cDC maturation. Also, when investigating TF dynamics from cDC2s to migratory cDCs, we identified 22 up- and 23 down-regulated TFs, respectively (**Fig. 4l** and Supplementary information, **Fig. S4j**).

### B-cells, comprehensive taxonomy and developmental trajectory

Amongst the 15,247 B-cells, we identified 8 clusters using unaligned clustering (**Fig. 5a**) Three of these represented follicular B-cells (*MS4A1*/CD20), which reside in lymphoid follicles of intra-tumour tertiary lymphoid structures, while 4 clusters were antibody-secreting plasma cells (*MZB1* and *SDC1*/CD138) (Supplementary information, **Fig. S5a-b**). We also retrieved a T-cell (C9_CD3D) doublet cluster (Supplementary information, **Fig. S5c**). CCA identified 2 additional clusters: one unaligned follicular B-cell cluster, which was split into 2 separate clusters (C2 and C3, **Fig. 5a-b**) and one additional cancer cell (C10_KRT8) doublet cluster (Supplementary information, **S5c**). Overall, this resulted in 8 relevant B-cell clusters, each of them characterised by functional gene sets (**Fig. 5c**).

**Fig. 5.**
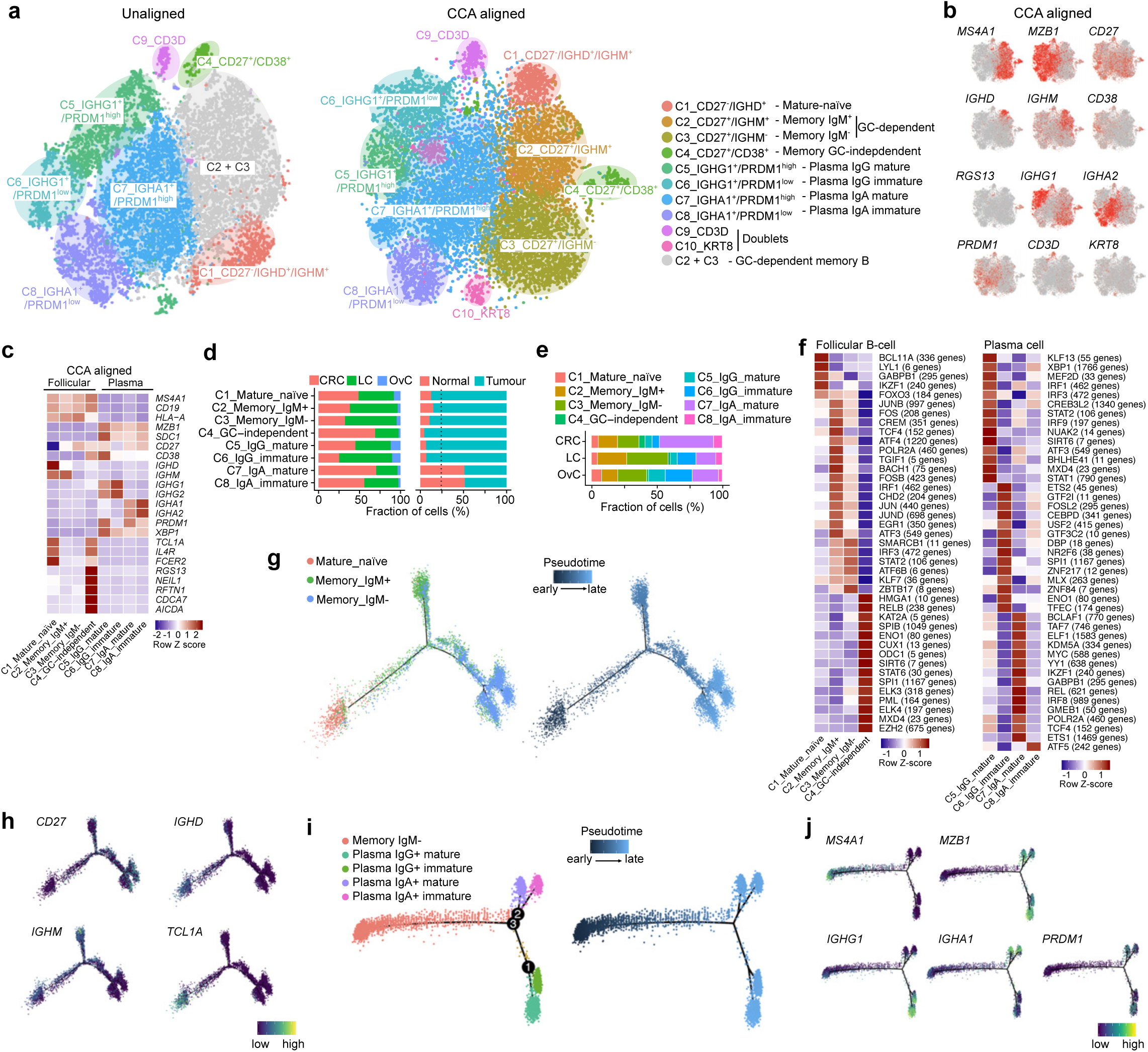
B-cell taxonomy and developmental trajectory. **a** t-SNEs colour-coded for annotated B-cells using unaligned and CCA aligned clustering. **b** t-SNEs with marker gene expression in CCA clusters. **c** Heatmap of functional gene sets in CCA clusters. **d** Fraction of B-cell clusters per cancer type (left) and sample origin (right). **e** Fraction of cells in each cancer type per cluster. **f** Heatmap with TF activity by SCENIC, for follicular B-cell (left) or plasma cell clusters (right). **g** Developmental trajectory for GC-dependent memory B-cells, colour-coded by cell type (left) and pseudotime (right). **h** Marker gene expression of the GC-memory B-cell trajectory as in (**g**). **i** Trajectory of IgM^-^ memory B to IgG^+^ or IgA^+^ plasma cells, colour-coded by branch type (left) and pseudotime (right). **j** Marker gene expression dynamics during plasma cell differentiation as in (**i**).

Follicular B-cells were composed of mature-naïve (*CD27*^-^, C1) and memory (*CD27*^+^, C2-4) B-cells (**Fig. 5c**). The former are characterised by a unique *CD27*^-^ /*IGHD*^+^(IgD)/*IGHM*^+^(IgM) signature and give rise to the latter by migrating through the germinal centre (GC; referred to as GC-memory B-cells). This process requires expression of migratory factors *CCR7* (for GC entry) and *GPR183* (for GC exit; Supplementary information, **Fig. S5d**)^47^. In the GC, *IGHM* undergoes class-switch recombination to form other immunoglobulin isotypes. Indeed, GC-memory B-cells separated into *IGHM*^+^ and *IGHM*^-^ populations, i.e., C2 *IGHM*^*+*^ and C3 *IGHM*^*-*^ clusters (**Fig. 5a-c**). A rare population of memory B-cells is generated independently of the GC^48^. These GC-independent memory B-cells corresponded to C4_CD27^+^/CD38^+^s, lacking GC migratory factors *GPR183* and *CCR7*, but expressing the anti-GC migration factor *RGS13*, which may form the basis for their GC exclusion (**Fig. 5b** and Supplementary information, **Fig. S5d**)^49^. Although little is known about GC-independent B-cells, they appear early during immune response and respond to a broader range of antigens with less specificity as GC-memory B-cells^50^. Interestingly, C4s exhibited an expression signature intermediate to mature-naïve and GC-memory B-cells (Supplementary information, **Fig. S5e**). Expression of *IGHD* and *IGHM* was low, while *IGHG1* and *IGHG3* were elevated (Supplementary information, **Fig. S5f**), suggesting C4s to have completed class-switch recombination. Indeed, *AICDA* expression, which induces mutations in class-switch regions during recombination^50^, was elevated in C4s (**Fig. 5c**). They were also characterised by several uniquely expressed genes and enriched for proliferative cells (Supplementary information, **Fig. S5g** and **Table S5**). Next to follicular B-cells, we identified 4 clusters of plasma B-cells (C5-C8), which can be separated based on expression of immunoglobulin heavy chains, i.e. *IGHG1* (IgG) *versus IGHA1* (IgA). Both could be further stratified based on their antibody-secreting capacity as determined by *PRDM1* (Blimp-1)^50^: low *versus* high for immature *versus* mature plasma cells, overall resulting in 4 plasma B-cell clusters (**Fig. 5c**).

Importantly, B-cell clusters were enriched in all tumours, except for IgA-expressing plasma cells, which mainly resided in mucosa-rich normal colon (**Fig. 5d,e** and Supplementary information, **Fig. S5h-j**). Additionally, GC-independent memory B-cells were most prevalent in CRC. B-cells were also enriched in border *versus* core fractions of LC tumours (Supplementary information, **Fig. S5k**). Using SCENIC, each B-cell cluster was characterised by a unique set of TFs (**Fig. 5f**). For instance, GC-independent memory B-cells upregulated NF-κB (RELB) and STAT6, which is known to suppress *GPR183*^51^. Some TFs were upregulated in mature (*PRDM1*^high^) plasma cells, irrespective of their heavy chain expression. These included multiple immediate-early response TFs (FOS, JUND and EGR1) and the interferon regulatory factor IRF1 (Supplementary information, **Fig. S5l**), suggesting they are involved in plasma cell maturation. C5_IgG_mature B-cells, relative to all other plasma B-cells, exhibited strong activation of nearly all cancer hallmark pathways, indicating an active role of C5s in the TME (Supplementary information, **Fig. S5m**).

Trajectory inference analysis confirmed that mature-naïve B-cells differentiate into either GC-memory IgM^+^ or IgM^-^ branches. As expected, IgM^+^ but not IgM^-^ cells were located halfway the trajectory (**Fig. 5g** and Supplementary information, **Fig. S5n**), confirming IgM^+^ cells to undergo class-switch recombination into IgM^-^ cells. Memory B-cells of the IgM^+^ and IgM^-^ lineages were similarly distributed in OvC and CRC, but in LC they were more differentiated (Supplementary information, **Fig. S5o**). By overlaying gene expression dynamics on the trajectory, we identified several genes up- or down-regulated along the pseudotime, including *CD27* and *TCL1A*, respectively (**Fig. 5h**; Supplementary information, **Fig. S5p,q** and **Table S9**). In line with *CCR7* and *GPR183* determining GC entry and exit, *CCR7* was expressed in mature-naïve B-cells (C1, before entry) but disappeared in *IGHM*^-^ B-cells. *Vice versa, GPR183* was only expressed after GC entry (C2 and C3, Supplementary information, **Fig. S5d,q**). Similarly, we assessed the trajectory of class-switched GC-memory B-cells (C3) differentiating into plasma cells. We confirmed that GC-memory B-cells differentiate into either IgG^+^- or IgA^+^-expressing plasma cells (**Fig. 5i**) and that both branches subsequently dichotomize into mature or immature states based on *PRDM1* expression (**Fig. 5j**). Cells were similarly distributed along the trajectory regardless of the cancer type, although in LC there was an enrichment towards the beginning of the IgA lineage (Supplementary information, **Fig. S5r**). Further, when assessing underlying expression dynamics along the trajectory, we identified several genes staging the differentiation process (Supplementary information, **Fig. S5s** and **Table S9**). For example, we found *TNFRSF17* (also known as B-cell maturation antigen) to increase along the IgA^+^ plasma cell trajectory^52^.

### T-/NK-cells show cancer type-dependent prevalence

Altogether, 52,494 T- and natural killer (NK) cells clustered into 12 and 11 clusters using unaligned and CCA-aligned methods (**Fig. 6a,b**). The additional cluster identified by unaligned clustering (C12) was composed of cells from normal lung tissue (Supplementary information, **Fig. S6a,b**). CCA did not affect clustering of T-/NK-cells in tumours, indicating that T-cells have limited cancer type-specific differences. Besides C12 and a low-quality cluster (C11, Supplementary information, **Fig. S6c,d**), T-/NK-cells consisted of 10 phenotypes, including 4 CD8^+^ T-cell (C1-C4), 4 CD4^+^ T-cell (C5-C8) and 2 NK-cell clusters (C9-C10).

**Fig. 6.**
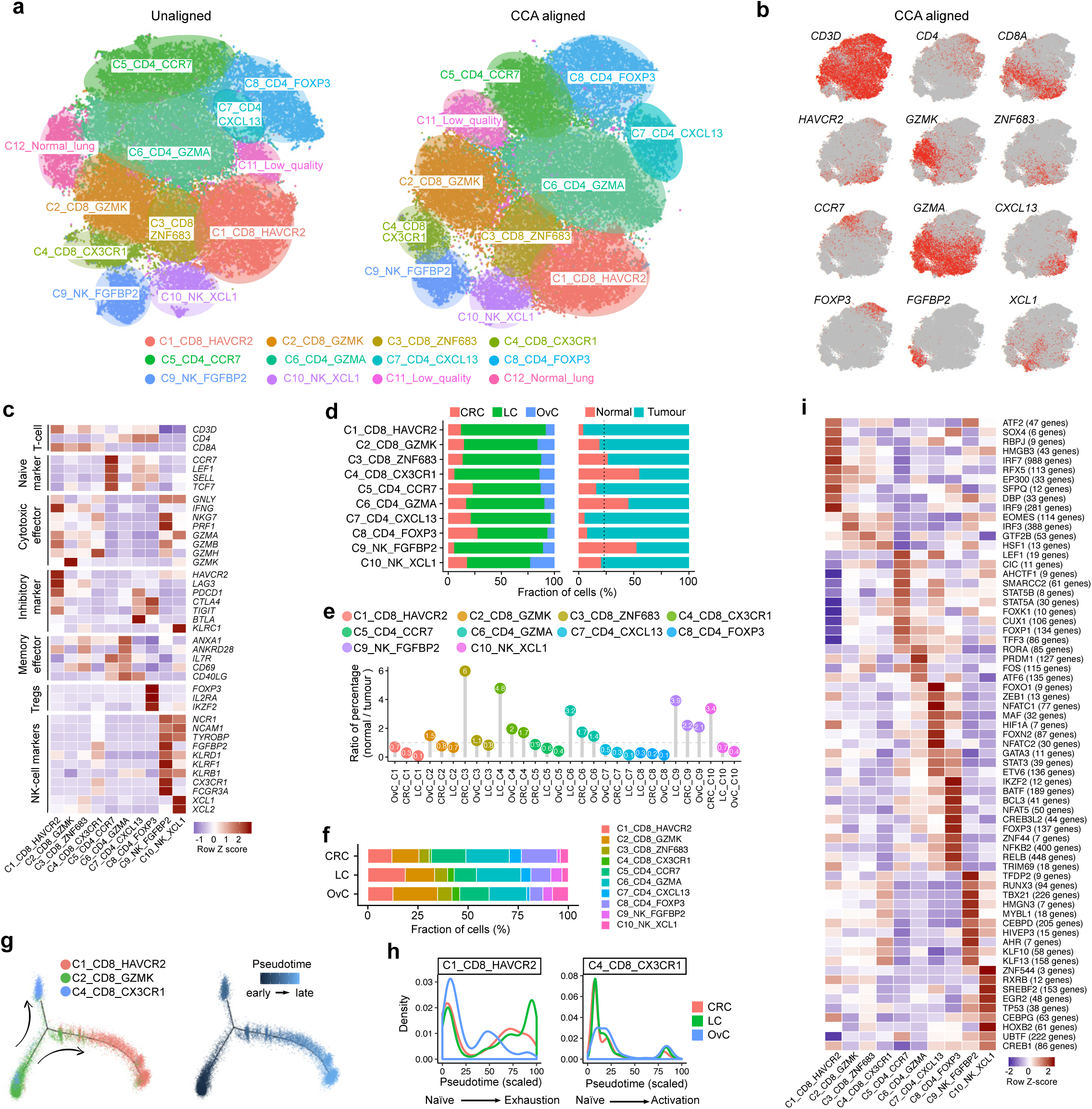
Profiling 52,494 T-/NK-cells. **a** t-SNEs colour-coded for annotated T-/NK-cell using unaligned and CCA aligned clustering. **b** t-SNEs with marker gene expression in CCA clusters. **c** Heatmap of functional gene sets in CCA clusters. **d** Fraction of cells for T-/NK-cell clusters per cancer type (left) and sample origin (right). **e** Normal/tumour ratio of relative % of T-/NK-cell clusters, <1 indicates tumour enrichment. C1, C2, C5, C7, C8 were enriched in tumour (FDR<5.1×10^−25^), C9 was enriched in normal (FDR=1.5×10^−219^). **f** Fraction of T-/NK-cells in each cancer type per cluster. C4 and C8 were rare in CRC (FDR=0.019) and OvC (FDR=0.034), respectively. **g** Heatmap with TF activity of T-/NK-cell clusters by SCENIC. **h** Differentiation trajectory for CD8+ T cell lineages, colour-coded by cell type (left) and pseudotime (right). **i** Density plots for CRC, LC and OvC along the two CD8+ T-cell trajectories.

The C1_CD8_HAVCR2 cluster consisted of exhausted CD8^+^ cytotoxic T-cells characterised by cytotoxic effectors (*GZMB, GNLY, IFNG*) and inhibitory markers (*HAVCR2, PDCD1, CTLA4, LAG3, TIGIT*; **Fig. 6c**). C2_CD8_GZMKs represented pre-effector cells as expression of *GZMK* was high, but expression of cytotoxic effectors low. C3_CD8_ZNF683s constituted memory CD8^+^ T-cells based on *ZNF683* expression^53^, while C4_CD8_CX3CR1s corresponded to effector T-cells due to high cytotoxic marker expression. Remarkably, C4s also expressed markers typically observed in NK-cells (*KLRD1, FGFBP2, CX3CR1*), suggesting they are endowed with NK T-cell (NKT) activity. Similarly, based on marker gene expression, we assigned C5_CD4_CCR7s to naïve (*CCR7, SELL, LEF1*), C6_CD4_GZMAs to CD4^+^ memory/effector (*GZMA, ANXA1*) and C7_CD4_CXCL13s to exhausted CD4^+^ effector T-cells (*CXCL13, PDCD1, CTLA4, BTLA*). Based on *FOXP3* expression C8_FOXP3s were assigned CD4^+^ regulatory T-cells (Tregs). Finally, two clusters contained NK-cells based on NK- (*NCR1, NCAM1*) but not T-cell (*CD3D, CD4, CD8A*; **Fig. 6b,c**) marker gene expression. Particularly, C9_NK_FGFBP2s represented cytotoxic NK-cells due to expression of *FGFBP2, FCGR3A* and cytotoxic genes including *GZMB, NKG7* and *PRF1*, while C10_NK_XCL1s appeared to be less cytotoxic, but positive for *XCL1* and *XCL2*, two chemo-attractants involved in DC recruitment enhancing immunosurveillance^54^.

Interestingly, T-cell clusters were highly similar to the T-cell taxonomy derived from breast, liver and lung cancer, despite underlying differences in sample preparation and single-cell technology (Supplementary information, **Fig. S6e**)^53,55,56^. Indeed, C8 cells could be re-clustered into CLTA4^high^ and CLTA4^low^ clusters with corresponding marker genes (Supplementary information, **Fig. S6f-g**), as reported^53,56^, while also both NK clusters corresponded to recently identified NK subclusters shared across organs and species^57^.

Several T-cell phenotypes, especially those with inhibitory markers, were enriched in tumour tissue (**Fig. 6d,e**, Supplementary information, **Fig. S6h**). C9_NK_FGFBP2s were more prevalent in normal tissue, suggesting these to represent tissue-patrolling phenotypes of NK-cells. All T-cell clusters were more frequent in LC, while cytotoxic T-cells were rare in CRC and regulatory T-cells underrepresented in OvC (**Fig. 6f**). Expression of inhibitory markers (*HAVCR2, LAG3, PDCD1*) was enhanced in exhausted/cytotoxic C1_CD8_HAVCR2s residing in tumour *versus* normal tissue (Supplementary information, **Fig. S6i**). We also observed expression of KLRC1 (*NKG2A*), a novel checkpoint^58,59^ exclusively in C10 NK-cells (**Fig. 6c**). CD8^+^ T-cell trajectory analysis revealed that C2 pre-effector T-cells also contained naïve CD8^+^ T-cells, which expressed *CCR7, TFC7* and *SELL*, and formed the root of the trajectory (Supplementary information, **Fig. S6j,k**). Pre-effector T-cells then differentiated into either exhausted (C1_CD8_HAVCR2) or effector (C4_CD8_CX3CR1) T-cells (**Fig. 6g**). Dynamic expression of marker genes along both trajectories confirmed high expression of *IFNG*, inhibitory and cytotoxicity markers in the HAVCR2 trajectory (Supplementary information, **Fig. S6l**). Interestingly, LC CD8^+^ T-cells were more differentiated in this trajectory and thus more exhausted compared to T-cells from CRC and OvC (**Fig. 6h**).

TFs underlying each T-/NK-cell phenotype were identified by SCENIC (**Fig. 6i**): for instance, FOXP3 was specific for C8s, as expected, while IRF9, which induces *PDCD1*^60^, was increased in exhausted CD8^+^ T-cells (C1). C1_CD8_HAVCR2 T-cells exhibited high interferon activation based on cancer hallmark analysis (Supplementary information, **Fig. S6m**), while metabolic pathway analysis revealed upregulation of glycolysis and nucleotide metabolism in T-cell phenotypes enriched in tumours (C1, C7-C8; Supplementary information, **Fig. S6n**). Finally, we noticed a negative correlation between the prevalence of cancer and immune cells, including several T-cell phenotypes (Supplementary information, **Fig. S3m**). When scoring cancer cells for cancer hallmark pathways and comparing these scores with stromal cell phenotype abundance, some remarkable associations were noticed. Specifically, C1_CD8_HAVCR2 T-cells were positively correlated with augmented interferon signalling, inflammation and IL6/JAK/STAT3 signalling in cancer cells (Supplementary information, **Fig. S6o**).

### Trajectory of monocyte-to-macrophage differentiation revealed

In the 32,721 myeloid cells, we identified 12 unaligned clusters, including 2 monocyte (C1-C2), 7 macrophage (C3-C9) and 1 neutrophil (C10) clusters (**Fig. 7a,b**). A low-quality cluster (C11) and myeloid/T-cell doublet cluster (C12_CD3D) are not discussed (Supplementary information, **S7a,b**). Only C8 macrophages were tissue-specific, while remaining cells clustered similarly with CCA as with unaligned clustering, expressing the same marker genes and functional gene sets (**Fig. 7c**, Supplementary information, **Fig. S7c,d**).

**Fig. 7.**
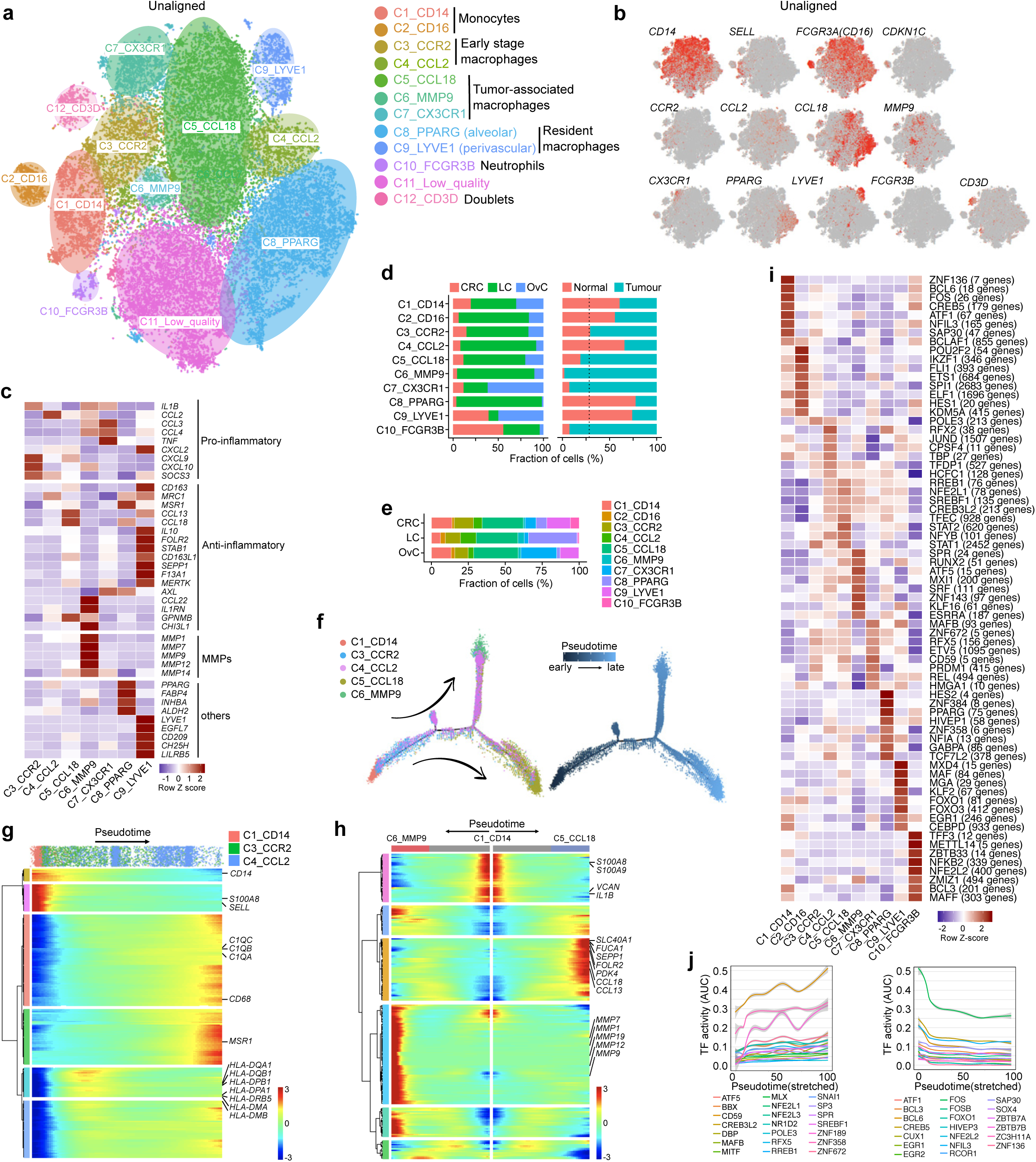
Profiling of monocytes, macrophages and neutrophils. **a** t-SNE colour-coded for annotated myeloid cell using unaligned clustering. **b** t-SNEs with marker gene expression in myeloid clusters. **c** Heatmap of functional gene sets in myeloid clusters. **d** Fraction of myeloid clusters per cancer type (left) and sample origin (right). C9 was enriched in normal (FDR=3.0×10^−31^) and C8 in normal lung (FDR ≈0) tissue. C5-C7 and C10 (FDR < 3.3×10^−31^) were enriched in tumour. **e** Fraction of cells in each cancer type per cluster. **f** Monocyte-to-macrophage differentiation trajectory, colour-coded by cluster (left) or pseudotime (right). **g,h** Gene expression dynamics during differentiation of C1 monocytes to C4 macrophages (**g**), or terminal differentiation of C5/C7 macrophages (**h**). **i** Heatmap showing TF activity by SCENIC. **j** TF activation (left) or inactivation (right) during monocyte-to-macrophage differentiation, before branching into terminal differentiation.

Monocytes clustered separately from macrophages based on reduced macrophage marker expression (*CD68, MSR1, MRC1*) and a phylogenetic reconstruction (Supplementary information, **Fig. S7e,f**). C1_CD14 monocytes represented classical monocytes based on high *CD14* and *S100A8*/*9* expression and typically are recruited during inflammation. They expressed the monocyte trafficking factors *SELL* (CD62L) *-*involved in EC adhesion*-* and *CCR2*, a receptor for the pro-migratory cytokine CCL2. C2_CD16s were less abundant and represented non-classical monocytes based on low *CD14*, but high expression of *FCGR3A* (CD16) and other marker genes (*CDKN1C, MTSS1*; Supplementary information, **Fig. S7f**)^61^. C2s constantly patrol the vasculature, express *CX3CR1* (Supplementary information, **Fig. S7d,g**) and migrate into tissues in response to CX3CL1 derived from inflamed ECs.

Macrophages are classified based on origin (tissue-resident *versus* recruited) or their pro-*versus* anti-inflammatory role (M1-like *versus* M2-like, **Fig. 7c**). C3_CCR2s and C4_CCL2s represented early-stage macrophages that were closely-related, not enriched in tumours (**Fig. 7d** and Supplementary information, **Fig. S7e**) and become replenished by classical monocytes. Specifically, C3 macrophages represented immature macrophages closely related to C1 monocytes, as they also express *CCR2* (**Fig. 7b**). They were characterised by pronounced M1 marker gene expression (*IL1B, CXCL9, CXCL10, SOCS3*; **Fig. 7c**). C4_CCL2s were characterised by *CCL2* expression, which is another M1 marker promoting immune cell recruitment to inflammatory sites. Compared to C3s, C4 macrophages expressed less *CCR2*, but moderate levels of the M2 marker gene *MRC1*, suggesting an intermediate pro-inflammatory phenotype.

Macrophages belonging to C5-C7 clusters were enriched in malignant tissue and represented tumour-associated macrophages (TAMs, **Fig. 7d**, Supplementary information, **Fig. S7h**). C5_CCL18s represented ∼72% of all TAMs and were characterised by M2 marker expression, including *CCL18* and *GPNMB* (**Fig. 7c)**. Additional heterogeneity separated C5 cells into intermediate and more differentiated M2 macrophages, although differences were graded, consistent with a continuous phenotypic spectrum (Supplementary information, **Fig. S7i**). Indeed, there was more pronounced M2 marker expression (e.g. *SEPP1, STAB1, CCL13*) in 34% of C5s^62^. These also expressed key metabolic pathway regulators, i.e. *SLC40A1* (iron), *FOLR2* (folate), *FUCA1* (fucose) and *PDK4* (pyruvate), linking M2 differentiation with metabolic reprogramming. C6_MMP9 macrophages expressed a unique subset of M2 markers (*CCL22, IL1RN, CHI3L1*) and several MMPs, suggesting a role in tumour tissue remodelling. Cancer hallmark analysis revealed enrichment in EMT, hypoxia, glycolysis and many other pathways (Supplementary information, **Fig. S7j**). C7_CX3CR1 macrophages expressed genes involved both in M1 and M2 polarization (*CCL3, CCL4, TNF, AXL*, respectively, **Fig. 7c**). Interestingly, AXL is involved in apoptotic cell clearance^63^, whereas other M2 markers involved in pathogen clearance, i.e. *MRC1* and *CD163*, were absent, suggesting a unique phagocytic pattern of C7 cells. They are also correlated with poor prognosis in OvC and CRC^64,65^. Of note, C7 macrophages shared their CD16^high^/CX3CR1^high^ phenotype with C2 non-classical monocytes, suggesting both clusters may be related (Supplementary information, **Fig. S7g**). C8_PPARG macrophages corresponded to resident alveolar macrophages due to expression of the resident alveolar macrophage marker *PPARG*. They were exclusive to normal lung tissue (**Fig. 7d**), expressed established M2 markers (*MSR1, CCL18, AXL*) ^62,66^ in addition to anti-inflammatory genes (*FABP4, ALDH2*) ^67,68^. C9_LYVE1 macrophages also represented resident macrophages with pronounced M2 marker expression and enrichment in normal tissue. They often locate at the perivasculature of different tissues where they contribute to both angiogenesis and vasculature integrity^69–71^. Indeed, C9 macrophages expressed the angiogenic factor *EGFL7*, but also immunomodulators *CD209, CH25H* and *LILRB5*, which are implicated in both innate and adaptive immunity^62,72,73^.

Finally, the C10_FCGR3B cluster represented neutrophils expressing the neutrophil-specific antigen CD16B (encoded by *FCGR3B*), but not *MPO*, which is typically expressed in neutrophils during inflammation and microbial infection. C10 cells expressed pro-inflammatory factors (*CXCL8, IL1B, CCL3, CCL4*; Supplementary information, **Fig. S7g**) and, in line with their pro-tumour activity, also pro-angiogenic factors (*VEGFA, PROK2*)^74^. Notably, neutrophils were strongly enriched in malignant tissue, but were characterised by low transcriptional activity (689 detected genes/cell; **Fig. 7d**, Supplementary information, **Fig. S7b**).

Interestingly, except for resident alveolar macrophages (C8), all myeloid clusters were present in each cancer type, albeit with some preferences (**Fig. 7d,e)**. Notably, similar to other scRNA-seq studies^4,6,7,75^, we also failed to identify myeloid-derived suppressor cells (MDSCs). To delineate monocyte-to-macrophage differentiation, we performed a trajectory inference analysis. We excluded non-classical monocytes and related macrophages (C2, C7), and resident macrophages (C8, C9). In the trajectory, C1 monocytes were progenitor cells for C3 immature macrophages (**Fig. 7f**). Next on the time scale were C4 macrophages, which further separated into C5 and C6 macrophages, suggesting C4 macrophages to be endowed with high plasticity prior to M2 differentiation. Interestingly, LC macrophages were more differentiated in both lineages (Supplementary information, **Fig. S7k**). Profiling of gene expression dynamics along the trajectory (**Fig. 7g,h**) revealed a reduction of known monocyte markers (*CD14, S100A8, SELL*) and increased expression of 230 other genes (Supplementary information, **Table S9**), including several M2 markers. SCENIC identified several TFs underlying each myeloid phenotype or the monocyte-to-macrophage differentiation trajectory (**Fig. 7i,j** and Supplementary information, **S7l,m**). For example, there was a gradual increase of MAFB and decrease of FOS, FOSB and EGR1 along the trajectory, as reported^76,77^. Interestingly, terminally differentiated clusters (C5, C6) were characterised by distinct TFs, but also shared TFs, including the hypoxia-induced HIF-2α (EPAS1; Supplementary information, **Fig. S7n**)^78^.

Finally, we also identified 1,962 mast cells. These cells represent a rare stromal cell type that was not enriched for in tumours, and that could be subclustered into 4 cellular phenotypes (Supplementary information, **Fig. S8a-h**).

### Mapping the blueprint in breast cancer

In 3 different cancers, we identified 68 stromal cell (sub)types, of which 46 were shared. To confirm this heterogeneity in another cancer type, we profiled 14 treatment-naïve breast cancers (BC) using 5’-scRNA-seq and clustered the 44,024 cells with high quality data (Methods). After assigning cell types (**Fig. 8a**, Supplementary information, **Fig. S9a**), we re-clustered cells per cell type using unaligned clustering, or after pooling cell type data from BC with those from other cancer types, while applying CCA alignment for 5’ *versus* 3’-scRNA-seq. Both approaches clustered the 14,413 T-cells from BC into their 10 cellular phenotypes, each with similar expression signatures as described for 3’-scRNA-seq (**Fig. 8b** and Supplementary information, **Fig. S9b**). However, in other cell types unaligned clustering failed to identify the cellular phenotypes, especially when they were less abundant. In contrast, CCA recovered 43 out of the 46 shared phenotypes (**Fig. 8b,c**, Supplementary information, **Fig. S9c**). Only for mast cells, for which too few cells were detected (n=360), CCA also failed to identify the respective phenotypes. Notably, across cancer types all cellular phenotypes were characterised by a highly similar expression of marker genes and underlying TFs (**Fig. 8d,e** and Supplementary information, **Fig. S9d-h**). These data confirm that the stromal cell blueprint can also be assigned to other cancer types.

**Fig. 8.**
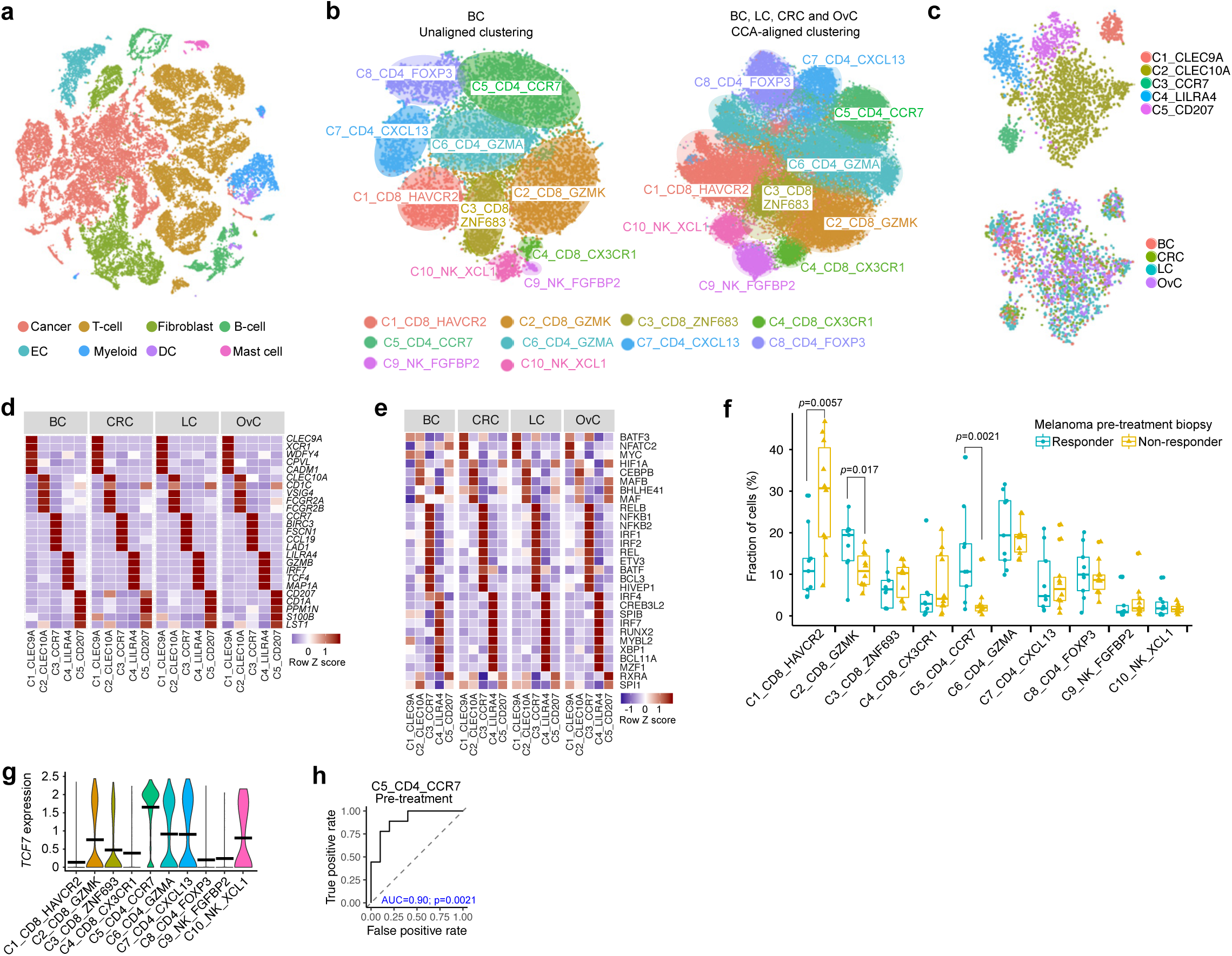
Validation of the stromal blueprint. **a** t-SNE of BC cells colour-coded for cell types. **b** t-SNEs of T-/NK-cells by unaligned clustering or CCA-aligned clustering with 3’-scRNA-seq data. **c** t-SNEs of CCA-aligned clusters colour-coded for annotated DCs (upper) and cancer type (lower). **d** Heatmap of marker gene expression across DC clusters in different cancer types. **e** TF activity across DC subclusters in different cancer types. **f** Fraction of T-/NK-cell clusters in pre-treatment biopsies from melanoma patients treated with ICI. **g** Violin plot showing *TCF7* expression in T-/NK-cell clusters from pre-treatment melanoma patients. **h** Receiver operating characteristic (ROC) analysis to evaluate the predictive effect of naïve CD4^+^ T-cells on response to checkpoint immunotherapy. The area under the ROC curve (AUC) was used to quantify response prediction.

When subsequently comparing stromal cell type distribution between BC and all other cancers, we found more T-cells in BC than CRC or OvC, but not LC (Supplementary information, **Fig. S9i**). At the subcluster level, BC was enriched for pDCs (C4_LILRA4), but had few lymphatic ECs (C5_PROX1; Supplementary information, **Fig. S9j**). Possibly, this is because most patients (8/14) had a triple-negative BC, which is more immunogenic, without lymph node involvement.

### The blueprint as a guide to interpret scRNA-seq studies

We also applied our blueprint to SMART-seq2 data from melanomas treated with immune checkpoint inhibitors (ICIs). We clustered our T-/NK-cells from the blueprint with the 12,681 T-/NK-cells profiled by SMART-seq2^8^, while performing CCA for technology. This resulted in the 10 T-/NK-cell phenotypes of the blueprint (Supplementary information, **Fig. S10a-c**). Cells profiled by both technologies contributed to every phenotypic T-/NK-cell cluster, each with similar expression signatures, suggesting effective CCA alignment. Next, we confirmed findings by Sade-Feldman et al.^8^, showing that *i)* presence of exhausted CD8^+^ T-cells (C1) in melanoma tumours predicts resistance to ICI, while *ii)* increased expression of the naïve T-cell marker *TCF7* across CD8^+^ T-cells predicts response to ICI (Supplementary information, **Fig. S10d**). However, when assessing *TCF7* in the context of the blueprint, we found it was expressed in 2 out of 4 CD8^+^ T-cell phenotypes (C2-C3), of which only pre-effector CD8^+^ T-cells (C2) were significantly more prevalent in responders (**Fig. 8f,g**). Additionally, *TCF7* expression was high in naïve CD4^+^ T-cells (C5), which were also enriched in responders (*p*=0.0021). Receiver operating characteristic (ROC) analysis to evaluate the predictive effect of the C5 cluster revealed an AUC of 0.90 (*p*=0.0021; **Fig. 8h**). Albeit to a lesser extent, C1 and C2 clusters were also enriched in non-responders and responders, respectively (Supplementary information, **Fig. S10e**). Notably, CD4^+^ TCF7^+^ T-cells resided outside of blood vessels, within the tumour at the peritumoral front (Supplementary information, **Fig. S10f**).

Next, we applied our blueprint to monitor changes in T-/NK-cells during ICI. When comparing pre-*versus* on-treatment biopsies (n=4 with response *versus* n=6 without response), we observed an increase in exhausted CD8^+^ T-cells (C1_CD8_HAVCR2) in on-treatment biopsies. *Vice versa*, there was a relative decrease in naïve CD4^+^ (C5_CD4_CCR7) T-cells (Supplementary information, **Fig. S10g,h**). Notably, these differences were only observed in responding patients, suggesting that during response, phenotypic clusters that predict resistance in the pre-treatment biopsy increase, while those predicting response decrease in prevalence. Overall, these data illustrate that single-cell data obtained with various technologies can be re-analysed in the context of the blueprint.

### Validation of the blueprint at protein level

With the availability of CITE-seq, we can now simultaneously detect RNA and protein expression at single-cell level^79^. To confirm the cancer blueprint at protein level, a panel of 198 antibodies (Supplementary information, **Table S10**) compatible with 3’-scRNA-seq was used. We processed 5 BCs, obtaining 6,194 cells with both transcriptome and proteome data. Independent clustering of both datasets revealed how cell types could be discerned based on either marker gene or protein expression (**Fig. 9a,b**). Since antibodies were mainly directed against immune cells, especially T-cells, we focused our subclustering efforts on this cell type. We pooled 1,310 T-/NK-cells with both RNA and protein data together with T-/NK-cells from the blueprint. Subsequent clustering based on scRNA-seq data accurately assigned each T-/NK-cell to its phenotypic cluster (**Fig.9c,d**). Next, we selected marker genes amongst the 198 antibodies and explored protein expression per cluster (**Fig. 9e**). A combination of CD3, CD4, CD8 and NCR1 effectively discriminated CD4^+^, CD8^+^ T-cells and NK-cells. The T-cell exhaustion marker PD-1 discriminated exhausted CD4^+^ and CD8^+^ T-cell phenotypes (C1, C7), while IL2RA (CD25) was specific for CD4^+^ Tregs (C8). CD8^+^ memory T-cells (C3) were characterised by high ITGA1 but low PDCD1. Both the cytotoxic T-/NK-cells (C4, C9) had high levels of KLRG1, while CD4^+^ naïve cells had high ITGA6 and SELL (C5). Unfortunately, there were no antibodies specific for C2 and C6 cells. Despite this limitation, a random forest model developed to predict major cell types and T-cell phenotypes based on CITE-seq classified >80% of cells into the same cell (sub)type compared to scRNA-seq data.

**Fig. 9.**
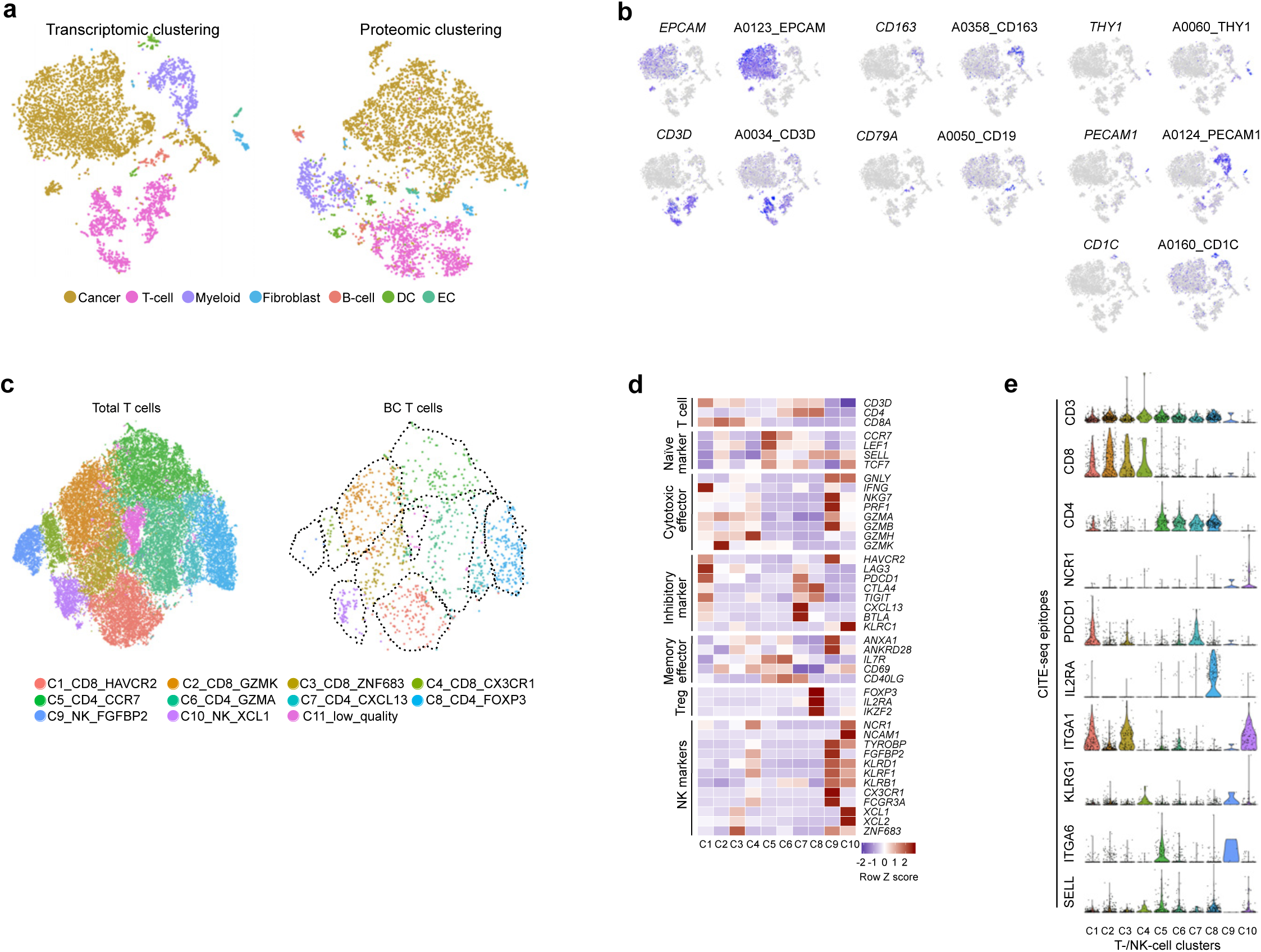
Validation of the stromal blueprint by CITE-seq. **a** t-SNEs of CITE-seq profiled BC cells clustered into cell types based on RNA (left) or protein (right) data. **b** Marker gene or protein expression for each cell type. **c** t-SNE plots showing BC T-/NK-cells co-clustered with 3’-scRNA-seq data from other cancer types (left), while highlighting only T-/NK-cells with BC origin (right). **d** Heatmap with marker gene expression of T-/NK-cell clusters. **e** Expression by CITE-seq markers per T-/NK-cell cluster.

## Discussion

Here, we performed scRNA-seq on 233,591 single cells from 36 patients with either lung, colon, ovarian or breast cancer. By applying two different clustering approaches *-*one designed to detect tissue-specific differences, the other to find shared heterogeneity amongst stromal cell types*-* we constructed a pan-cancer blueprint of stromal cell heterogeneity. Briefly, we found that tissue-resident cell types, including ECs and fibroblasts, were characterised by considerable patient and tissue specificity in the normal tissue, but that part of this heterogeneity disappeared within the TME. On the other hand, phenotypes involving non-residential cell types, which encompass most of the tumour-infiltrating immune cells, were often shared amongst all patients and cancer types. Overall, we identified 68 stromal phenotypes, of which 46 were shared between cancer types and 22 were cancer type-unique. Amongst the shared phenotypes, several have not previously been described at single-cell level, including tumour-associated pericytes and other fibroblast phenotypes, mast cells, GC-independent B-cells, neutrophils, *etc.* Of note, by applying a CITE-seq approach to simultaneously profile gene and protein expression, we confirmed all major cell types and T-cell phenotypes identified by scRNA-seq.

An important merit of our study is the public availability of the scRNA-seq data and the stromal blueprint we describe, which can all be interactively accessed via our blueprint server. This will allow scientists to co-cluster their own scRNA-seq data together with blueprint data and assign each of their individual cells to a cellular phenotype. This can also be achieved by feeding our stromal blueprint dataset to established machine learning pipelines, e.g. CellAssign^80^, and assigning each new cell to the most likely proxy. Such strategy would indeed be highly relevant, as several of our cellular phenotypes are missed when a smaller number of cells is analysed. Interestingly, as illustrated for melanoma, pooling new with existing scRNA-seq data was even possible when a different single-cell technology was used. Similarly, this blueprint could serve as training matrix to estimate the prevalence from specific cell (sub)types in bulk tissue transcriptomes using newly developed deconvolution methods, i.e. CIBERSORTx^81^. This is important, as bulk RNA-seq data of tumour tissues are often available for multiple large and homogeneous cohorts of cancer patients.

We also built trajectories between relevant cell phenotypes, highlighting how several of these do not represent separate entities. Stratification of these trajectories for cancer type revealed some intriguing differences. For instance, LC contained more exhausted CD8^+^ cytotoxic T-cells in the C1_CD8_HAVCR2 trajectory. Moreover, LC appeared more inflammatory as it was enriched for differentiated myeloid cells along both the CCL18 and MMP9 lineage. Also, memory B-cells were more differentiated in LC, while cDC2s got stuck early in the trajectory in OvC. Most probably, these differences are due to the fact that LC is an immune-infiltrated cancer with a high tumour mutation burden (TMB) and neoepitope load^82^, while OvC and CRC are cold tumours with a low TMB.

We believe our blueprint is also useful when monitoring dynamic changes in the TME during cancer treatment. Indeed, by performing scRNA-seq on individual biopsies obtained before and during treatment, individual cells can be assigned to each phenotypic cluster and changes can easily be interpreted in the context of the blueprint. For instance, when re-analysing a set of pre-*versus* on-treatment biopsies from melanomas exposed ICIs, we observed that exhausted CD8^+^ T-cells became gradually more common during treatment, while naïve CD4^+^ T-cells became less common. Notably, these shifts were only observed in patients responding to the treatment. Although findings that naïve CD4^+^ helper T-cells predict checkpoint immunotherapy are novel, these findings are not unexpected. Firstly, CD4^+^ helper T-cells can also express PD1, and are thus targeted by the treatment. Furthermore, they can enhance CD8^+^ T-cell infiltration^83^, improve antibody penetration^84^, T-cell memory formation, or have a direct cytolytic capacity^85^. Several other studies suggest the role of both naïve CD4^+^ and CD8^+^ T-cells in priming anti-tumour activity^86^. Overall, we believe that our approach to monitor how blueprint phenotypes change in response to cancer treatment and gradually also contribute to therapeutic resistance, will allow scientists to gain important insights into the mechanisms of action of novel cancer drugs.

## Materials and Methods

### Patients

This study was approved by the local ethics committee at the University Hospital Leuven for each cancer type. Only patients provided with informed consent were included in this study. The clinical information of all patients was summarised in **Table S1**.

### Preparation of single-cell suspensions

Following resection, samples from the tumour and adjacent non-malignant tissue were rapidly processed for single-cell RNA-sequencing. Samples were rinsed with PBS, minced on ice to pieces of <1 mm^3^ and transferred to 10 ml digestion medium containing collagenase P (2mg ml^-1^, ThermoFisher Scientific) and DNAse I (10U µl^-1^ Sigma) in DMEM (ThermoFisher Scientific). Samples were incubated for 15 min at 37 °C, with manual shaking every 5min. Samples were then vortexed for 10 s and pipetted up and down for 1 min using pipettes of descending sizes (25 ml, 10 ml and 5 ml). Next, 30 ml ice-cold PBS containing 2% fetal bovine serum was added and samples were filtered using a 40µm nylon mesh (ThermoFisher Scientific). Following centrifugation at 120×g and 4 °C for 5 min, the supernatant was decanted and discarded, and the cell pellet was resuspended in red blood cell lysis buffer. Following a 5-min incubation at room temperature, samples were centrifuged (120×g, 4°C, 5 min) and resuspended in 1 ml PBS containing 8 µl UltraPure BSA (50mg ml^−1^; AM2616, ThermoFisher Scientific) and filtered over Flowmi 40µm cell strainers (VWR) using wide-bore 1 ml low-retention filter tips (Mettler-Toledo). Next, 10 µl of this cell suspension was counted using an automated cell counter (Luna) to determine the concentration of live cells. The entire procedure was completed in less than 1 h (typically about 45 min).

### Single cell RNA-seq data acquisition and pre-processing

Libraries for scRNA-seq were generated using the Chromium Single Cell 3’ or 5’ library and Gel Bead & Multiplex Kit from 10x Genomics (**Table S2**). We aimed to profile 5,000 cells per library (if sufficient cells were retained during dissociation). All libraries were sequenced on Illumina NextSeq, HiSeq4000 or NovaSeq6000 until sufficient saturation was reached (73.8% on average, **Table S2**). After quality control, raw sequencing reads were aligned to the human reference genome GRCh38 and processed to a matrix representing the UMI’s per cell barcode per gene using CellRanger (10x Genomics, v2.0).

### Single-cell RNA analysis to determine major cell types and cell phenotypes

Raw gene expression matrices generated per sample were merged and analysed with the Seurat package (v2.3.4). Matrices were filtered by removing cell barcodes with <401 UMIs, <201 expressed genes, >6,000 expressed genes or >25% of reads mapping to mitochondrial RNA. The remaining cells were normalized and genes with a normalized expression between 0.125 and 3, and a quantile-normalized variance >0.5 were selected as variable genes. The number of variably-expressed genes differs for each clustering step (**Table S4**). When clustering cell types, we regressed out confounding factors: number of UMIs, % of mitochondrial RNA, patient ID and cell cycle (S and G2M phase scores calculated by the CellCycleScoring function in Seurat). After regression for confounding factors, all variably-expressed genes were used to construct principal components (PCs) and PCs covering the highest variance in the dataset were selected. The selection of these PCs was based on elbow and Jackstraw plots. Clusters were calculated by the FindClusters function with a resolution between 0.2 and 2, and visualised using the t-SNE dimensional reduction method. Differential gene-expression analysis was performed for clusters generated at various resolutions by both the Wilcoxon rank sum test and Model-based Analysis of Single-cell Transcriptomics (MAST) using the FindMarkers function. A specific resolution was selected when known cell types were identified as a cluster at a given resolution, but not at a lower resolution (**Table S5**), with the minimal constraint that each cluster has at least 10 significantly differentially expressed genes (FDR <0.01 with both methods) with at least a 2-fold difference in expression compared to all other clusters. Annotation of the resulting clusters to cell types was based on the expression of marker genes (Supplementary information, **Fig. S1c**). All major cell types were identified in one clustering step, except for DCs; pDCs co-clustered with B-cells, while other DCs co-clustered with myeloid cells. Therefore, we first separated DCs per cancer type based on established marker genes (pDC: *LILRA4* and *CXCR3*; cCDs: *CLEC9A, XCR1, CD1C, CCR7, CCL17, CCL19*, Langerhans-like: *CD1A, CD207*) ^2,40^ and then pooled these DCs for subclustering.

Next, all cells assigned to a given cell type per cancer type were merged and further subclustered into functional phenotypes using the same strategy, which we refer to as the unaligned clustering approach in the manuscript. However, the confounding factors used for cell types were not sufficient to reduce patient-specific effects when performing the subclustering. Instead of directly applying an unsupervised batch correction algorithm, we found that the interferon response (BROWNE_INTERFERON_RESPONSIVE_GENES in the Molecular Signatures Database or MSigDB v6.2) and the sample dissociation-induced gene signatures^87^ represent common patient-specific confounders, which were therefore regressed out. We additionally regressed out the hypoxia signature^88^ for myeloid cells to avoid clusters driven by hypoxia state instead of its origin or (anti-)inflammatory functions. Since hemoglobin and immunoglobulin genes are common contaminants from ambient RNA, hemoglobin genes were excluded for PCA. This also applied to immunoglobulin genes, except when subclustering B-cells. For T-cell subclustering, variable genes of T-cell receptor (*TRAV*s, *TRBV*s, *TRDV*s, *TRGV*s) were excluded to avoid somatic hypermutation associated variances. Similarly, variable genes of B-cell receptor (*IGLV*s, *IGKV*s, *IGHV*s) were all excluded when subclustering B-cells.

To reveal similarities between the subclusters across cancer types, we performed canonical correlation analysis (CCA, RunMultiCCA function) by aligning data from different cancer types into a subspace with the maximal correlation^11^. The selection of CCA dimensions or canonical correction vectors (CCs) for subspace alignment were guided by the CC bicor saturation plot (MetageneBicorPlot function). Resolution was determined similar to the PCA-based approach described above, followed by marker gene-based cluster annotation. Since CCA is designed to identify shared clusters, we performed CCA alignment without cancer-type specific cells defined by PCA-based approach for fibroblasts and myeloid cells. Low quality clusters were identified based on the number of detected genes within subclusters and the lack of marker genes. Doublet clusters expressed marker genes from other cell lineages, and had a higher than expected (3.9% according to the User Guide from 10x Genomics) doublets rate, as predicted by the artificial k-nearest neighbours algorithm implemented in DoubletFinder (v1.0)^89^. We also used Scrublet^90^ to identify doublet cells and could predict the same clusters as predicted by DoubletFinder. As an example, we evaluate for each of the B-cell clusters, *i)* the expression of marker genes from other cell types, *ii)* the higher number of detected genes, and *iii)* the overlap of cells predicted to be doublets by DoubletFinder and Scrublet (Supplementary Information, **Fig. S11a-d**).

For a comprehensive statistical analysis, we used a single-cell specific method based on mixed-effects modelling of associations of single cells (MASC) (Fonseka et al., 2018). The analysis systematically addressed two major questions: which cell types are enriched or depleted in all cancers or in a particular cancer type, and which cell types or stromal phenotypes are enriched or depleted in tumours versus normal tissue in all cancers or in a particular cancer type. Events with FDR < 0.05 were considered significant as summarised in **Table S6**.

### SCENIC analysis

Transcription factor (TF) activity was analysed using SCENIC (v1.0.0.3) per cell type with raw count matrices as input. The regulons and TF activity (AUC) for each cell were calculated with the pySCENIC (v0.8.9) pipeline with motif collection version mc9nr. The differentially activated TFs of each subcluster were identified by the Wilcoxon rank sum test against all the other cells of the same cell type. TFs with log-fold-change >0.1 and an adjusted p-value <1e-5 were considered as significantly upregulated.

### Trajectory inference analysis

We applied the Monocle (v2.8.0) algorithm to determine the potential lineage between diverse stromal cell phenotypes^91^. Seurat objects were imported to Monocle using importCDS function. DDRTree-based dimension reduction was performed with conserved and differentially expressed genes. These genes were calculated for each subcluster across LC, CRC and OvC using FindConservedMarkers function in Seurat using the metap (v1.0) algorithm and Wilcoxon rank sum test (max_pval < 0.01, minimum_p_val < 1e-5). PC selection was determined using the PC variance plot (plot_pc_variance_explained function in Monocle, 3-5 PCs). Genes with branch-dependent expression dynamics were calculated using the BEAM test in Monocle. Genes with a q-value <1e-10 were plotted in heatmaps. The dynamics of transcription factor activity (or AUC) was calculated by SCENIC and plotted per branch of trajectory along the pseudotime calculated by Monocle. For each TF, the AUC and pseudotime, smoothed as a natural spline using sm.ns function, were fitted in vector generalised linear model (VGLM) using VGAM package v1.1. TF with q-value <1e-50 were selected for plotting. Two other trajectory inference pipelines, i.e., Slingshot and SCORPIUS^92,93^, were also used. Since SCORPIUS cannot handle branched trajectories, we analysed both trajectories separately with the branching topology informed by Monocle analysis. To assess consistency between these pipelines, scaled pseudotime between Monocle, Slingshot and SCORPIUS were compared and high correlations were consistently observed between all lineages. Additionally, we compared expression of key marker genes along the trajectories of all 3 tools (Supplementary Information, **Fig. S12a-k**).

### Metabolic and cancer hallmark pathways and geneset enrichment analysis

Metabolic pathway activities were estimated with gene signatures from a curated database^94^. For robustness of the analysis, lowly expressed genes (< 1% cells) or genes shared by multiple pathways were trimmed. And pathways with less than 3 genes were excluded. Cancer hallmark gene sets from Molecular Signatures Database (MSigDB v6.1) were used. The activity of individual cells for each gene set was estimated by AUCell package (v1.2.4). The differentially activated pathways of each subcluster were identified by running the Wilcoxon rank sum test against other cells of the same cell type. Pathways with log-fold-change > 0.05 and an adjusted p-value < 0.01 were considered as significantly upregulated. GO and REACOTOME geneset enrichment analysis were performed using hypeR package^95^, geneset over-representation was determined by hypergeometric test.

### CITE-seq

We adopted the established CITE-seq protocol^79^ with some modifications. Briefly, 100,000–500,000 single cells of breast tumours were suspended in 100μl staining buffer (2% BSA, 0.01% Tween in PBS) before adding 10µl Fc-blocking reagent (FcX, BioLegend). and incubating during 10 min on ice. This was followed by the addition of 25µl TotalSeq-A (Biolegend) antibody-oligo pool (1:1000 diluted in staining buffer) and another 30 min incubation on ice. Cells were washed 3 times with staining buffer and filtered through a 40µm flowmi strainer before processing with 3’-scRNA-seq library kits. ADT (Antibody-Derived Tags) additive primers were added to increase yield of the ADT product. ADT-derived and mRNA-derived cDNAs were separated by SPRI purification and amplified for library construction and subsequent sequencing. For each cell barcode detected in the corresponding RNA library, ADTs were counted in the raw sequencing reads of CITE-seq experiments using CITE-seq-Count version 1.4. In the resulting UMI per ADT matrix, the noise level was calculated for each cell by taking the average signal increased with 3x the standard deviation of 10 control probes. Signals below this level were excluded. We divided the UMIs by the total UMI count for each cell to account for differences in library size and a centred log-ratio (CLR) normalization specific for each gene was computed. Clustering of protein data was done using the Euclidean distance matrix between cells and t-SNE coordinates were calculated using this distance matrix. The random forest algorithm incorporated in Seurat was iteratively applied on a training and test set, consisting of 67% and 33% of cells respectively, to predict cell type and T-/NK-cell phenotypes.

### Immunofluorescence assay and analysis

A 5µm-section of a formalin-fixed, paraffin-embedded (FFPE) microarray containing 14 melanoma metastasis from 9 patients was stained with antibodies against SOX10 (SCBT; sc-365692), CD4 (abcam; ab133616), CD31 (LSBio; LS-C173974) and TCF7 (R&D systems; AF5596) at a concentration of 1 µg/ml according to the Multiple Iterative Labeling by Antibody Neodeposition (MILAN) protocol, as described^96^.

### Tumour mutation detection

Whole-exome sequencing was performed as described previously^97^. The average sequencing depth was 161±67x coverage. Mutation of CRC samples were detected using Illumina Trusight26 Tumour kit.

## Data Availability

Raw sequencing reads of the single-cell RNA experiments have been deposited in the ArrayExpress database at EMBL-EBI and will be made accessible upon publication. An interactive web server for scRNA-seq data visualisation and exploration, based on SCope package^98^, is available at http://blueprint.lambrechtslab.org.

## Acknowledgements

We thank T. Van Brussel, R. Schepers, and E. Vanderheyden for technical assistance. This work was supported by a VIB TechWatch Grant to D.L. and B.T., ERC Consolidator Grants to D.L. (CHAMELEON), Funds for Research - Flanders grants to D.L. (G065615N), KU Leuven grants to D.L. and to B.T. (BOFZAP) and a VIB Grand Challenge grant to D.L. The computational resources used in this work were provided by the Flemish Supercomputer Center (VSC), funded by the Hercules Foundation and the Flemish Government, Department of Economy, Science and Innovation (EWI) and Stichting tegen Kanker (STK).

## Author Contributions

J.Q. and D.L. designed and supervised the study and wrote the manuscript; J.Q. and B.B. performed data analysis with significant contributions from P.B. and J.X.; I.V., A.S., S.T. and E.W. coordinated sample collection and clinical annotation with assistance from S.O., H.V., E.E., V.P., S.V., A.B., M.V.B., A.F. and G.F.; F.M.B., Y.V.H. and A.A. performed MILAN for melanoma samples. Dam.L. and B.T. contributed with critical data interpretation. All the authors have read the manuscript and provided useful comments.

