## Supplemental Figures 1-12 and Tables 1-2 for "A Pan-cancer Blueprint of the Heterogeneous Tumour Microenvironment Revealed by Single-Cell Profiling"

**Items Included**

**Supplementary Figures**

Fig. S1. Major cell type clustering
Fig. S2. Profiling of endothelial cells
Fig. S3. Characterization of fibroblasts
Fig. S4. Subclustering of dendritic cells
Fig. S5. Delineation of B-cell taxonomy
Fig. S7. Profiling of monocytes, macrophages and neutrophils
Fig. S8. Characterization of mast cells
Fig. S9. Mapping the breast cancer TME
Fig. S10. Mapping the melanoma SMART-seq2 dataset
Fig. S11. Doublet cluster prediction for B-cell subclusters
Fig. S12. Validation of trajectory inference results using Monocle, SCORPIUS and Slingshot

**Supplementary Tables**

Table S1. Characteristics of the patients included in this study
Table S2. Sequencing metrics of the samples included in this study

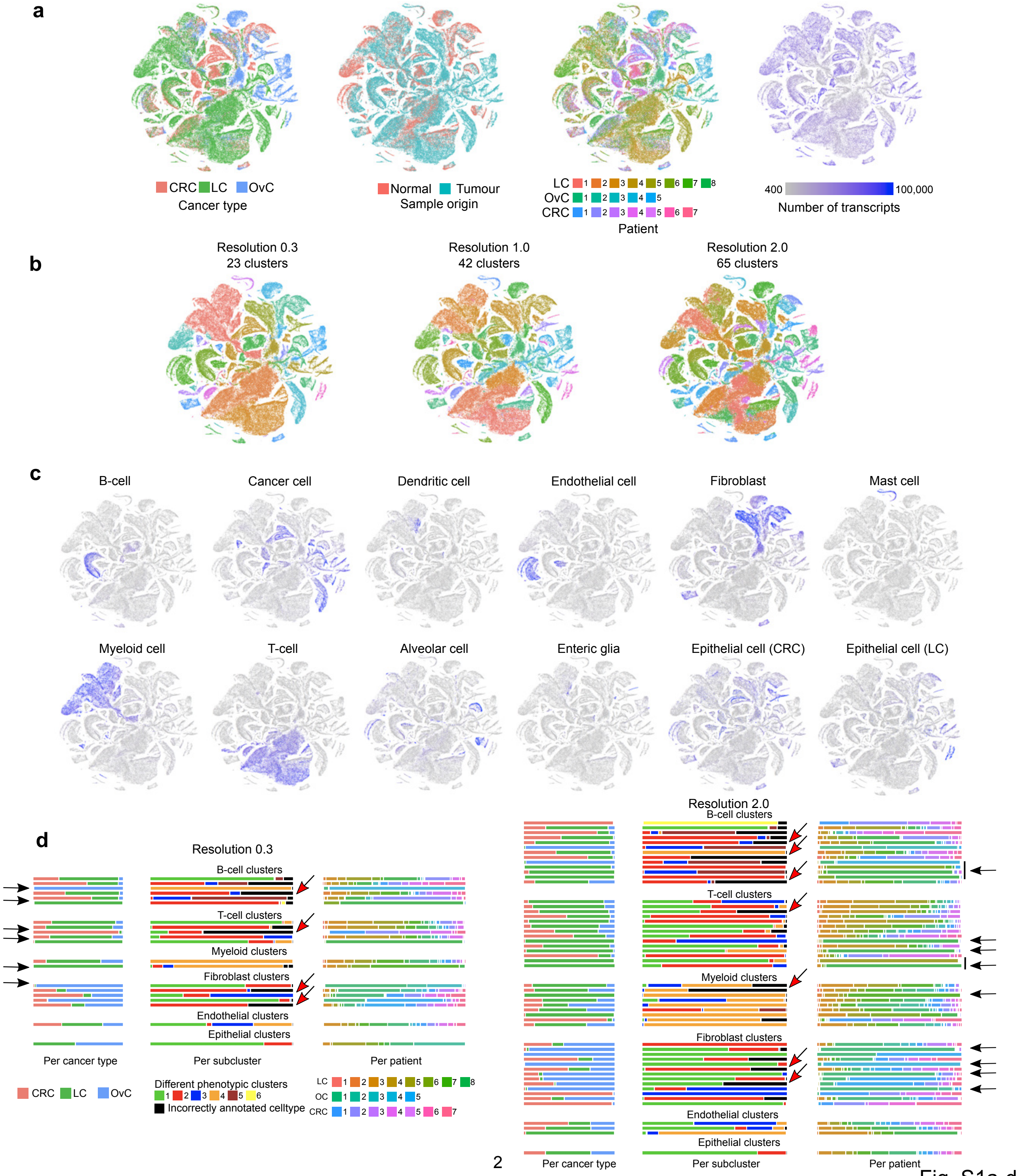

a
CRC LC OvC
Cancer type
Normal Tumour
Sample origin
LC 1 2 3 4 5 6 7 8
OvC 1 2 3 4 5
CRC 1 2 3 4 5 6 7
Patient
400 100,000
Number of transcripts
b
Resolution 0.3
23 clusters
Resolution 1.0
42 clusters
Resolution 2.0
65 clusters
c
B-cell
Cancer cell
Dendritic cell
Endothelial cell
Fibroblast
Mast cell
Myeloid cell
T-cell
Alveolar cell
Enteric glia
Epithelial cell (CRC)
Epithelial cell (LC)
d
Resolution 0.3
B-cell clusters
T-cell clusters
Myeloid clusters
Fibroblast clusters
Endothelial clusters
Epithelial clusters
Per cancer type
Per subcluster
Per patient
CRC LC OvC
Different phenotypic clusters
1 2 3 4 5 6
Incorrectly annotated celltype
LC 1 2 3 4 5 6 7 8
OC 1 2 3 4 5
CRC 1 2 3 4 5 6 7
Resolution 2.0
B-cell clusters
T-cell clusters
Myeloid clusters
Fibroblast clusters
Endothelial clusters
Epithelial clusters
Per cancer type
Per subcluster
Per patient




Fig. S1a-d

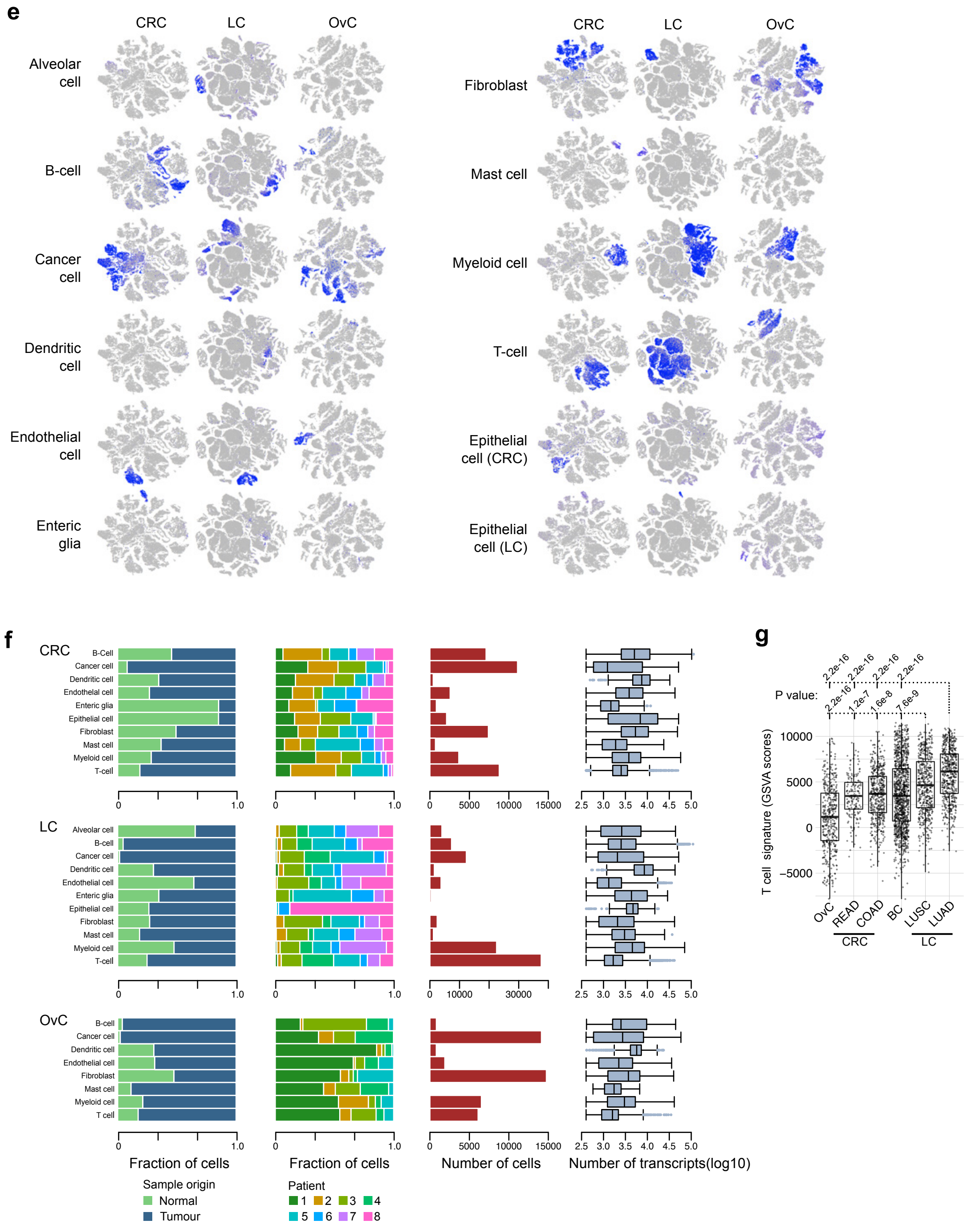
e
CRC
LC
OvC
Alveolar cell
B-cell
Cancer cell
Dendritic cell
Endothelial cell
Enteric glia
Fibroblast
Mast cell
Myeloid cell
T-cell
Epithelial cell (CRC)
Epithelial cell (LC)
f
Fraction of cells
Number of cells
Number of transcripts(log10)
Sample origin
Normal
Tumour
Patient
1 2 3 4 5 6 7 8
g
P value:
2.2e-16
1.2e-7
1.6e-8
7.6e-9
T cell signature (GSVA scores)
OvC
READ
COAD
BC
LUSC
LUAD
CRC
LC



Fig. S1e-g

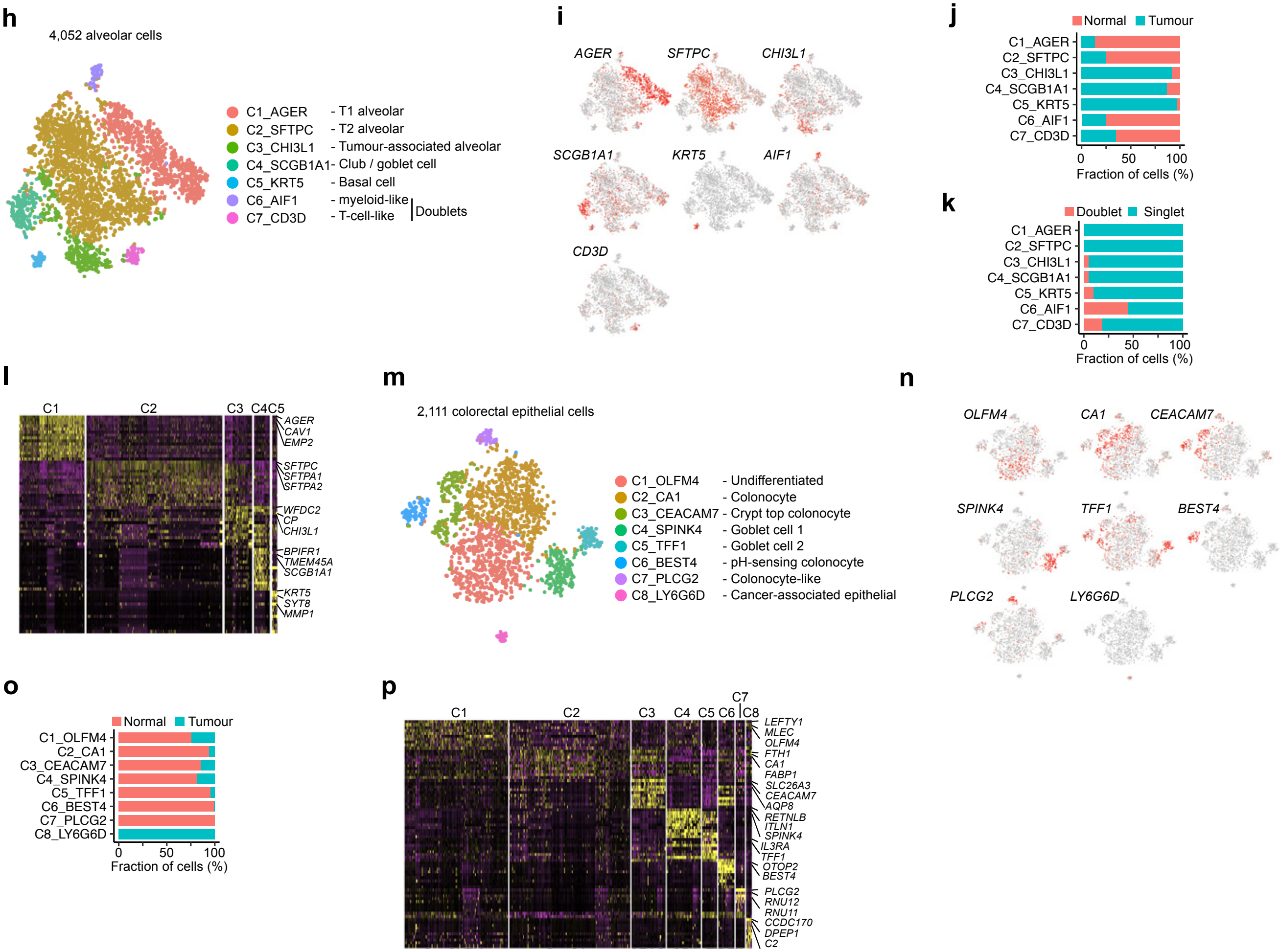
h
4,052 alveolar cells
C1_AGER - T1 alveolar
C2_SFTPC - T2 alveolar
C3_CHI3L1 - Tumour-associated alveolar
C4_SCGB1A1 - Club / goblet cell
C5_KRT5 - Basal cell
C6_AIF1 - myeloid-like
C7_CD3D - T-cell-like
Doublets
i
AGER
SFTPC
CHI3L1
SCGB1A1
KRT5
AIF1
CD3D
j
Normal
Tumour
C1_AGER
C2_SFTPC
C3_CHI3L1
C4_SCGB1A1
C5_KRT5
C6_AIF1
C7_CD3D
0 50 100
Fraction of cells (%)
k
Doublet
Singlet
C1_AGER
C2_SFTPC
C3_CHI3L1
C4_SCGB1A1
C5_KRT5
C6_AIF1
C7_CD3D
0 50 100
Fraction of cells (%)
l
C1 C2 C3 C4C5
AGER
CAV1
EMP2
SFTPC
SFTPA1
SFTPA2
WFDC2
CP
CHI3L1
BPIFR1
TMEM45A
SCGB1A1
KRT5
SYT8
MMP1
m
2,111 colorectal epithelial cells
C1_OLFM4 - Undifferentiated
C2_CA1 - Colonocyte
C3_CEACAM7 - Crypt top colonocyte
C4_SPINK4 - Goblet cell 1
C5_TFF1 - Goblet cell 2
C6_BEST4 - pH-sensing colonocyte
C7_PLCG2 - Colonocyte-like
C8_LY6G6D - Cancer-associated epithelial
n
OLFM4
CA1
CEACAM7
SPINK4
TFF1
BEST4
PLCG2
LY6G6D
o
Normal
Tumour
C1_OLFM4
C2_CA1
C3_CEACAM7
C4_SPINK4
C5_TFF1
C6_BEST4
C7_PLCG2
C8_LY6G6D
0 50 100
Fraction of cells (%)
p
C1 C2 C3 C4 C5 C6 C7 C8
LEFTY1
MLEC
OLFM4
FTH1
CA1
FABP1
SLC26A3
CEACAM7
AQP8
RETNLB
ITLN1
SPINK4
IL3RA
TFF1
OTOP2
BEST4
PLCG2
RNU12
RNU11
CCDC170
DPEP1
C2

Fig. S1h-p

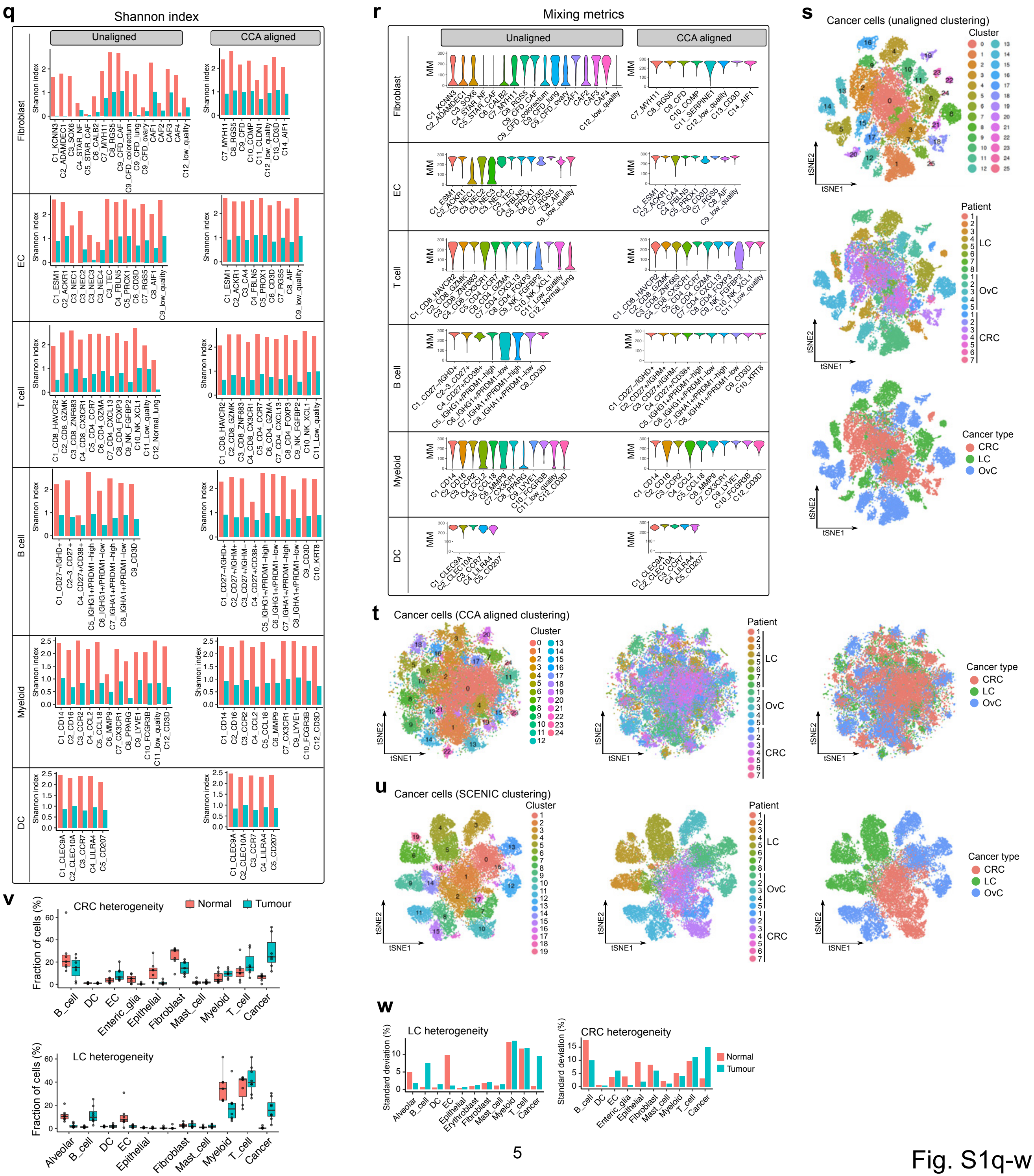
q
Shannon index
Unaligned
CCA aligned
Fibroblast
EC
T cell
B cell
Myeloid
DC
r
Mixing metrics
Unaligned
CCA aligned
MM
s
Cancer cells (unaligned clustering)
Cluster
Patient
LC
OvC
CRC
Cancer type
tSNE1
tSNE2
t
Cancer cells (CCA aligned clustering)
u
Cancer cells (SCENIC clustering)
v
CRC heterogeneity
Normal
Tumour
Fraction of cells (%)
LC heterogeneity
w
Standard deviation (%)



Fig. S1q-w

**Fig. S1. Major cell type clustering**

**a** t-SNE plots of 183,376 single cells colour-coded per cancer type, sample of origin, patient, and number of transcripts. **b** t-SNE plots colour-coded by different clusters generated by PCA based clustering at different resolutions. **c** Expression of metagenes projected on t-SNE plots. Composition of the metagenes: B-cell (*CD79A*, *CD79B*), cancer (*EPCAM*, *KRT7*, *KRT18*), DC (*CD1C*, *CD207*, *CLEC9A*, *LILRA4*, *CCL17*), EC (*CLDN5*, *PECAM1*, *VWF*), fibroblast (*COL1A*, *BGN*, *DCN*), mast cell (*MS4A2*, *TPSAB1*, *CPA3*), myeloid (*CD68*, *LYZ*, *AIF1*), T-cell (*CD3D*, *CD3E*, *CD3G*), alveolar cell (*CLDN18*, *SFTPA1*, *SFTPA2*, *SFTPC*), enteric glia (*S100B*, *PLP1*), epithelial cell (colorectal, *MT1E*, *MT1G*, *ITLN1*, *ZG16*), epithelial cell (lung, *CAPS*, *TPP3*). **d** Barplots ordered per major cell type at resolutions 0.3 (left column) and 2.0 (right column). The barplots represent the composition per cancer type (left panel), the annotation per subcluster, as previously described (middle panel) [2] and the contribution per patient (right panel). Cancer and tissue specific clusters (alveolar) are excluded. At the lower resolution arrows indicate the tissue specific clusters (black arrows), at the higher resolution arrows indicate patient specific clusters (black arrows). Cells clustered into an incorrect major cell type are highlighted by red arrows at both resolutions. **e** Expression of metagenes projected on t-SNE plots per cancer type. Same metagenes were used as in (**c**). **f** Barplots representing per cell type in different cancers from left to right: fraction of cells per origin, fraction of cells per patient, number of cells, and the total number of detected transcripts. **g** Boxplot showing the prevalence of T-cells based on GSVA scores in bulk RNA-seq of different TCGA cancer types, including LC (both for LUAD and LUSC with 541 and 502 patients respectively), BC (1,119 patients), CRC (rectal cancer or READ and colorectal cancer or COAD with 167 and 483 patients respectively) and OvC (430 patients). *P*-values for LUAD and LUSC versus all other cancer types are depicted. **h** t-SNE plot showing 4,052 alveolar cells colour-coded by clusters. **i** t-SNE plots showing marker gene expressions of alveolar clusters. **j** Fraction of cells derived from normal (red) or tumour tissue (green) for each alveolar cluster. **k** Fraction of singlet and doublet cells in each alveolar cluster calculated by DoubletFinder. C6 and C7 have higher doublet percentage than other clusters. **l** Heatmap for differential gene expression of alveolar clusters. **m** t-SNE plot showing 2,111 colorectal epithelial cells colour-coded by clusters. **n** t-SNE plots showing marker gene expressions of colorectal epithelial cell clusters. **o** Fraction of cells derived from normal (red) or tumour tissue (green) for each colorectal epithelial cluster. **p** Heatmap for differential gene expression of colorectal epithelial cell clusters. **q-r** Shannon index (**q**) and mixing metrics (**r**) in all cell type subclusters after unaligned and CCA-aligned clustering; Red and green denotes the Shannon index for patient and cancer type, respectively. **s-u** t-SNEs of cancer cells colour-coded for each separate cluster (left), per individual patient (middle) and cancer type (right) by unaligned clustering (**s**), CCA aligned clustering (**t**) and SCENIC clustering that leveraging transcription factor activity calculated by SCENIC pipeline (**u**). **v** Boxplots showing the relative percentage of cancer cells and each stromal subcluster derived from normal (red) or tumour (green) in CRC and LC, respectively. **w** Standard deviation of the relative percentage of cancer cells and each stromal subcluster in normal and tumour tissue in CRC and LC, respectively. The standard deviation was considered a measure of inter-tumour heterogeneity.

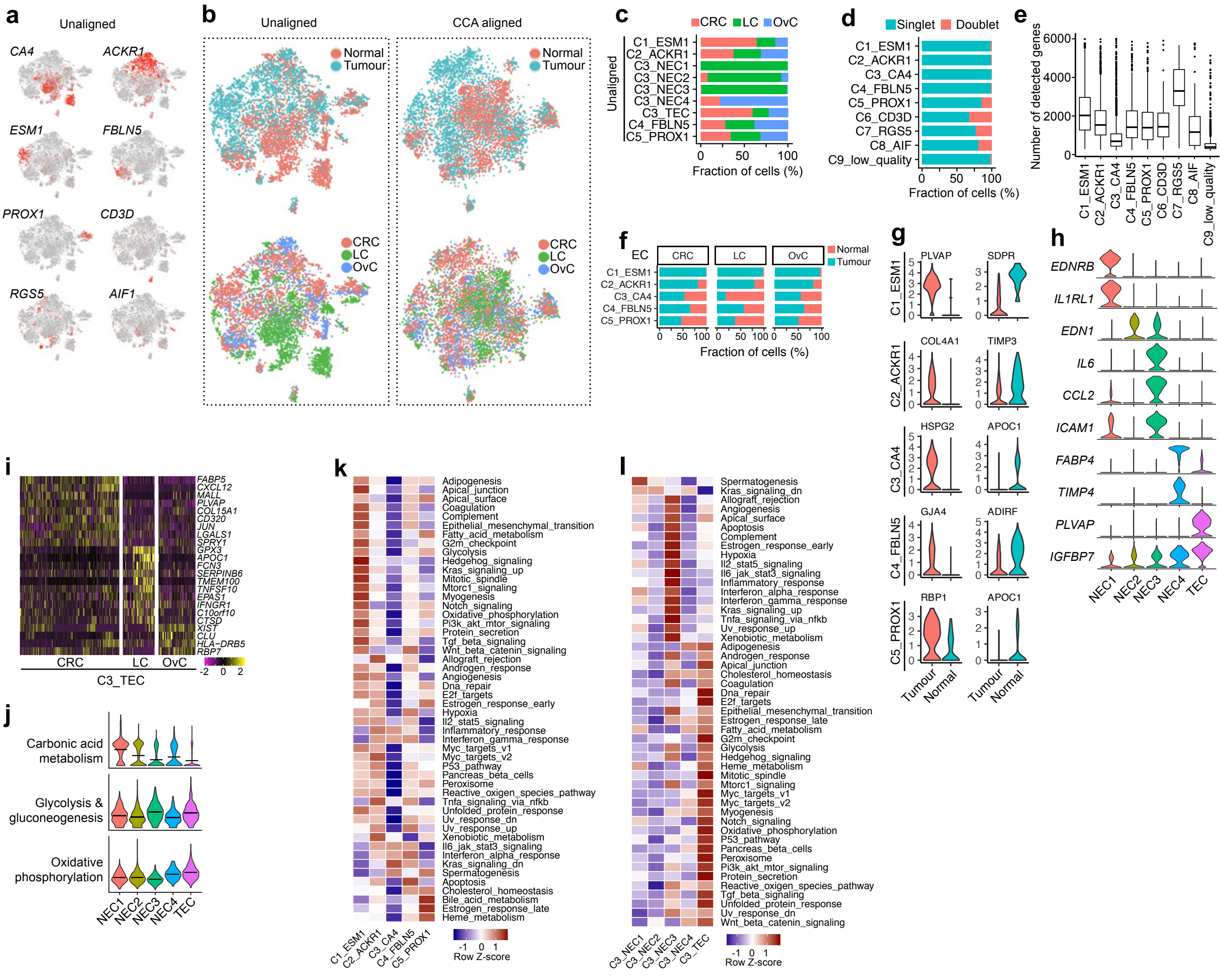
a
Unaligned
CA4
ACKR1
ESM1
FBLN5
PROX1
CD3D
RGS5
AIF1
b
Unaligned
CCA aligned
Normal
Tumour
CRC
LC
OvC
c
CRC LC OvC
Unaligned
C1_ESM1
C2_ACKR1
C3_NEC1
C3_NEC2
C3_NEC3
C3_NEC4
C3_TEC
C4_FBLN5
C5_PROX1
0 50 100
Fraction of cells (%)
d
Singlet Doublet
C1_ESM1
C2_ACKR1
C3_CA4
C4_FBLN5
C5_PROX1
C6_CD3D
C7_RGS5
C8_AIF
C9_low_quality
0 50 100
Fraction of cells (%)
e
Number of detected genes
0 2000 4000 6000
C1_ESM1
C2_ACKR1
C3_CA4
C4_FBLN5
C5_PROX1
C6_CD3D
C7_RGS5
C8_AIF
C9_low_quality
f
EC
CRC
LC
OvC
Normal
Tumour
C1_ESM1
C2_ACKR1
C3_CA4
C4_FBLN5
C5_PROX1
0 50 100
Fraction of cells (%)
g
C1_ESM1
PLVAP
SDPR
C2_ACKR1
COL4A1
TIMP3
C3_CA4
HSPG2
APOC1
C4_FBLN5
GJA4
ADIRF
C5_PROX1
RBP1
APOC1
Tumour
Normal
h
EDNRB
IL1RL1
EDN1
IL6
CCL2
ICAM1
FABP4
TIMP4
PLVAP
IGFBP7
NEC1
NEC2
NEC3
NEC4
TEC
i
FABP5
CXCL12
MALL
PLVAP
COL15A1
CD320
JUN
LGALS1
SPRY1
GPX3
APOC1
FCN3
SERPINB6
TMEM100
TNFSF10
EPAS1
IFNGR1
C10orf10
CTSD
XIST
CLU
HLA−DRB5
RBP7
CRC
LC
OvC
C3_TEC
-2 0 2
j
Carbonic acid metabolism
Glycolysis & gluconeogenesis
Oxidative phosphorylation
NEC1
NEC2
NEC3
NEC4
TEC
k
Adipogenesis
Apical_junction
Apical_surface
Coagulation
Complement
Epithelial_mesenchymal_transition
Fatty_acid_metabolism
G2m_checkpoint
Glycolysis
Hedgehog_signaling
Kras_signaling_up
Mitotic_spindle
Mtorc1_signaling
Myogenesis
Notch_signaling
Oxidative_phosphorylation
Pi3k_akt_mtor_signaling
Protein_secretion
Tgf_beta_signaling
Wnt_beta_catenin_signaling
Allograft_rejection
Androgen_response
Angiogenesis
Dna_repair
E2f_targets
Estrogen_response_early
Hypoxia
Il2_stat5_signaling
Inflammatory_response
Interferon_gamma_response
Myc_targets_v1
Myc_targets_v2
P53_pathway
Pancreas_beta_cells
Peroxisome
Reactive_oxigen_species_pathway
Tnfa_signaling_via_nfkb
Unfolded_protein_response
Uv_response_dn
Uv_response_up
Xenobiotic_metabolism
Il6_jak_stat3_signaling
Interferon_alpha_response
Kras_signaling_dn
Spermatogenesis
Apoptosis
Cholesterol_homeostasis
Bile_acid_metabolism
Estrogen_response_late
Heme_metabolism
C1_ESM1
C2_ACKR1
C3_CA4
C4_FBLN5
C5_PROX1
-1 0 1
Row Z-score
l
Spermatogenesis
Kras_signaling_dn
Allograft_rejection
Angiogenesis
Apical_surface
Apoptosis
Complement
Estrogen_response_early
Hypoxia
Il2_stat5_signaling
Il6_jak_stat3_signaling
Inflammatory_response
Interferon_alpha_response
Interferon_gamma_response
Kras_signaling_up
Tnfa_signaling_via_nfkb
Uv_response_up
Xenobiotic_metabolism
Adipogenesis
Androgen_response
Apical_junction
Cholesterol_homeostasis
Coagulation
Dna_repair
E2f_targets
Epithelial_mesenchymal_transition
Estrogen_response_late
Fatty_acid_metabolism
G2m_checkpoint
Glycolysis
Hedgehog_signaling
Heme_metabolism
Mitotic_spindle
Mtorc1_signaling
Myc_targets_v1
Myc_targets_v2
Myogenesis
Notch_signaling
Oxidative_phosphorylation
P53_pathway
Pancreas_beta_cells
Peroxisome
Pi3k_akt_mtor_signaling
Protein_secretion
Reactive_oxigen_species_pathway
Tgf_beta_signaling
Unfolded_protein_response
Uv_response_dn
Wnt_beta_catenin_signaling
C3_NEC1
C3_NEC2
C3_NEC3
C3_NEC4
C3_TEC
-1 0 1
Row Z-score



Fig. S2

**Fig. S2. Profiling of endothelial cells**

**a** t-SNE plots showing EC marker gene expressions for unaligned clusters. **b** t-SNE plots showing unaligned or CCA aligned clusters colour-coded by sample origin (upper panels) and cancer type (low panels). **c** Fraction of cells for unaligned EC clusters per cancer type. **d** Fraction of singlet and doublet cells in each EC subcluster calculated by DoubletFinder. **e** Box plot showing number of detected genes for each EC subcluster. **f** Fraction of cells derived from normal (red) or tumour tissue (green) for each EC subcluster in each cancer type. **g** Violin plots of genes specifically expressed in tumour or normal tissue for each of the 5 EC subclusters. **h** Violin plot showing marker gene expressions for capillary EC subclusters. **i** Heatmap of differentially expressed genes between CRC, LC and OvC in the C3_TEC clusters. **j** Violin plot for metabolic activity for capillary EC subclusters scored by AUCell. **k-l** Differences in cancer hallmark pathway activities scored per cell by AUCell for all major EC subclusters (**k**) or subclusters of capillary ECs (**l**).

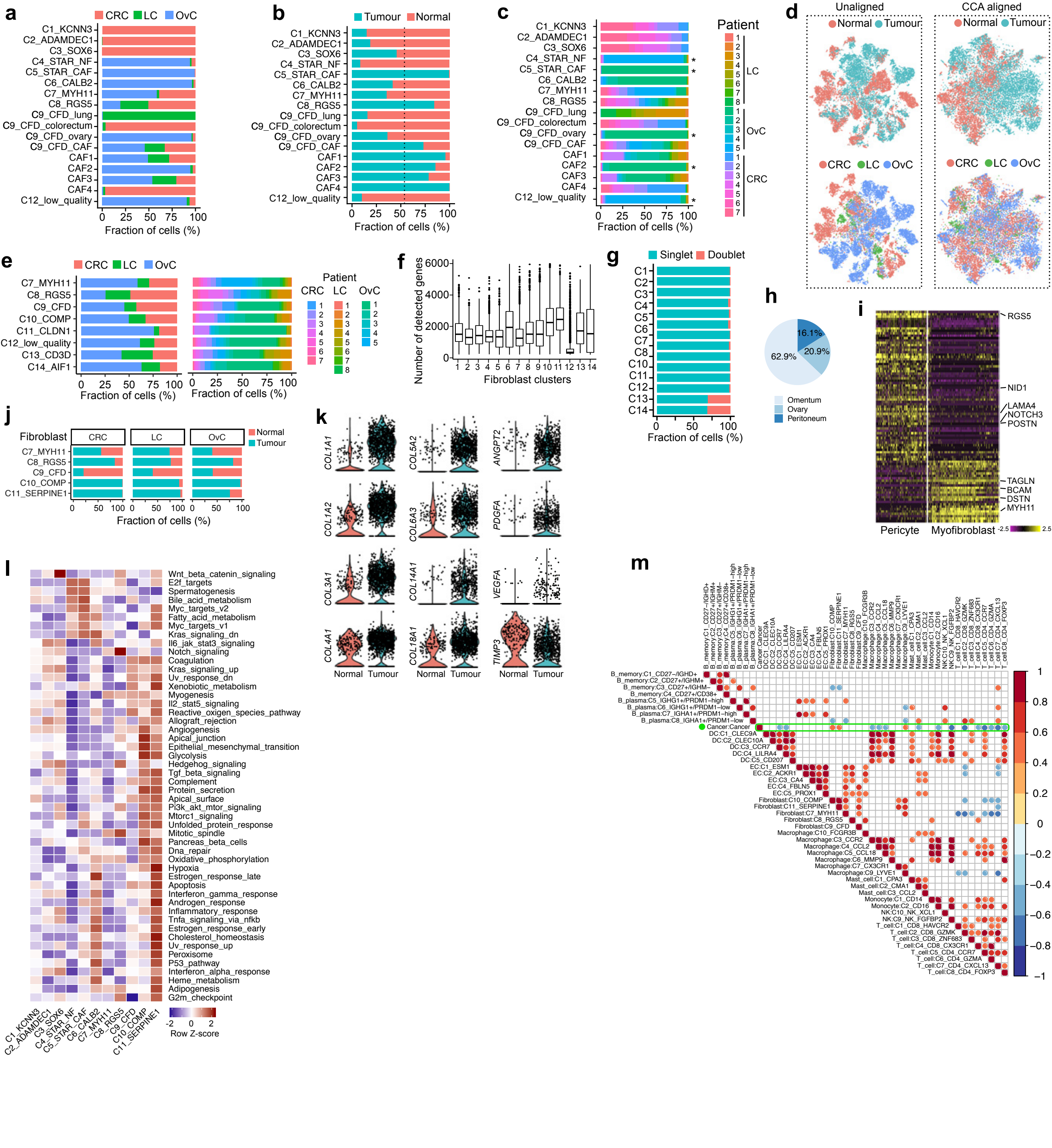
a
CRC
LC
OvC
C1_KCNN3
C2_ADAMDEC1
C3_SOX6
C4_STAR_NF
C5_STAR_CAF
C6_CALB2
C7_MYH11
C8_RGS5
C9_CFD_lung
C9_CFD_colorectum
C9_CFD_ovary
C9_CFD_CAF
CAF1
CAF2
CAF3
CAF4
C12_low_quality
Fraction of cells (%)
b
Tumour
Normal
c
Patient
d
Unaligned
CCA aligned
e
C7_MYH11
C8_RGS5
C9_CFD
C10_COMP
C11_CLDN1
C12_low_quality
C13_CD3D
C14_AIF1
f
Number of detected genes
Fibroblast clusters
g
Singlet
Doublet
h
62.9%
20.9%
16.1%
Omentum
Ovary
Peritoneum
i
RGS5
NID1
LAMA4
NOTCH3
POSTN
TAGLN
BCAM
DSTN
MYH11
Pericyte
Myofibroblast
j
Fibroblast
C11_SERPINE1
k
COL1A1
COL1A2
COL3A1
COL4A1
COL5A2
COL6A3
COL14A1
COL18A1
ANGPT2
PDGFA
VEGFA
TIMP3
l
Wnt_beta_catenin_signaling
E2f_targets
Spermatogenesis
Bile_acid_metabolism
Myc_targets_v2
Fatty_acid_metabolism
Myc_targets_v1
Kras_signaling_dn
Il6_jak_stat3_signaling
Notch_signaling
Coagulation
Kras_signaling_up
Uv_response_dn
Xenobiotic_metabolism
Myogenesis
Il2_stat5_signaling
Reactive_oxigen_species_pathway
Allograft_rejection
Angiogenesis
Apical_junction
Epithelial_mesenchymal_transition
Glycolysis
Hedgehog_signaling
Tgf_beta_signaling
Complement
Protein_secretion
Apical_surface
Pi3k_akt_mtor_signaling
Mtorc1_signaling
Unfolded_protein_response
Mitotic_spindle
Pancreas_beta_cells
Dna_repair
Oxidative_phosphorylation
Hypoxia
Estrogen_response_late
Apoptosis
Interferon_gamma_response
Androgen_response
Inflammatory_response
Tnfa_signaling_via_nfkb
Estrogen_response_early
Cholesterol_homeostasis
Uv_response_up
Peroxisome
P53_pathway
Interferon_alpha_response
Heme_metabolism
Adipogenesis
G2m_checkpoint
Row Z-score
m
Cancer:Cancer

Fig. S3

**Fig. S3. Characterization of fibroblasts**

**a-c** Fraction of cells in the unaligned fibroblast clusters colour-coded per cancer type (**a**), sample origin (**b**), or patient (**c**). **d** t-SNE plots showing unaligned and CCA aligned clusters colour-coded by sample origin (upper panels) and cancer type (low panels). **e** Fraction of cells in CCA-aligned fibroblast subclusters colour-coded per cancer type (left) or patient (right). **f** Number of detected genes for fibroblast clusters, combining both tissue-specific and shared subclusters. **g** Fraction of singlet and doublet cells detected for fibroblast clusters. **h** Fraction of mesothelium-derived cells from different tissues. **i** Heatmap of differentially expressed genes between pericytes and myofibroblasts. **j** Fraction of cells derived from normal (red) or tumour tissue (green) for shared fibroblast subclusters in each cancer type. **k** Violin plot showing differentially expressed genes from normal and tumour tissues. **l** Differences in cancer-hallmark pathway activities scored per cell by AUCell for fibroblast clusters. **m** Heatmap showing positive (red) and negative (blue) correlations between the prevalence of the different cell phenotypes in each patient. Only significant correlations ($p < 0.05$), as assessed by Pearson correlation, are shown. Positive correlations between cancer cells and CAFs (C10, C11) were detected.

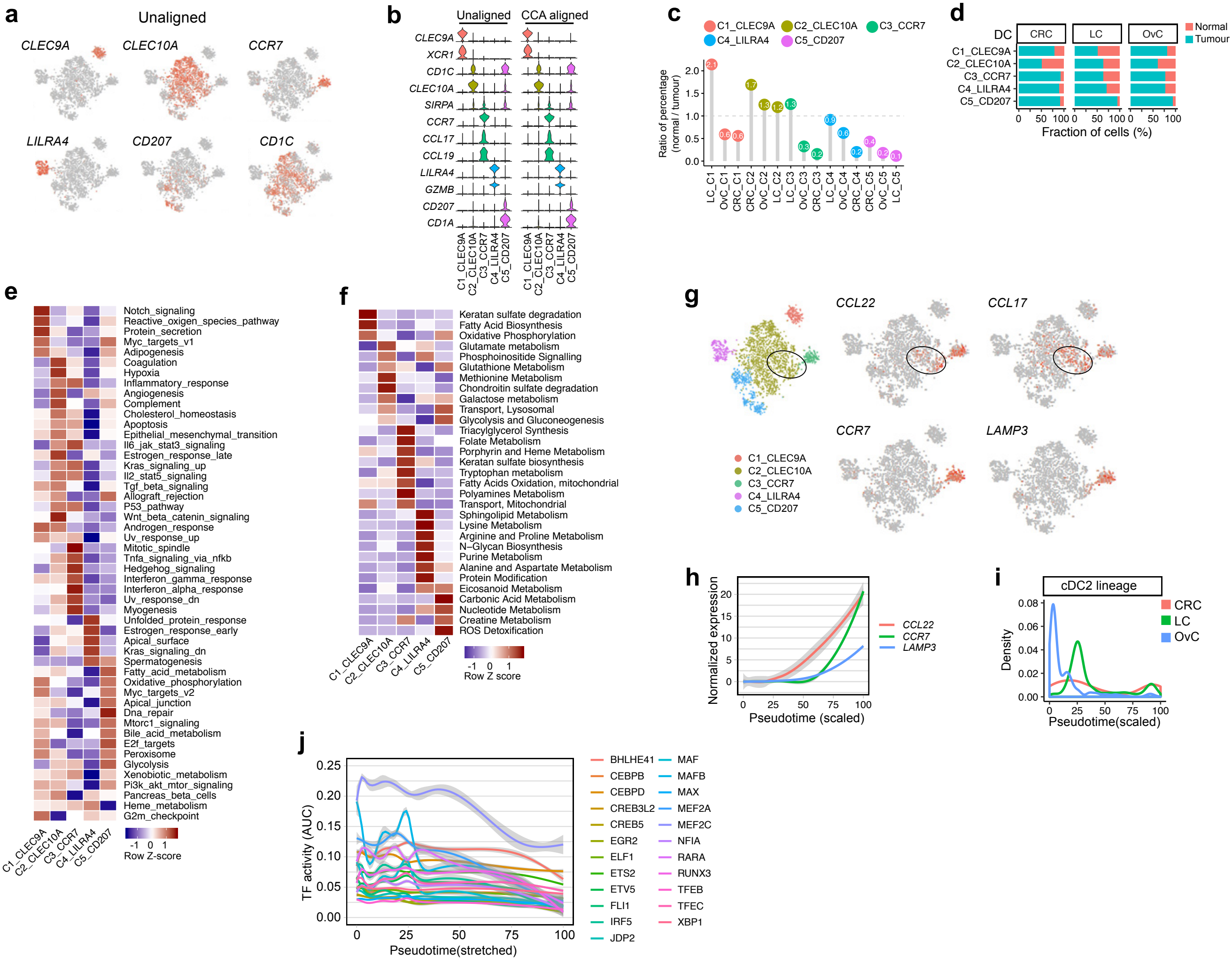
a
Unaligned
CLEC9A
CLEC10A
CCR7
LILRA4
CD207
CD1C
b
Unaligned
CCA aligned
CLEC9A
XCR1
CD1C
CLEC10A
SIRPA
CCR7
CCL17
CCL19
LILRA4
GZMB
CD207
CD1A
c
C1_CLEC9A
C2_CLEC10A
C3_CCR7
C4_LILRA4
C5_CD207
Ratio of percentage (normal / tumour)
d
DC
CRC
LC
OvC
Normal
Tumour
Fraction of cells (%)
e
Row Z-score
f
Row Z score
g
CCL22
CCL17
CCR7
LAMP3
h
Normalized expression
Pseudotime (scaled)
i
cDC2 lineage
Density
Pseudotime(scaled)
j
TF activity (AUC)
Pseudotime(stretched)



Fig. S4

**Fig. S4. Subclustering of dendritic cells**

**a** t-SNE plots showing marker gene expressions for unaligned DC clusters. **b** Violin plot showing marker gene expressions for both unaligned and CCA aligned DC clusters. **c** Ratio of relative percentage for DC subclusters between normal and tumour tissues. Less than 1 indicates tumour enrichment. Langerhans-like DCs were enriched in tumour (FDR=$1.5 \times 10^{-31}$). **d** Fraction of cells derived from normal (red) or tumour tissue (green) for each DC phenotype in each cancer type. **e** Differences in cancer-hallmark pathway activities scored per cell by AUCell for DC phenotypes. **f** Metabolic activity of each unaligned DC phenotype scored by AUCell. **g** t-SNEs of DCs colour-coded for marker genes: *CCL22* and *CCL17* (early activation markers of the C3_DC cluster), as well as *CCR7* and *LAMP3* (late activation markers of the C3_DC cluster). **h** Expression pattern of *CCL22*, *CCR7* and *LAMP3* along the trajectory pseudotime of the C2 to C3 trajectory. **i** Density plots for CRC, LC and OvC along the cDC2 trajectory. **j** Transcription factor inactivation dynamics during cDC2 to migratory cDC differentiation.

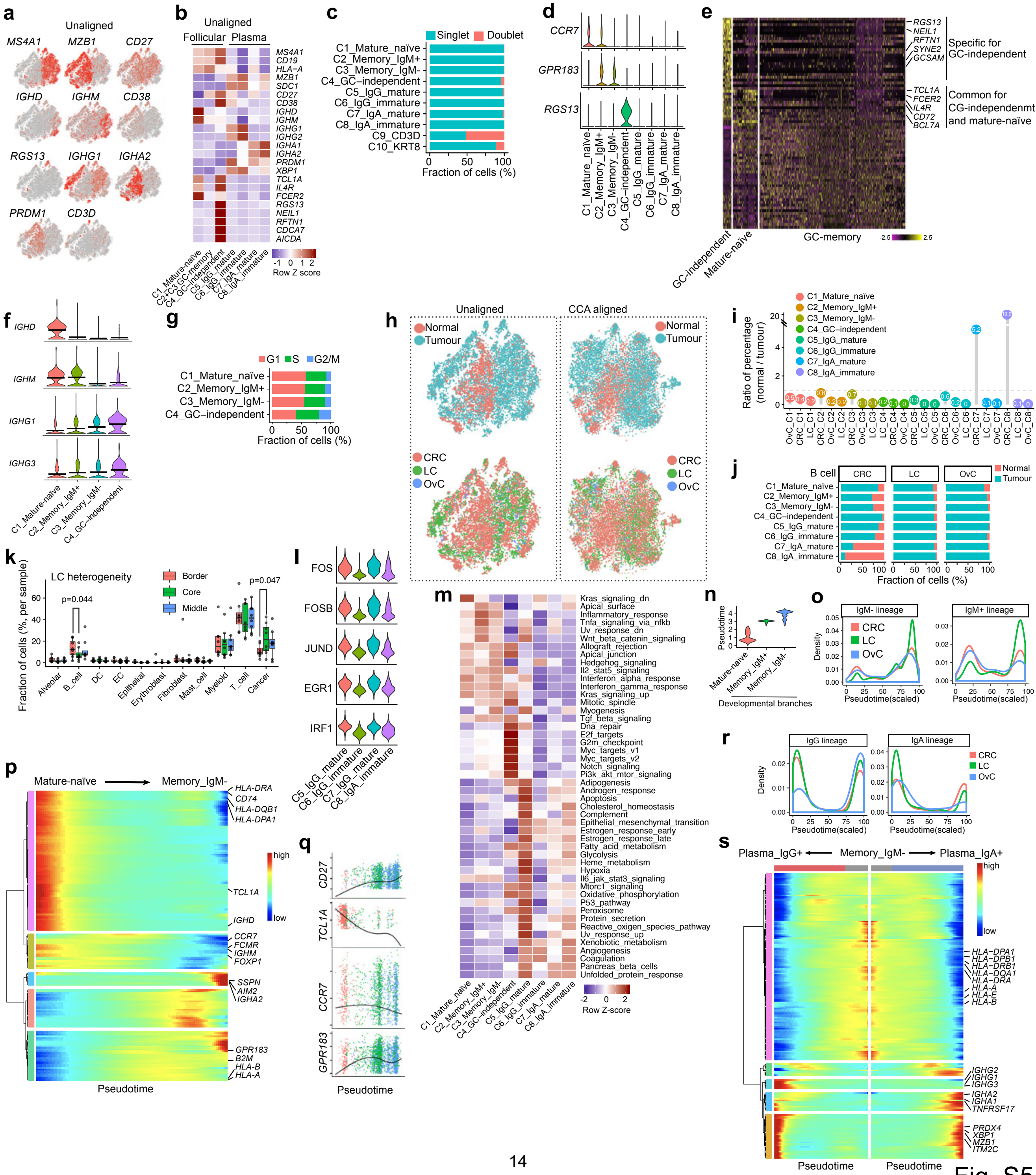
a
Unaligned
MS4A1
MZB1
CD27
IGHD
IGHM
CD38
RGS13
IGHG1
IGHA2
PRDM1
CD3D
b
Unaligned
Follicular
Plasma
MS4A1
CD19
HLA−A
MZB1
SDC1
CD27
CD38
IGHD
IGHM
IGHG1
IGHG2
IGHA1
IGHA2
PRDM1
XBP1
TCL1A
IL4R
FCER2
RGS13
NEIL1
RFTN1
CDCA7
AICDA
C1_Mature-naïve
C2+C3 GC-memory
C4_GC-independent
C5_IgG_mature
C6_IgG_immature
C7_IgA_mature
C8_IgA_immature
Row Z score
c
Singlet
Doublet
C1_Mature_naïve
C2_Memory_IgM+
C3_Memory_IgM-
C4_GC−independent
C5_IgG_mature
C6_IgG_immature
C7_IgA_mature
C8_IgA_immature
C9_CD3D
C10_KRT8
Fraction of cells (%)
d
CCR7
GPR183
RGS13
e
RGS13
NEIL1
RFTN1
SYNE2
GCSAM
Specific for GC-independent
TCL1A
FCER2
IL4R
CD72
BCL7A
Common for CG-independenmt and mature-naïve
GC-independent
Mature-naïve
GC-memory
f
IGHD
IGHM
IGHG1
IGHG3
C1_Mature-naïve
C2_Memory_IgM+
C3_Memory_IgM-
C4_GC-independent
g
G1
S
G2/M
Fraction of cells (%)
h
Unaligned
CCA aligned
Normal
Tumour
CRC
LC
OvC
i
Ratio of percentage (normal / tumour)
j
B cell
CRC
LC
OvC
Normal
Tumour
Fraction of cells (%)
k
LC heterogeneity
Border
Core
Middle
p=0.044
p=0.047
Fraction of cells (%, per sample)
Alveolar
B_cell
DC
EC
Epithelial
Erythroblast
Fibroblast
Mast_cell
Myeloid
T_cell
Cancer
l
FOS
FOSB
JUND
EGR1
IRF1
m
Kras_signaling_dn
Apical_surface
Inflammatory_response
Tnfa_signaling_via_nfkb
Uv_response_dn
Wnt_beta_catenin_signaling
Allograft_rejection
Apical_junction
Hedgehog_signaling
Il2_stat5_signaling
Interferon_alpha_response
Interferon_gamma_response
Kras_signaling_up
Mitotic_spindle
Myogenesis
Tgf_beta_signaling
Dna_repair
E2f_targets
G2m_checkpoint
Myc_targets_v1
Myc_targets_v2
Notch_signaling
Pi3k_akt_mtor_signaling
Adipogenesis
Androgen_response
Apoptosis
Cholesterol_homeostasis
Complement
Epithelial_mesenchymal_transition
Estrogen_response_early
Estrogen_response_late
Fatty_acid_metabolism
Glycolysis
Heme_metabolism
Hypoxia
Il6_jak_stat3_signaling
Mtorc1_signaling
Oxidative_phosphorylation
P53_pathway
Peroxisome
Protein_secretion
Reactive_oxigen_species_pathway
Uv_response_up
Xenobiotic_metabolism
Angiogenesis
Coagulation
Pancreas_beta_cells
Unfolded_protein_response
Row Z-score
n
Pseudotime
Mature-naïve
Memory_IgM+
Memory_IgM-
Developmental branches
o
IgM- lineage
IgM+ lineage
Density
Pseudotime(scaled)
p
Mature-naïve → Memory_IgM-
HLA-DRA
CD74
HLA-DQB1
HLA-DPA1
TCL1A
IGHD
CCR7
FCMR
IGHM
FOXP1
SSPN
AIM2
IGHA2
GPR183
B2M
HLA-B
HLA-A
Pseudotime
high
low
q
CD27
TCL1A
CCR7
GPR183
Pseudotime
r
IgG lineage
IgA lineage
Density
Pseudotime(scaled)
s
Plasma_IgG+ ← Memory_IgM- → Plasma_IgA+
HLA−DPA1
HLA−DPB1
HLA−DRB1
HLA−DQA1
HLA−DRA
HLA−A
HLA−E
HLA−B
IGHG2
IGHG1
IGHG3
IGHA2
IGHA1
TNFRSF17
PRDX4
XBP1
MZB1
ITM2C
Pseudotime
Pseudotime



Fig. S5

**Fig. S5. Delineation of B-cell taxonomy**

**a** t-SNE plots showing marker gene expressions for unaligned clusters. **b** Heatmap of functional gene sets for unaligned clusters. **c** Fraction of singlet and doublet cells predicted by DoubletFinder. **d** Violin plot showing expressions of GC-migration related genes. **e** Heatmap of differentially expressed genes between follicular B-cell subclusters. **f** Differentially expressed immunoglobulin genes between follicular B-cell subclusters, mean expression indicated as horizontal bars. **g** Fraction of cells for follicular B-cell subclusters with predicted cell cycle phases. **h** t-SNE plots for unaligned (left panels) and CCA-aligned (right panels) methods colour-coded by sample origin (upper panels), or cancer type (lower panels). **i** Ratio of relative % for B-cell subclusters by comparing normal with tumour tissues, <1 indicates tumour enrichment. Most B-cells (C1-C6) were enriched in tumour (FDR < $5.2x10^{-8}$), while $IgA^{+}$ plasma cells (C7-C8) were enriched in normal colon (FDR < $8.9x10^{-118}$). **j** Fraction of cells derived from normal (red) or tumour tissue (green) for each B-cell subcluster in each cancer type. **k** Boxplot showing the prevalence of each cell type stratified for tissue taken from the border, middle or core tumour region in LC. P-values < 0.05 comparing border versus core regions are shown. **l** Violin plot showing transcription factor activities (AUC score) calculated by SCENIC for plasma cell subclusters. **m** Differences in cancer hallmark pathway activities scored per cell by AUCell for B-cell clusters. **n** Violin plot showing pseudotime distribution of different branches indicated in Fig. 5g. **o** Density plots for CRC, LC and OvC along the trajectories of B memory cells of the $IgM^{+}$ and $IgM^{-}$ lineages. **p** Heatmap of expression dynamics of differentially expressed genes along the pseudotime from mature-naïve to class-switched memory B-cell (IgM-). **q** Marker gene dynamics along the pseudotime from mature-naïve to class-switched memory B-cells ($IgM^{-}$) colour-coded as in Fig. 5g. **r** Density plots for CRC, LC and OvC along the differentiation trajectories of $IgM^{-}$ to plasma cells (both the $IgG^{+}$ and $IgA^{+}$ lineages are shown). **s** Heatmap of expression dynamics of differentially expressed genes along the pseudotime from $IgM^{-}$ memory B-cells to $IgG^{+}$ or $IgA^{+}$ plasma cells.

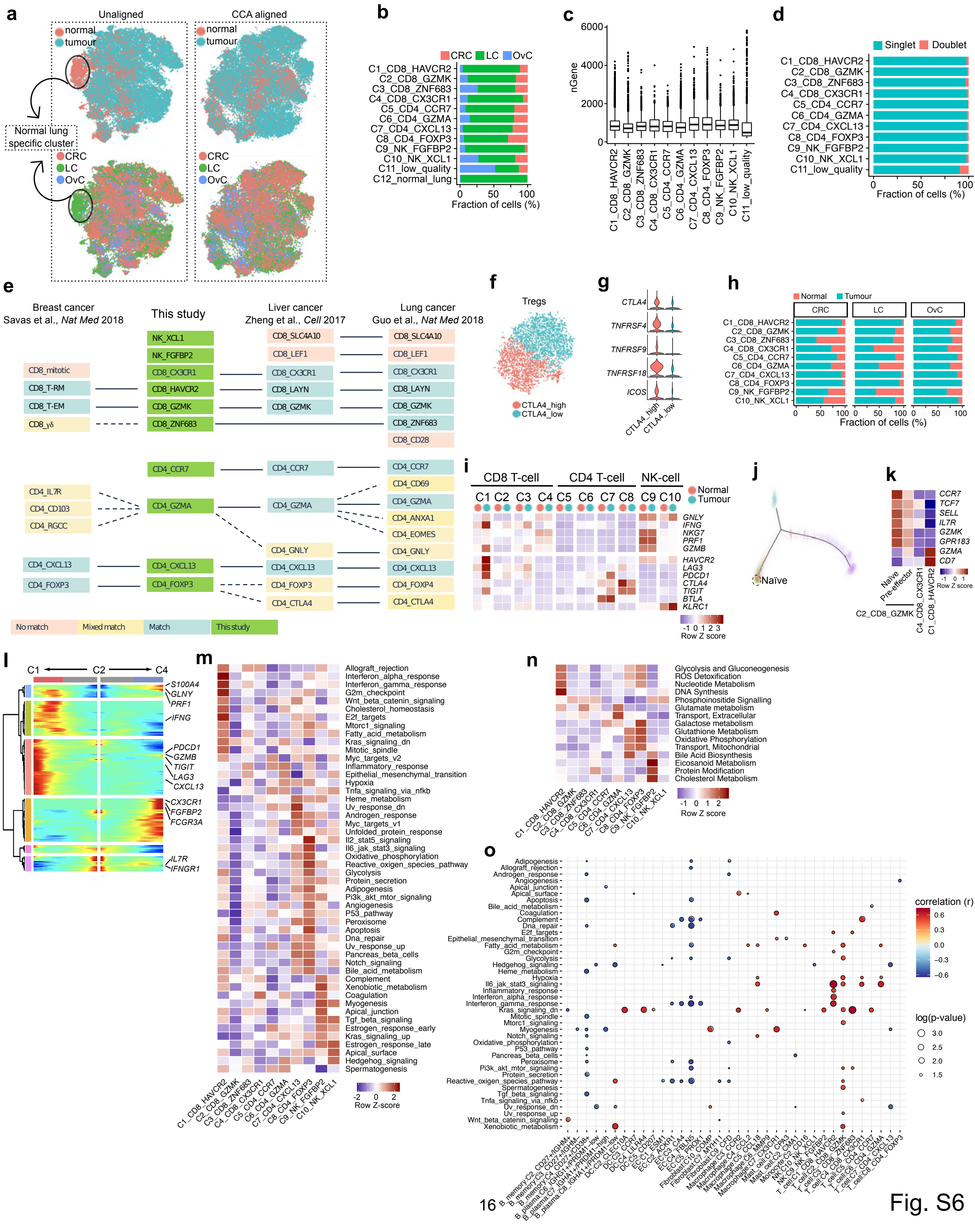
a
Unaligned
CCA aligned
normal
tumour
Normal lung specific cluster
CRC
LC
OvC
b
CRC LC OvC
Fraction of cells (%)
c
nGene
d
Singlet Doublet
Fraction of cells (%)
e
Breast cancer Savas et al., Nat Med 2018
This study
Liver cancer Zheng et al., Cell 2017
Lung cancer Guo et al., Nat Med 2018
No match
Mixed match
Match
This study
f
Tregs
CTLA4_high
CTLA4_low
g
h
Normal Tumour
Fraction of cells (%)
i
CD8 T-cell
CD4 T-cell
NK-cell
Normal
Tumour
Row Z score
j
Naïve
k
Naïve
Pre-effector
C2_CD8_GZMK
Row Z score
l
m
Row Z-score
n
Row Z score
o
correlation (r)
log(p-value)



Fig. S6

**Fig. S6. Profiling of T- and NK-cells**

**a** t-SNE plots showing unaligned (left panels) or CCA-aligned (right panels) clusters colour-coded by sample origin (upper panels), or cancer type (lower panels). Normal lung specific cluster are indicated by circles and arrows. **b** Fraction of cells for T-/NK-cell subclusters (unaligned) from different cancer types. **c** Number of genes detected from different T-cell subclusters (CCA aligned). **d** Fraction of singlet and doublet cells for T-/NK- cell subclusters predicted by DoubletFinder. **e** Comparison of T-/NK-cell subclusters defined in this pan-cancer study and previous breast, liver and lung cancer studies [52,54,55]. **f** t-SNE plot showing subclusters within CD4_FOXP3 Tregs after further subclustering. **g** Violin plot showing marker gene expressions between two Tregs subclusters. **h** Fraction of cells derived from normal (red) or tumour tissue (green) for each T-/NK-cell subcluster in each cancer type. **i** Heatmap of differentially expressed genes for T-/NK-cell subclusters with different sample origins. **j** Trajectories of $CD8^+$ T-cell differentiation. Naïve cells are located at the root of the C2_CD8_GMZK branch (dash line circled). **k** Heatmap of naïve marker gene expression. The naïve subcluster within the C2_CD8_GZMK cluster, as circled in **j**, expresses a number of naïve markers, including *CCR7* and *TCF7*. **l** Heatmap of expression dynamics of differentially expressed genes along the pseudotime of the 2 $CD8^+$ lineages (from C2_CD8_GZMK to C1_CD8_HAVCR1 or C4_CD8_CX3CR1). **m** Differences in cancer hallmark pathway activities scored per cell by AUCell for T-/NK-cell subclusters. **n** Metabolic activity of each T-/NK-cell pheenotype scored by AUCell. **o** Heatmap showing positive (red) and negative (blue) correlations between cancer hallmark pathway activation in cancer cells (Y-axis) and the prevalence of the stromal cell phenotypes.

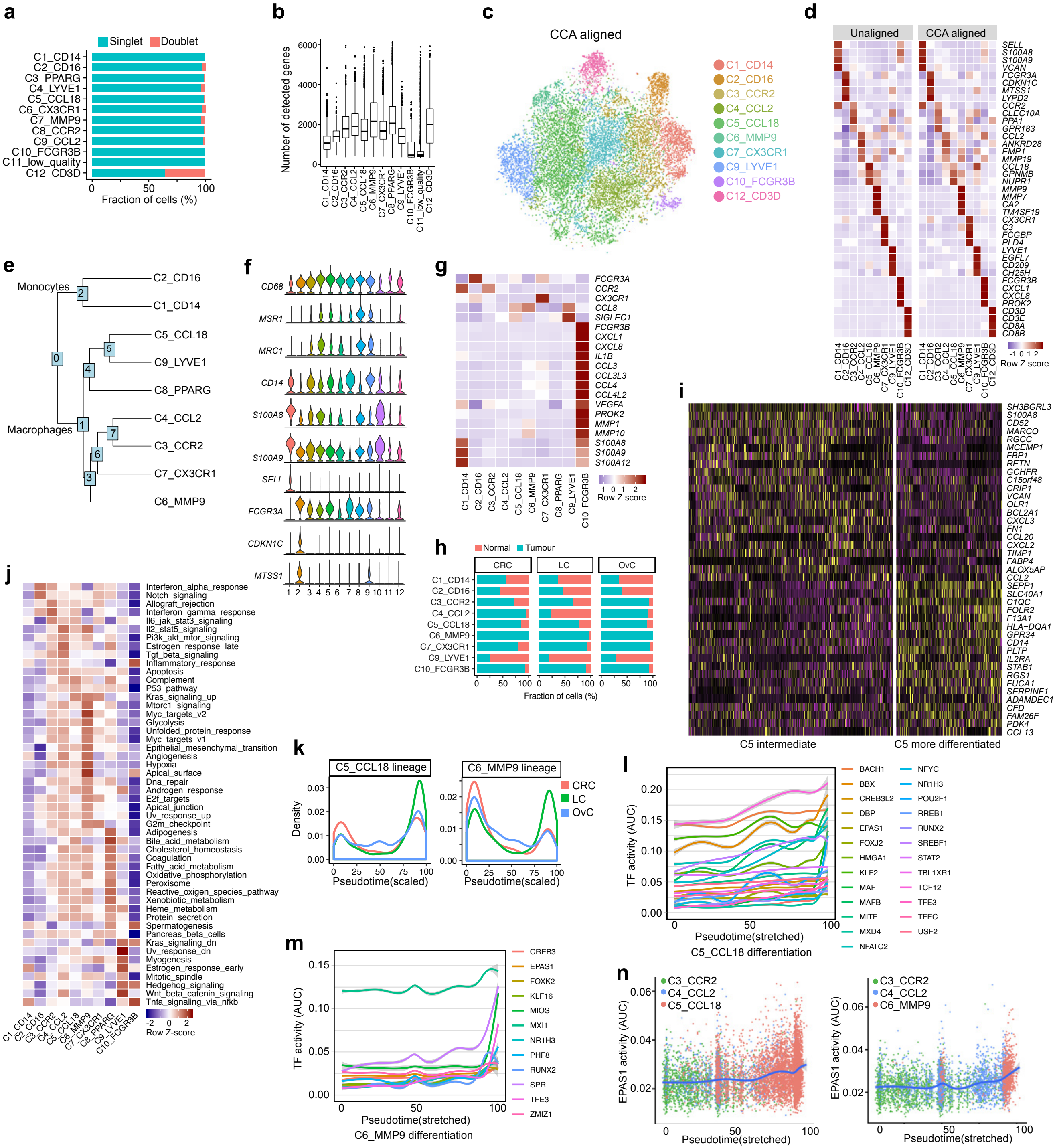
a
Singlet
Doublet
C1_CD14
C2_CD16
C3_PPARG
C4_LYVE1
C5_CCL18
C6_CX3CR1
C7_MMP9
C8_CCR2
C9_CCL2
C10_FCGR3B
C11_low_quality
C12_CD3D
Fraction of cells (%)
b
Number of detected genes
c
CCA aligned
C1_CD14
C2_CD16
C3_CCR2
C4_CCL2
C5_CCL18
C6_MMP9
C7_CX3CR1
C9_LYVE1
C10_FCGR3B
C12_CD3D
d
Unaligned
CCA aligned
SELL
S100A8
S100A9
VCAN
FCGR3A
CDKN1C
MTSS1
LYPD2
CCR2
CLEC10A
PPA1
GPR183
CCL2
ANKRD28
EMP1
MMP19
CCL18
GPNMB
NUPR1
MMP9
MMP7
CA2
TM4SF19
CX3CR1
C3
FCGBP
PLD4
LYVE1
EGFL7
CD209
CH25H
FCGR3B
CXCL1
CXCL8
PROK2
CD3D
CD3E
CD8A
CD8B
Row Z score
e
Monocytes
Macrophages
C2_CD16
C1_CD14
C5_CCL18
C9_LYVE1
C8_PPARG
C4_CCL2
C3_CCR2
C7_CX3CR1
C6_MMP9
f
CD68
MSR1
MRC1
CD14
S100A8
S100A9
SELL
FCGR3A
CDKN1C
MTSS1
g
FCGR3A
CCR2
CX3CR1
CCL8
SIGLEC1
FCGR3B
CXCL1
CXCL8
IL1B
CCL3
CCL3L3
CCL4
CCL4L2
VEGFA
PROK2
MMP1
MMP10
S100A8
S100A9
S100A12
Row Z score
h
Normal
Tumour
CRC
LC
OvC
Fraction of cells (%)
i
SH3BGRL3
S100A8
CD52
MARCO
RGCC
MCEMP1
FBP1
RETN
GCHFR
C15orf48
CRIP1
VCAN
OLR1
BCL2A1
CXCL3
FN1
CCL20
CXCL2
TIMP1
FABP4
ALOX5AP
CCL2
SEPP1
SLC40A1
C1QC
FOLR2
F13A1
HLA−DQA1
GPR34
CD14
PLTP
IL2RA
STAB1
RGS1
FUCA1
SERPINF1
ADAMDEC1
CFD
FAM26F
PDK4
CCL13
C5 intermediate
C5 more differentiated
j
Interferon_alpha_response
Notch_signaling
Allograft_rejection
Interferon_gamma_response
Il6_jak_stat3_signaling
Il2_stat5_signaling
Pi3k_akt_mtor_signaling
Estrogen_response_late
Tgf_beta_signaling
Inflammatory_response
Apoptosis
Complement
P53_pathway
Kras_signaling_up
Mtorc1_signaling
Myc_targets_v2
Glycolysis
Unfolded_protein_response
Myc_targets_v1
Epithelial_mesenchymal_transition
Angiogenesis
Hypoxia
Apical_surface
Dna_repair
Androgen_response
E2f_targets
Apical_junction
Uv_response_up
G2m_checkpoint
Adipogenesis
Bile_acid_metabolism
Cholesterol_homeostasis
Coagulation
Fatty_acid_metabolism
Oxidative_phosphorylation
Peroxisome
Reactive_oxigen_species_pathway
Xenobiotic_metabolism
Heme_metabolism
Protein_secretion
Spermatogenesis
Pancreas_beta_cells
Kras_signaling_dn
Uv_response_dn
Myogenesis
Estrogen_response_early
Mitotic_spindle
Hedgehog_signaling
Wnt_beta_catenin_signaling
Tnfa_signaling_via_nfkb
Row Z-score
k
C5_CCL18 lineage
C6_MMP9 lineage
CRC
LC
OvC
Density
Pseudotime(scaled)
l
TF activity (AUC)
Pseudotime(stretched)
C5_CCL18 differentiation
BACH1
BBX
CREB3L2
DBP
EPAS1
FOXJ2
HMGA1
KLF2
MAF
MAFB
MITF
MXD4
NFATC2
NFYC
NR1H3
POU2F1
RREB1
RUNX2
SREBF1
STAT2
TBL1XR1
TCF12
TFE3
TFEC
USF2
m
TF activity (AUC)
Pseudotime(stretched)
C6_MMP9 differentiation
CREB3
EPAS1
FOXK2
KLF16
MIOS
MXI1
NR1H3
PHF8
RUNX2
SPR
TFE3
ZMIZ1
n
C3_CCR2
C4_CCL2
C5_CCL18
C6_MMP9
EPAS1 activity (AUC)
Pseudotime(stretched)

Fig. S7

**Fig. S7. Profiling of monocytes, macrophages and neutrophils**

**a** Barplot showing fraction of singlet and doublet cells predicted by DoubletFinder for unaligned myeloid clusters. **b** Number of detected genes in unaligned myeloid clusters. **c** t-SNE plot showing myeloid clusters after CCA alignment. **d** Heatmaps comparing marker gene expression signatures of common clusters generated by unaligned (left) and CCA aligned (right) methods, respectively. **e** Phylogenetic analysis of myeloid subclusters using the BuildClusterTree function Seurat. By relating the 'average' cell from each identity class, this analysis confirms the separation between monocyte and macrophage subclusters. **f** Violin plot showing marker gene expressions for unaligned myeloid clusters. **g** Heatmap of some differentially expressed genes in unaligned clusters. The expression of *FCGR3A*(CD16) and *CX3CR1* are higher in both C2_CD16s and C7_CX3CR1s. **h** Fraction of cells derived from normal (red) or tumour tissue (green) for each myeloid cell subcluster in each cancer type. **i** Heatmap of differentially expressed genes between two subclusters of C5_CCL18 macrophages. **j** Differences in cancer hallmark pathway activities scored per cell by AUCell for myeloid cell subclusters. **k** Density plots for CRC, LC and OvC for monocyte-macrophage differentiation along the CCL18 and MMP9 lineages. **l-m** Activation dynamics of transcription factors for terminal differentiation of C5_CCL18 (**l**) and C6_MMP9 (**m**) macrophages. **n** EPAS1 activation dynamics in terminal differentiation of C5_CCL18 (left) or C6_MMP9 (right) macrophages.

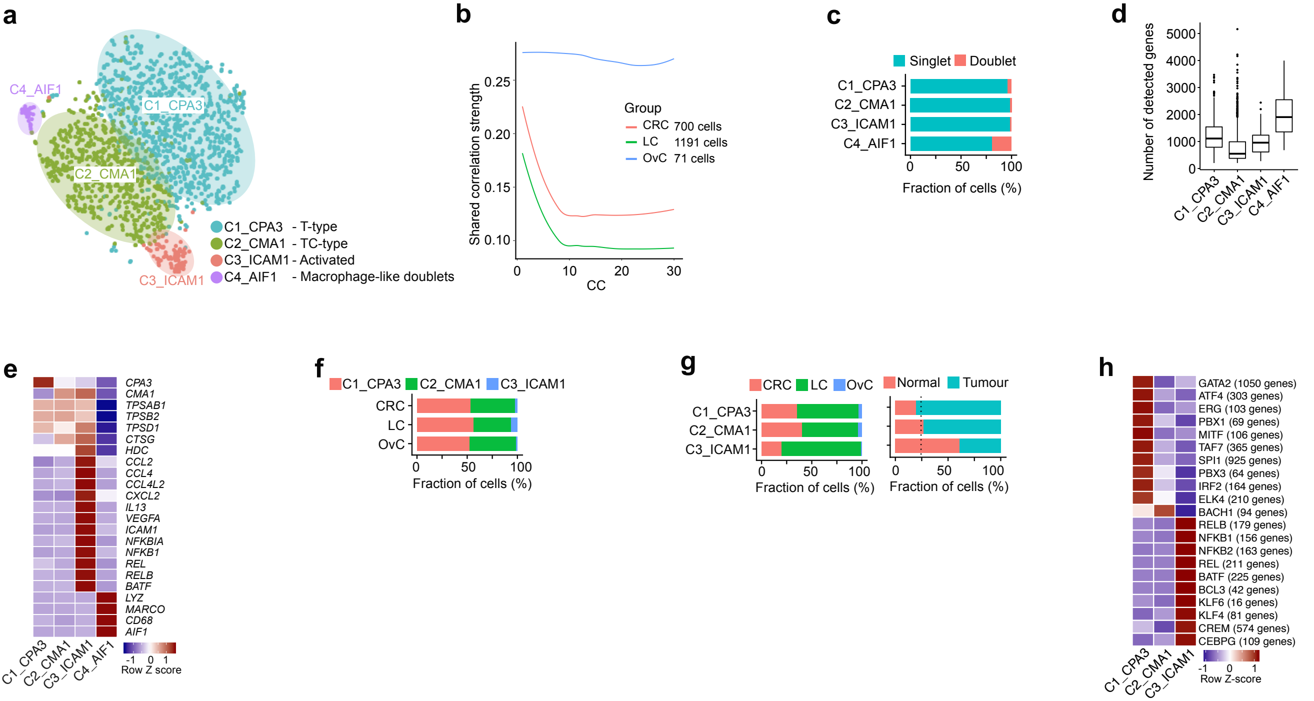
a
C4_AIF1
C1_CPA3
C2_CMA1
C3_ICAM1
C1_CPA3 - T-type
C2_CMA1 - TC-type
C3_ICAM1 - Activated
C4_AIF1 - Macrophage-like doublets
b
Shared correlation strength
Group
CRC 700 cells
LC 1191 cells
OvC 71 cells
CC
c
Singlet
Doublet
Fraction of cells (%)
d
Number of detected genes
e
CPA3
CMA1
TPSAB1
TPSB2
TPSD1
CTSG
HDC
CCL2
CCL4
CCL4L2
CXCL2
IL13
VEGFA
ICAM1
NFKBIA
NFKB1
REL
RELB
BATF
LYZ
MARCO
CD68
AIF1
Row Z score
f
CRC
LC
OvC
Fraction of cells (%)
g
Normal
Tumour
Fraction of cells (%)
h
GATA2 (1050 genes)
ATF4 (303 genes)
ERG (103 genes)
PBX1 (69 genes)
MITF (106 genes)
TAF7 (365 genes)
SPI1 (925 genes)
PBX3 (64 genes)
IRF2 (164 genes)
ELK4 (210 genes)
BACH1 (94 genes)
RELB (179 genes)
NFKB1 (156 genes)
NFKB2 (163 genes)
REL (211 genes)
BATF (225 genes)
BCL3 (42 genes)
KLF6 (16 genes)
KLF4 (81 genes)
CREM (574 genes)
CEBPG (109 genes)
Row Z-score



Fig. S8

**Fig. S8. Characterization of mast cells**

**a** t-SNE plot showing 1,962 mast cells (MCs) colour-coded by 4 clusters, namely C1_CPA3, C2_CMA1, C3_ICAM1 and C4_AIF1, which were generated using the unaligned clustering method. **b** Plot showing biweight midcorrelation score of CCs from different cancer types. The OvC dataset shows poor correlation with other cancers, indicating CCA was not applicable likely due to too few cells derived from OvC. **c** Fraction of singlet and doublet cells predicted by DoubletFinder for mast cell subclusters. **d** Box plot showing number of detected genes for mast cell subclusters. **e** Heatmap for differential expressed genes of each mast cell subcluster. Historically, MCs were classified based on expression of proteases in secretory granules, including tryptases (*TPSAB1*, *TPSB2*), chymase (*CMA1*) and carboxypeptidase A3 (*CPA3*)[99]. We frequently observed C1_CPA3s and C2_CMA1s, while C3_ICAM1s were rare. Tryptases were highly expressed in C1-C3 clusters, but C1_CPA3s were CPA3$^{high}$/CMA1$^{low}$, while *vice versa* C2_CMA1s and C3_ICAM1s were CPA3$^{low}$/CMA1$^{high}$. C3_ICAM1s represented activated MCs that additionally express immune homing factors (*ICAM1*), chemokines and cytokines (*CCL2*, *CCL4*, *CCL4L2* and *IL13*), histamine enzyme (*HDC*) and pro-angiogenic molecules (*VEGFA*). C4_AIF1s had higher predicted doublet rate (**c**), higher detected gene number (d), and higher expression of the macrophage markers (**e**), therefore represented macrophage doublets. **f** Fraction of cells for mast cell phenotypes per cancer type. **g** Fraction of cells for mast cell phenotypes from different cancer types (left), and sample origins (right). C3_ICAM1s were underrepresented in malignant tissue (FDR=$3.3 \times 10^{-23}$), suggesting that tumours do not favour mast cell activation. **h** Heatmap showing transcription factor activity (AUC score) calculated by SCENIC for each mast cell phenotype. It revealed that MC activation in C3_ICAM1s is determined by NF-κB signalling (NFKB1, NFKB2 and REL), while MIFT, a mediator of early MC development[100], was active in C1_CPA3s.

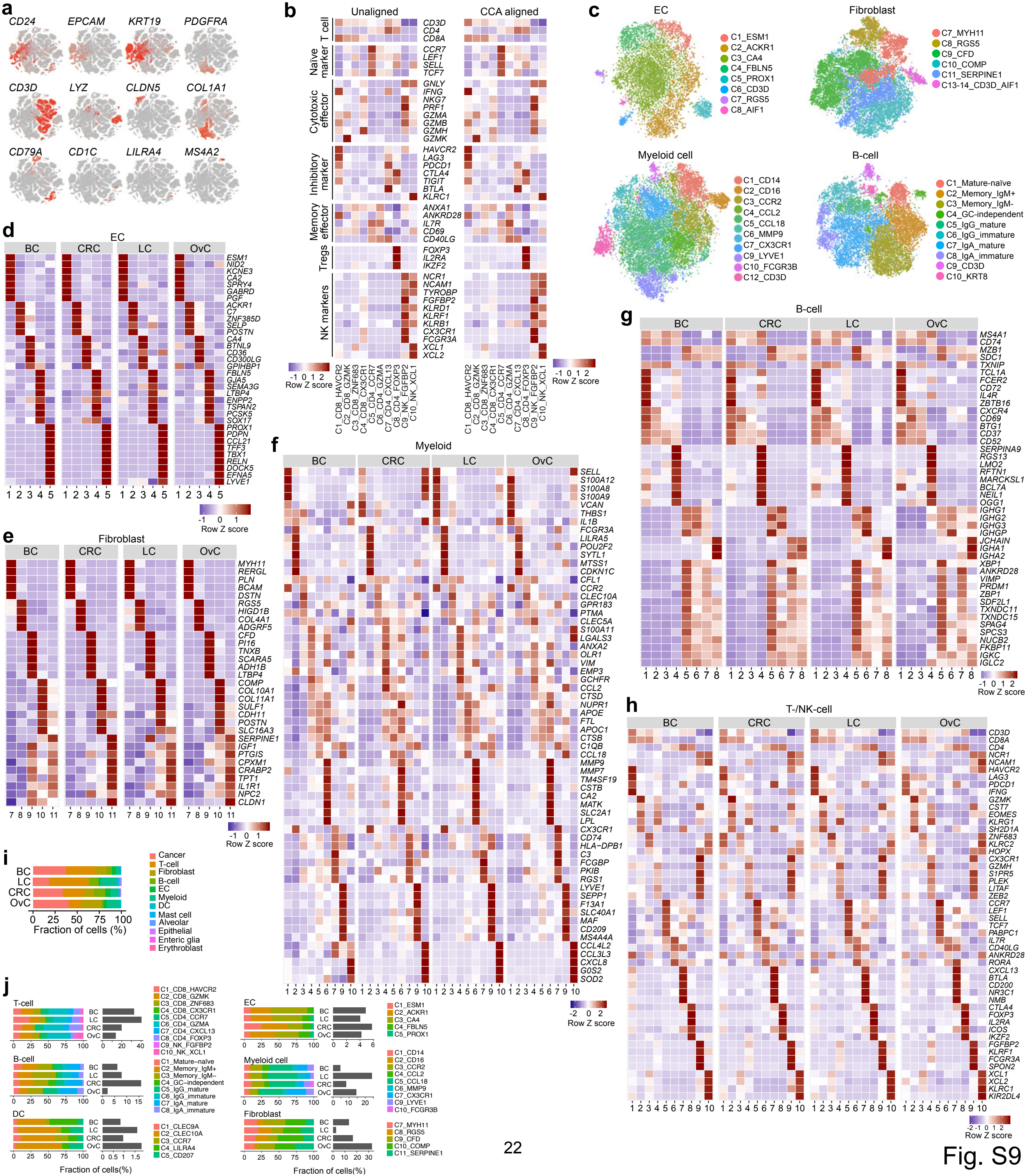
a
CD24
EPCAM
KRT19
PDGFRA
CD3D
LYZ
CLDN5
COL1A1
CD79A
CD1C
LILRA4
MS4A2
b
Unaligned
CCA aligned
T cell
CD3D
CD4
CD8A
Naïve marker
CCR7
LEF1
SELL
TCF7
Cytotoxic effector
GNLY
IFNG
NKG7
PRF1
GZMA
GZMB
GZMH
GZMK
Inhibitory marker
HAVCR2
LAG3
PDCD1
CTLA4
TIGIT
BTLA
KLRC1
Memory effector
ANXA1
ANKRD28
IL7R
CD69
CD40LG
Tregs
FOXP3
IL2RA
IKZF2
NK markers
NCR1
NCAM1
TYROBP
FGFBP2
KLRD1
KLRF1
KLRB1
CX3CR1
FCGR3A
XCL1
XCL2
-1 0 1 2
Row Z score
C1_CD8_HAVCR2
C2_CD8_GZMK
C3_CD8_ZNF683
C4_CD8_CX3CR1
C5_CD4_CCR7
C6_CD4_GZMA
C7_CD4_CXCL13
C8_CD4_FOXP3
C9_NK_FGFBP2
C10_NK_XCL1
c
EC
C1_ESM1
C2_ACKR1
C3_CA4
C4_FBLN5
C5_PROX1
C6_CD3D
C7_RGS5
C8_AIF1
Fibroblast
C7_MYH11
C8_RGS5
C9_CFD
C10_COMP
C11_SERPINE1
C13-14_CD3D_AIF1
Myeloid cell
C1_CD14
C2_CD16
C3_CCR2
C4_CCL2
C5_CCL18
C6_MMP9
C7_CX3CR1
C9_LYVE1
C10_FCGR3B
C12_CD3D
B-cell
C1_Mature-naïve
C2_Memory_IgM+
C3_Memory_IgM-
C4_GC-independent
C5_IgG_mature
C6_IgG_immature
C7_IgA_mature
C8_IgA_immature
C9_CD3D
C10_KRT8
d
EC
BC
CRC
LC
OvC
ESM1
NID2
KCNE3
CA2
SPRY4
GABRD
PGF
ACKR1
C7
ZNF385D
SELP
POSTN
CA4
BTNL9
CD36
CD300LG
GPIHBP1
FBLN5
GJA5
SEMA3G
LTBP4
ENPP2
TSPAN2
PCSK5
SOX17
PROX1
PDPN
CCL21
TFF3
TBX1
RELN
DOCK5
EFNA5
LYVE1
1 2 3 4 5
-1 0 1
Row Z score
e
Fibroblast
MYH11
RERGL
PLN
BCAM
DSTN
RGS5
HIGD1B
COL4A1
ADGRF5
CFD
PI16
TNXB
SCARA5
ADH1B
LTBP4
COMP
COL10A1
COL11A1
SULF1
CDH11
POSTN
SLC16A3
SERPINE1
IGF1
PTGIS
CPXM1
CRABP2
TPT1
IL1R1
NPC2
CLDN1
7 8 9 10 11
-1 0 1
Row Z score
f
Myeloid
SELL
S100A12
S100A8
S100A9
VCAN
THBS1
IL1B
FCGR3A
LILRA5
POU2F2
SYTL1
MTSS1
CDKN1C
CFL1
CCR2
CLEC10A
GPR183
PTMA
CLEC5A
S100A11
LGALS3
ANXA2
OLR1
VIM
EMP3
GCHFR
CCL2
CTSD
NUPR1
APOE
FTL
APOC1
CTSB
C1QB
CCL18
MMP9
MMP7
TM4SF19
CSTB
CA2
MATK
SLC2A1
LPL
CX3CR1
CD74
HLA−DPB1
C3
FCGBP
PKIB
RGS1
LYVE1
SEPP1
F13A1
SLC40A1
MAF
CD209
MS4A4A
CCL4L2
CCL3L3
CXCL8
G0S2
SOD2
1 2 3 4 5 6 7 9 10
-2 0 2
Row Z score
g
B-cell
MS4A1
CD74
MZB1
SDC1
TXNIP
TCL1A
FCER2
CD72
IL4R
ZBTB16
CXCR4
CD69
BTG1
CD37
CD52
SERPINA9
RGS13
LMO2
RFTN1
MARCKSL1
BCL7A
NEIL1
OGG1
IGHG1
IGHG2
IGHG3
IGHGP
JCHAIN
IGHA1
IGHA2
XBP1
ANKRD28
VIMP
PRDM1
ZBP1
SDF2L1
TXNDC11
TXNDC15
SPAG4
SPCS3
NUCB2
FKBP11
IGKC
IGLC2
1 2 3 4 5 6 7 8
-1 0 1 2
Row Z score
h
T-/NK-cell
CD3D
CD8A
CD4
NCR1
NCAM1
HAVCR2
LAG3
PDCD1
IFNG
GZMK
CST7
EOMES
KLRG1
SH2D1A
ZNF683
KLRC2
HOPX
CX3CR1
GZMH
S1PR5
PLEK
LITAF
ZEB2
CCR7
LEF1
SELL
TCF7
PABPC1
IL7R
CD40LG
ANKRD28
RORA
CXCL13
BTLA
CD200
NR3C1
NMB
CTLA4
FOXP3
IL2RA
ICOS
IKZF2
FGFBP2
KLRF1
FCGR3A
SPON2
XCL1
XCL2
KLRC1
KIR2DL4
1 2 3 4 5 6 7 8 9 10
-2 -1 0 1 2
Row Z score
i
BC
LC
CRC
OvC
0 25 50 75 100
Fraction of cells (%)
Cancer
T-cell
Fibroblast
B-cell
EC
Myeloid
DC
Mast cell
Alveolar
Epithelial
Enteric glia
Erythroblast
j
T-cell
C1_CD8_HAVCR2
C2_CD8_GZMK
C3_CD8_ZNF683
C4_CD8_CX3CR1
C5_CD4_CCR7
C6_CD4_GZMA
C7_CD4_CXCL13
C8_CD4_FOXP3
C9_NK_FGFBP2
C10_NK_XCL1
B-cell
C1_Mature–naïve
C2_Memory_IgM+
C3_Memory_IgM–
C4_GC–independent
C5_IgG_mature
C6_IgG_immature
C7_IgA_mature
C8_IgA_immature
DC
C1_CLEC9A
C2_CLEC10A
C3_CCR7
C4_LILRA4
C5_CD207
EC
C1_ESM1
C2_ACKR1
C3_CA4
C4_FBLN5
C5_PROX1
Myeloid cell
C1_CD14
C2_CD16
C3_CCR2
C4_CCL2
C5_CCL18
C6_MMP9
C7_CX3CR1
C9_LYVE1
C10_FCGR3B
Fibroblast
C7_MYH11
C8_RGS5
C9_CFD
C10_COMP
C11_SERPINE1
Fraction of cells(%)



Fig. S9

**Fig. S9. Mapping the breast cancer TME**

**a** t-SNE showing marker gene expressions in breast cancer. **b** Heatmaps comparing marker gene expression of unaligned (left, only BC) and CCA-aligned (right, BC aligned to LC, CRC and OvC) subclusters. **c** t-SNE showing subclusters of ECs, fibroblasts, myeloid cells and B-cells by alignment of 3' and 5'scRNAseq datasets with CCA. **d-h** Heatmaps of conserved marker gene expressions for subclusters of ECs (**d**), fibroblasts (**e**), myeloid cells (**f**), B-cells (**g**), T-/NK-cells (**h**) across different cancer types. **i** Fraction of cells in each cancer type per subcluster. **j** Per cell type, the fraction of cells (grey bars) and cell subclusters (coloured bars) within the tumour is given for each cancer type. BC was enriched for C4_LILRA4_ (FDR = $9.6x10^{-8}$), but had few C5_PROX1 (FDR = $3.4x10^{-5}$).

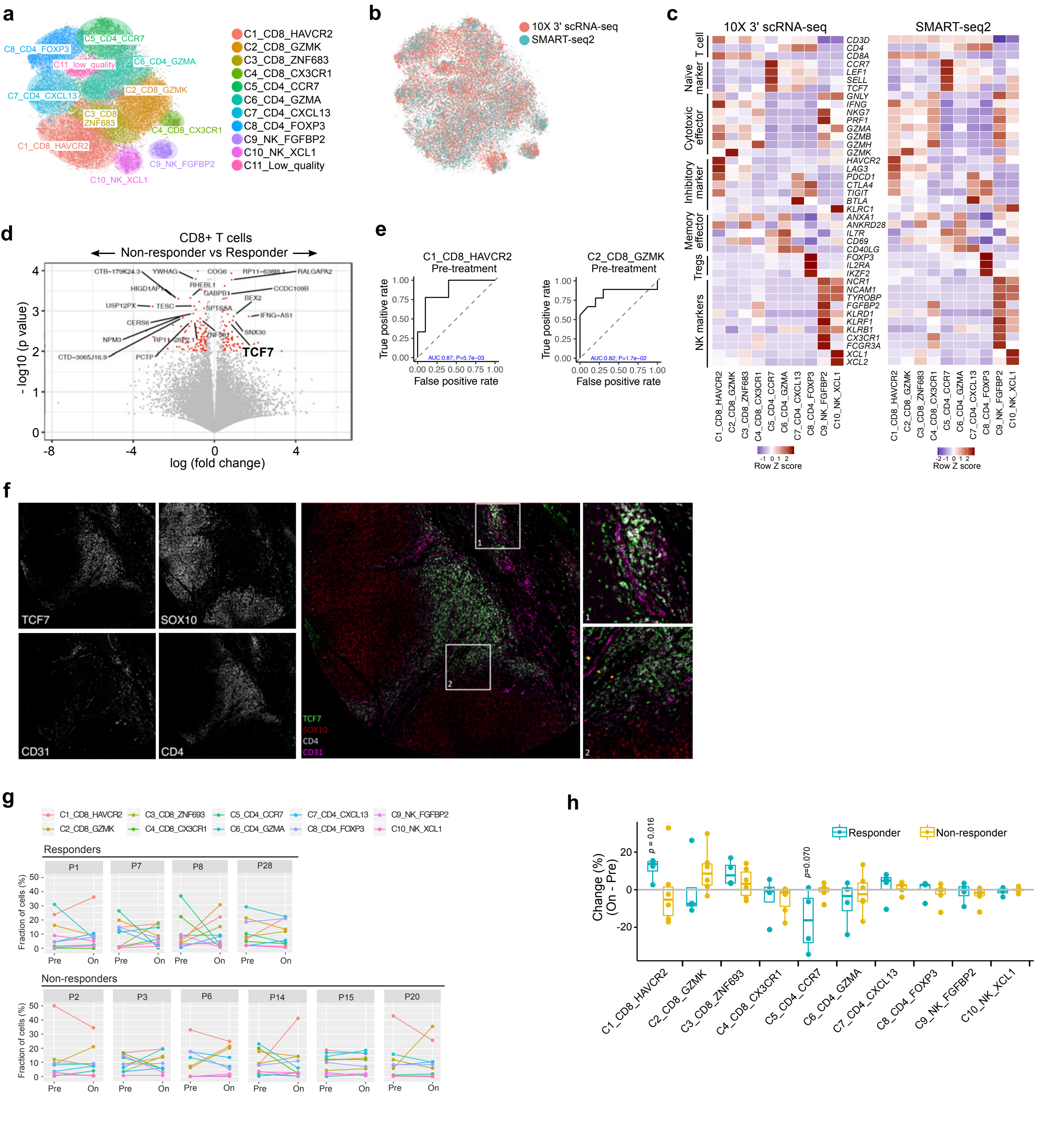



Fig. S10

**Fig. S10. Mapping the melanoma SMART-seq2 dataset**

**a,b** t-SNE plots showing T-/NK-cells clustered by CCA-alignment of SMART-seq2 dataset (melanoma) and 10X 3' scRNA-seq dataset (LC, CRC, OvC), and colour-coded for cluster names (**a**) or technologies (**b**). **c** Heatmaps comparing marker gene expression signatures of CCA-aligned clusters of T-/NK-cells originated from different technologies (left: 10x scRNA-seq, right: SMART-seq2). **d** Volcano plot showing differentially expressed genes in CD8+ T cells from immunotherapy responders and non-responders. **e** Receiver operating characteristic (ROC) analysis was performed to evaluate the predictive effect of C1_CD8_HAVCR2 and C2_CD8_GMZK on response to checkpoint immunotherapy. The area under the ROC curve (AUC) was used to quantify response prediction in pre-treatment biopsies. **f** Images of a representative core, showing 4 single markers (left, greyscale) and a composite RGB-coloured image on the right, with two magnified areas: 1) TCF7$^{+}$ CD4$^{+}$ T-cells (green, white) outside a blood vessel (CD31, purple) at the periphery of the tumour (SOX10, red), and 2) TCF7$^{+}$ CD4$^{+}$ T-cells in the peritumoural area, entering the tumour (SOX10, red) at its interface. **g** Percentage changes of T-/NK-cell clusters in melanoma patient biopsies from pre-treatment (Pre) and on-treatment (On) lesions per patient. The plots were grouped by responder (n=4) and non-responder (n=6). One responder (P4) was excluded due to lack of initial response. **h** Percentage changes of T-/NK-cells in on-treatment (On) and pre-treatment (Pre) lesions per patient. The change (%) = On (%) – Pre (%).

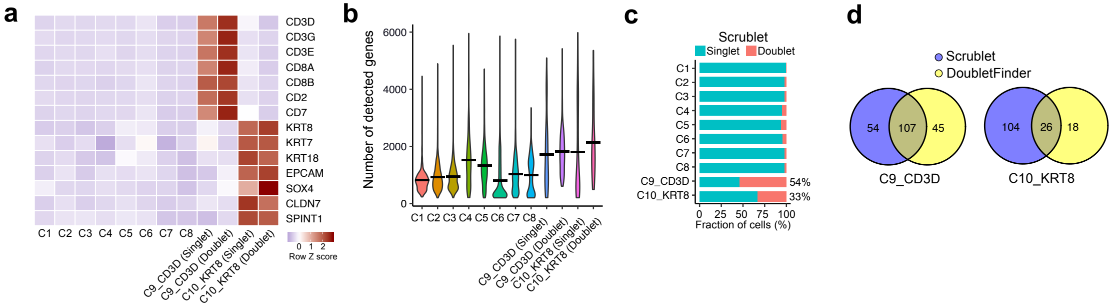
a
CD3D
CD3G
CD3E
CD8A
CD8B
CD2
CD7
KRT8
KRT7
KRT18
EPCAM
SOX4
CLDN7
SPINT1
C1 C2 C3 C4 C5 C6 C7 C8
C9_CD3D (Singlet)
C9_CD3D (Doublet)
C10_KRT8 (Singlet)
C10_KRT8 (Doublet)
0 1 2
Row Z score
b
Number of detected genes
6000
4000
2000
0
C1 C2 C3 C4 C5 C6 C7 C8
C9_CD3D (Singlet)
C9_CD3D (Doublet)
C10_KRT8 (Singlet)
C10_KRT8 (Doublet)
c
Scrublet
Singlet
Doublet
C1
C2
C3
C4
C5
C6
C7
C8
C9_CD3D
C10_KRT8
54%
33%
0 25 50 75 100
Fraction of cells (%)
d
Scrublet
DoubletFinder
54
107
45
C9_CD3D
104
26
18
C10_KRT8

Fig. S11

**Fig. S11. Doublet cluster prediction for B-cell subclusters**

**a** Heatmap of B-cell clusters (C1 to C10) showing the expression of T-cell and cancer cell marker genes. Clusters C9 and C10 were further divided for singlet or doublet subclusters as predicted by DoubletFinder, and cells in these subclusters expressed similarly high levels of T-cell and cancer cell marker genes, respectively. **b** Number of genes detected in the B-cell clusters. The C9 and C10 clusters were further divided for singlet or doublet subclusters as predicted by DoubletFinder. These subclusters had a similarly high level of detected genes as compared to C1-C8 clusters. **c** Percentage of doublets predicted by Scrublet. The C9 and C10 clusters have a higher percentage of doublet cells as compared to C1-C8 clusters, in consistence with the DoubletFinder results. **d** Overlap in cells predicted to be doublets by DoubletFinder and Scrublet, both for cells in the C9 and C10 clusters. This revealed that both tools often identified different cells to represent doublets, supporting the concept that all cells within these clusters should be considered doublet cells.

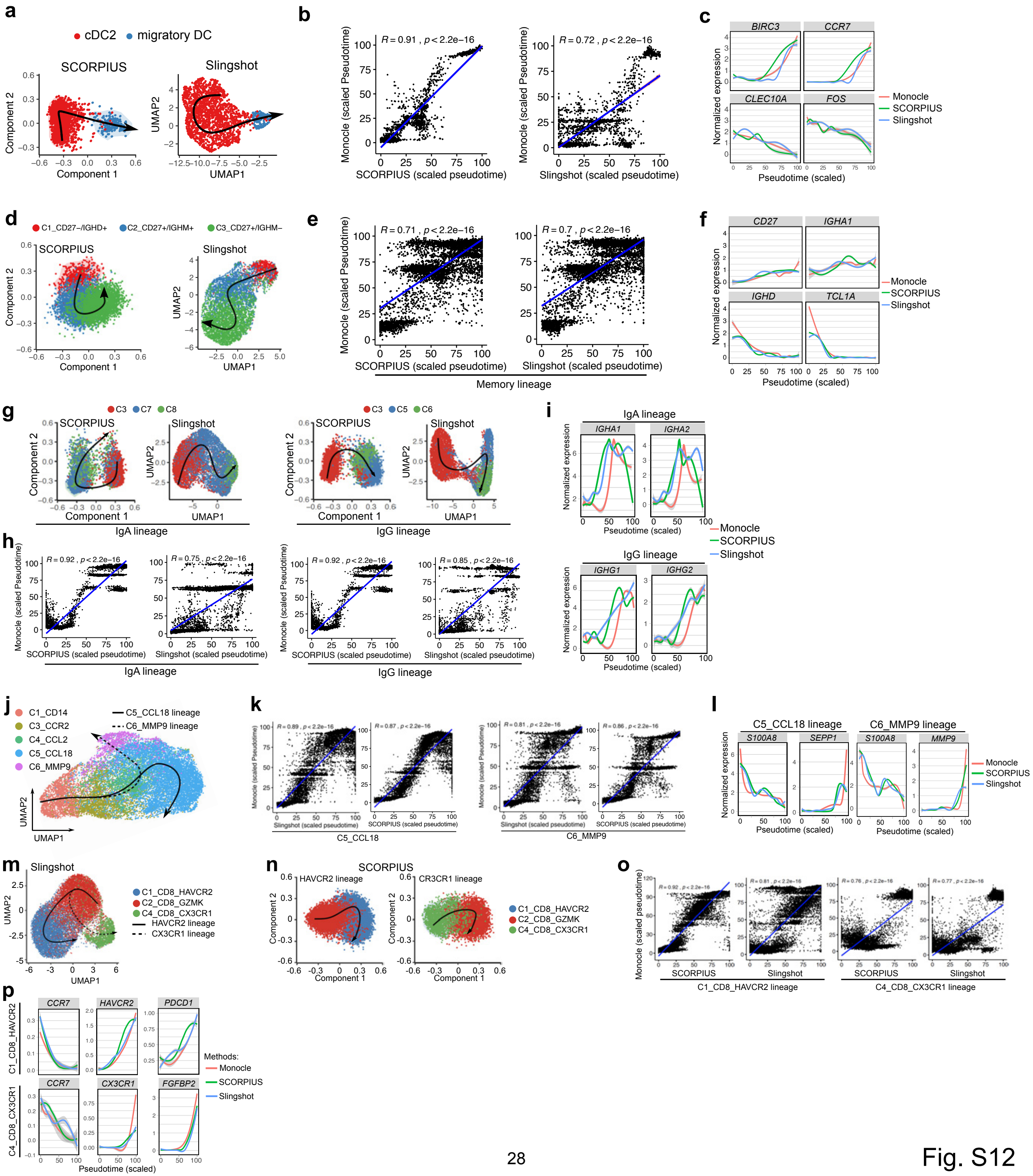
a
cDC2
migratory DC
SCORPIUS
Slingshot
Component 1
Component 2
UMAP1
UMAP2
b
Monocle (scaled Pseudotime)
SCORPIUS (scaled pseudotime)
Slingshot (scaled pseudotime)
c
BIRC3
CCR7
CLEC10A
FOS
Normalized expression
Pseudotime (scaled)
Monocle
SCORPIUS
Slingshot
d
C1_CD27−/IGHD+
C2_CD27+/IGHM+
C3_CD27+/IGHM−
e
Memory lineage
f
CD27
IGHA1
IGHD
TCL1A
g
C3
C7
C8
C5
C6
IgA lineage
IgG lineage
i
IGHA1
IGHA2
IGHG1
IGHG2
h
j
C1_CD14
C3_CCR2
C4_CCL2
C5_CCL18
C6_MMP9
C5_CCL18 lineage
C6_MMP9 lineage
k
C5_CCL18
C6_MMP9
l
S100A8
SEPP1
MMP9
m
C1_CD8_HAVCR2
C2_CD8_GZMK
C4_CD8_CX3CR1
HAVCR2 lineage
CX3CR1 lineage
n
CR3CR1 lineage
o
Monocle (scaled pseudotime)
C1_CD8_HAVCR2 lineage
C4_CD8_CX3CR1 lineage
p
HAVCR2
PDCD1
CX3CR1
FGFBP2
Methods:



Fig. S12

**Fig. S12. Validation of trajectory inference results using Monocle, SCORPIUS and Slingshot**

**a** Trajectories of cDC2 to migratory DC differentiation by SCORPIUS and Slingshot. **b** Correlation between cells ordered along their pseudotime using Monocle versus those ordered by SCORPIUS and Slingshot, respectively. **c** Expression of marker genes *BIRC3*, *CCR7*, *CLEC10A* and *FOS* along the DC lineage. **d** Trajectories of memory B-cells by SCORPIUS and Slingshot. **e** Correlation between cells ordered along their pseudotime using Monocle versus those ordered by SCORPIUS and Slingshot. **f** Expression of marker genes *CD27*, *IGHA1*, *IGHD* and *TCL1A* along the memory B-cell lineage. **g** Trajectories of plasma cell differentiation by SCORPIUS and Slingshot. **h** Correlation between cells ordered along their pseudotime using Monocle versus those ordered by SCORPIUS and Slingshot. **i** Expression of marker genes along the pseudotime of the IgA lineage (*IGHA1*, *IGHA2*) and IgG lineage (*IGHG1*, *IGHG2*). **j** Branched trajectories of myeloid cells by Slingshot. Trajectories are separated for the CCL18 and MMP9 lineage. **k** Correlation between cells ordered along their pseudotime using Monocle versus those ordered by SCORPIUS and Slingshot, respectively. **l** Expression of marker genes along the C5_CCL18 lineage and C6_MMP9 per method. **m-n** Trajectories of CD8+ T-cells generated by Slingshot(**m**) and SCORPIUS(**n**). As SCORPIUS cannot handle branched trajectories, the HAVCR2 and CX3CR1 lineage were analysed separately with SCORPIUS. **o** Correlation between cells ordered along their pseudotime using Monocle versus those ordered using Slingshot and SCORPIUS, respectively, for the HAVCR2 (C1_CD8_HAVCR2) lineage and the CX3CR1 (C4_CD8_CX3CR1) lineage. **p** Expression of marker genes *CCR7*, *HAVCR2*, *PDCD1* along the HAVCR2 lineage and of *CCR7*, *CX3CR1* and *FGFBP2* along the CX3CR1 lineage.

| Patient number | Tumor type | Gender | Age (years) | Stage | TNM classification | Pathological subtype | Molecular status |
|---|---|---|---|---|---|---|---|
| LC_1 | LC | Female | 70 | IIA | pT2bN0M0 | Squamous cell carcinoma | |
| LC_2 | LC | Male | 86 | IB | pT2bN0M0 | Squamous cell carcinoma | |
| LC_3 | LC | Male | 68 | IIIB | pT4N2M0 | Adenocarcinoma | |
| LC_4 | LC | Female | 64 | IIB | pT2aN1M0 | Adenocarcinoma | |
| LC_5 | LC | Male | 60 | IA3 | pT1cN0M0 | Large cell carcinoma | |
| LC_6 | LC | Male | 65 | IIIA | pT4N1M0 | Adenocarcinoma | |
| LC_7 | LC | Male | 60 | IB | pT2aN0M0 | Squamous cell carcinoma | |
| LC_8 | LC | Female | 55 | IIB | pT3N0M0 | Pleiomorphic carcinoma | |
| OvC_1 | OvC | Female | 73 | IIIC | pT3cNxM0 | High grade serous carcinoma | |
| OvC_2 | OvC | Female | 55 | IVB | pT3cNxM1b | High grade serous carcinoma | |
| OvC_3 | OvC | Female | 61 | IVB | pT3cNxM1b | High grade serous carcinoma | BRCA+ |
| OvC_4 | OvC | Female | 83 | IVB | pT3cNxM1b | High grade serous + clear cell carcinoma (mix) | |
| OvC_5 | OvC | Female | 65 | IA | pT1cN0M0 | High grade serous carcinoma | |
| CRC_1 | CRC | Female | 81 | IIB | pT4aN0M0 | Right caecum, global moderately differentiated adenocarcinoma with mixed glandulair, mucinous growth pattern, moderate budding | MSI-high |
| CRC_2 | CRC | Female | 86 | IIIB | pT3N1bM0 | Left rectosigmoid, moderately differentiated adenocarcinoma NST | MSS |
| CRC_3 | CRC | Female | 50 | IVA | pT4aN1aM1a | Left sigmoid, moderately differentiated adenocarcinoma | MSS |
| CRC_4 | CRC | Male | 81 | I | pT2N0M0 | Left sigmoid, moderately differentiated adenocarcinoma | MSS |
| CRC_5 | CRC | Male | 52 | IIA | pT3N0L1 | Left sigmoid, moderately differentiated adenocarcinoma | MSS |
| CRC_6 | CRC | Female | 77 | IIIB | pT3N1aM0 | Right caecum,moderately differentiated adenocarcinoma NOS | MSS |
| CRC_7 | CRC | Male | 84 | IIA | pT3N0L1 | Right ascending,moderately differentiated adenocarcinoma | MSS |
| BC_1 | BC | Female | 41 | III | pT1cN0M0 | Invasive ductal carcinoma | HER2 positive |
| BC_2 | BC | Female | 50 | III | pT2N0M0 | Invasive ductal carcinoma | Triple negative |
| BC_3 | BC | Female | 55 | III | pT3N0M0 | Invasive ductal carcinoma | Triple negative |
| BC_4 | BC | Female | 88 | III | pT1N0M0 | Invasive ductal carcinoma | Triple negative |
| BC_5 | BC | Female | 62 | III | pT2N1aM0 | Invasive apocrine carcinoma | Triple negative |
| BC_6 | BC | Female | 75 | III | pT2N0M0 | Invasive ductal carcinoma | HER2 positive |
| BC_7 | BC | Female | 80 | III | pT3N0M0 | Metaplastic carcinoma | Triple negative |
| BC_8 | BC | Female | 61 | III | pT3N0M0 | Invasive ductal carcinoma | HER2 positive |
| BC_9 | BC | Female | 43 | III | pT2N1M0 | Invasive ductal carcinoma | Luminal-HER2+ |
| BC_10 | BC | Female | 51 | III | pT2N0M0 | Invasive ductal carcinoma | Triple negative |
| BC_11 | BC | Female | 47 | III | pT1cN0M0 | Invasive ductal carcinoma | Triple negative |
| BC_12 | BC | Female | 78 | III | pT2NxM0 | Invasive ductal carcinoma | Triple negative |
| BC_13 | BC | Female | 62 | III | pT2N1M0 | Invasive ductal carcinoma | Luminal B-like |
| BC_14 | BC | Female | 48 | II | pT2N1M0 | Invasive ductal carcinoma | Luminal A-like |
| BC_15 | BC | Female | 66 | III | pT1N0M0 | Invasive ductal carcinoma | Triple negative |
| BC_16 | BC | Female | 60 | III | pT2N1M0 | Invasive ductal carcinoma | Luminal B –like |

**Supplementary Table S1**: Characteristics of the patients included in this study. Included cancer types are lung cancer (LC), ovarian cancer (OvC), colorectal cancer (CRC) and breast cancer (BC). Microsatellite Instable (MSI), microsatellite stable (MSS), Human epidermal growth factor receptor 2 (HER2). Please see Table S3 for tumour mutation signatures.

| Patient number | Cancer type | 10X version | Cells | Sample type | Tumour site | UMIs | Saturation (%) | Reads |
|---|---|---|---|---|---|---|---|---|
| LC_1 | LC | 3' V1 | 175 | Tumour | Core | 799,729 | 97.0 | 52,617,854 |
| LC_1 | LC | 3' V1 | 251 | Tumour | Core | 1,268,737 | 96.6 | 67,690,218 |
| LC_1 | LC | 3' V1 | 98 | Tumour | Middle | 616,179 | 98.1 | 80,569,121 |
| LC_1 | LC | 3' V1 | 302 | Tumour | Middle | 1,489,983 | 95.9 | 70,372,636 |
| LC_1 | LC | 3' V1 | 294 | Tumour | Border | 1,343,488 | 96.2 | 62,162,902 |
| LC_1 | LC | 3' V1 | 353 | Tumour | Border | 1,657,136 | 95.7 | 62,589,463 |
| LC_2 | LC | 3' V1 | 630 | Tumour | Middle | 3,000,702 | 93.2 | 72,138,927 |
| LC_2 | LC | 3' V1 | 31 | Tumour | Middle | 197,373 | 99.1 | 62,127,483 |
| LC_2 | LC | 3' V1 | 122 | Tumour | Border | 808,401 | 97.4 | 55,884,196 |
| LC_2 | LC | 3' V1 | 356 | Normal | na | 1,448,126 | 96.9 | 86,230,103 |
| LC_2 | LC | 3' V1 | 1,541 | Tumour | Border | 5,726,992 | 91.7 | 106,961,758 |
| LC_2 | LC | 3' V1 | 119 | Tumour | Core | 600,577 | 98.1 | 58,982,416 |
| LC_3 | LC | 3' V2 | 4,123 | Tumour | Border | 10,382,153 | 34.1 | 57,353,853 |
| LC_3 | LC | 3' V2 | 4,379 | Tumour | Middle | 16,341,942 | 66.6 | 269,417,085 |
| LC_3 | LC | 3' V2 | 3,517 | Tumour | Core | 16,849,364 | 82.4 | 577,327,377 |
| LC_3 | LC | 3' V2 | 4,772 | Normal | na | 23,306,352 | 84.8 | 379,171,303 |
| LC_4 | LC | 3' V2 | 3,333 | Normal | na | 6,237,310 | 28.0 | 28,594,502 |
| LC_4 | LC | 3' V2 | 3,282 | Tumour | Border | 8,821,660 | 41.4 | 57,131,717 |
| LC_4 | LC | 3' V2 | 5,164 | Tumour | Middle | 14,242,253 | 41.0 | 77,945,533 |
| LC_4 | LC | 3' V2 | 5,493 | Tumour | Core | 17,827,572 | 52.3 | 151,492,569 |
| LC_5 | LC | 3' V2 | 3,628 | Tumour | Core | 11,929,493 | 32.6 | 55,851,067 |
| LC_5 | LC | 3' V2 | 5,012 | Tumour | Outer | 12,470,613 | 39.2 | 72,825,969 |
| LC_5 | LC | 3' V2 | 6,124 | Tumour | Middle | 27,558,700 | 81.2 | 463,180,485 |
| LC_5 | LC | 3' V2 | 6,403 | Normal | na | 32,418,218 | 72.1 | 261,340,951 |
| LC_6 | LC | 3' V2 | 1,735 | Tumour | Core | 8,504,629 | 86.9 | 260,888,273 |
| LC_6 | LC | 3' V2 | 1,429 | Tumour | Middle | 7,971,658 | 84.3 | 245,477,754 |
| LC_6 | LC | 3' V2 | 814 | Tumour | Border | 3,292,751 | 92.8 | 289,816,712 |
| LC_6 | LC | 3' V2 | 2,971 | Normal | na | 27,581,832 | 82.6 | 285,371,297 |
| LC_7 | LC | 3' V2 | 2,902 | Tumour | Core | 18,403,635 | 74.1 | 125,811,122 |
| LC_7 | LC | 3' V2 | 3,708 | Tumour | Middle | 24,084,368 | 74.1 | 159,839,357 |
| LC_7 | LC | 3' V2 | 3,495 | Tumour | Border | 23,010,914 | 74.6 | 161,368,762 |
| LC_7 | LC | 3' V2 | 6,254 | Normal | na | 26,569,105 | 67.2 | 132,227,825 |
| LC_8 | LC | 3' V2 | 3,177 | Normal | na | 13,859,675 | 90.5 | 333,385,825 |
| LC_8 | LC | 3' V2 | 2,455 | Tumour | Border | 17,989,693 | 84.6 | 330,208,745 |
| LC_8 | LC | 3' V2 | 359 | Tumour | Middle | 22,717,876 | 85.7 | 333,583,084 |
| LC_8 | LC | 3' V2 | 1,624 | Tumour | Core | 7,118,507 | 94.2 | 363,006,012 |
| OvC_1 | OvC | 3' V2 | 8,623 | Normal | Omentum | 28,052,375 | 73.5 | 350,096,261 |
| OvC_1 | OvC | 3' V2 | 4,348 | Normal | Omentum | 18,068,251 | 82.5 | 339,448,488 |
| OvC_1 | OvC | 3' V2 | 1,432 | Tumour | Peritoneum | 4,355,115 | 93.9 | 250,974,825 |
| OvC_1 | OvC | 3' V2 | 1,501 | Tumour | Peritoneum | 4,958,614 | 92.9 | 280,613,748 |
| OvC_1 | OvC | 3' V2 | 6,351 | Tumour | Ovarium | 31,083,122 | 78.2 | 400,459,655 |
| OvC_2 | OvC | 3' V2 | 5,151 | Tumour | Peritoneum | 25,746,205 | 39.7 | 193,126,055 |
| OvC_3 | OvC | 3' V2 | 5,450 | Tumour | Peritoneum | 26,667,830 | 81.1 | 306,251,034 |
| OvC_4 | OvC | 3' V2 | 6,257 | Tumour | Peritoneum | 44,446,808 | 63.0 | 298,554,801 |
| OvC_5 | OvC | 3' V2 | 1,136 | Tumour | Ovarium | 5,213,201 | 87.5 | 146,559,477 |
| OvC_5 | OvC | 3' V2 | 4,865 | Normal | Ovarium | 19,384,194 | 67.5 | 120,336,815 |
| CRC_1 | CRC | 3' V2 | 2,712 | Tumour | Core | 23,040,771 | 74.1 | 267,375,232 |
| CRC_1 | CRC | 3' V2 | 2,901 | Tumour | Border | 26,281,216 | 72.2 | 275,655,234 |
| CRC_1 | CRC | 3' V2 | 2,666 | Normal | na | 22,077,384 | 67.6 | 161,020,595 |
| CRC_2 | CRC | 3' V2 | 4,936 | Tumour | Core | 23,780,187 | 76.5 | 266,870,550 |
| CRC_2 | CRC | 3' V2 | 4,461 | Tumour | Border | 20,942,817 | 79.5 | 255,400,094 |
| CRC_2 | CRC | 3' V2 | 2,478 | Normal | na | 20,498,399 | 67.8 | 149,987,327 |
| CRC_3 | CRC | 3' V2 | 2,871 | Tumour | Core | 21,895,911 | 68.5 | 173,029,484 |
| CRC_3 | CRC | 3' V2 | 2,323 | Tumour | Border | 16,041,374 | 72.5 | 148,821,159 |
| CRC_3 | CRC | 3' V2 | 1,904 | Normal | na | 8,600,112 | 77.6 | 126,457,430 |
| CRC_4 | CRC | 3' V2 | 3,770 | Tumour | Core | 16,529,472 | 55.0 | 200,491,544 |
| CRC_4 | CRC | 3' V2 | 2,271 | Tumour | Border | 13,847,003 | 56.1 | 158,677,458 |
| CRC_4 | CRC | 3' V2 | 1,997 | Normal | na | 12,355,632 | 64.4 | 163,981,182 |
| CRC_5 | CRC | 3' V2 | 709 | Tumour | Core | 2,362,128 | 83.6 | 73,852,856 |
| CRC_5 | CRC | 3' V2 | 621 | Tumour | Border | 2,802,982 | 64.7 | 69,557,278 |
| CRC_5 | CRC | 3' V2 | 1,261 | Normal | na | 4,250,554 | 77.0 | 102,694,399 |
| CRC_6 | CRC | 3' V2 | 530 | Tumour | Core | 1,280,817 | 69.5 | 60,655,914 |
| CRC_6 | CRC | 3' V2 | 720 | Tumour | Border | 3,609,540 | 54.5 | 97,567,804 |
| CRC_6 | CRC | 3' V2 | 1,119 | Normal | na | 4,621,711 | 65.4 | 119,764,778 |
| CRC_7 | CRC | 3' V2 | 747 | Tumour | Core | 5,046,944 | 46.9 | 75,192,324 |
| CRC_7 | CRC | 3' V2 | 1,054 | Tumour | Border | 8,723,525 | 51.5 | 96,220,238 |
| CRC_7 | CRC | 3' V2 | 2,633 | Normal | na | 14,030,213 | 67.8 | 179,638,932 |
| BC_1 | BC | 5' V2 | 3,921 | Tumour | Biopsy | 23,817,225 | 83.4 | 331,894,515 |
| BC_2 | BC | 5' V2 | 4,876 | Tumour | Biopsy | 16,583,928 | 65.8 | 162,530,552 |
| BC_3 | BC | 5' V2 | 4,251 | Tumour | Biopsy | 20,633,409 | 85.6 | 309,676,287 |
| BC_4 | BC | 5' V2 | 3,558 | Tumour | Biopsy | 13,810,634 | 72.7 | 154,003,539 |
| BC_5 | BC | 5' V2 | 4,472 | Tumour | Biopsy | 22,094,301 | 65.8 | 150,279,221 |
| BC_6 | BC | 5' V2 | 201 | Tumour | Biopsy | 1,310,634 | 83.0 | 39,183,120 |
| BC_7 | BC | 5' V2 | 1,719 | Tumour | Biopsy | 15,089,867 | 71.4 | 146,585,647 |
| BC_8 | BC | 5' V2 | 2,427 | Tumour | Biopsy | 9,087,189 | 69.3 | 151,070,133 |
| BC_9 | BC | 5' V2 | 4,616 | Tumour | Biopsy | 28,428,991 | 66.8 | 265,636,248 |
| BC_10 | BC | 5' V2 | 902 | Tumour | Biopsy | 4,088,085 | 76.3 | 39,349,726 |
| BC_11 | BC | 5' V2 | 4,862 | Tumour | Biopsy | 28,400,652 | 65.9 | 275,631,788 |
| BC_12 | BC | 5' V2 | 362 | Tumour | Biopsy | 1,725,747 | 81.3 | 24,816,828 |
| BC_12 | BC | 5' V2 | 64 | Tumour | Biopsy | 323,793 | 93.5 | 15,193,301 |
| BC_12 | BC | 3' V2 | 1,372 | Tumour | Biopsy | 8,866,716 | 77.5 | 118,432,604 |
| BC_13 | BC | 5' V2 | 3,641 | Tumour | Biopsy | 19,494,370 | 70.3 | 219,414,682 |
| BC_14 | BC | 5' V2 | 4,216 | Tumour | Biopsy | 34,749,191 | 65.8 | 284,244,840 |
| BC_15 | BC | 3' V2 | 4,143 | Tumour | Biopsy | 20,268,164 | 56.9 | 295,259,214 |
| BC_15 | BC | 3' V2 | 347 | Tumour | Biopsy | 2,851,324 | 55.9 | 157,883,076 |
| BC_16 | BC | 3' V2 | 269 | Tumour | Biopsy | 1,070,916 | 67.4 | 195,421,321 |

**Supplementary Table S2**: Sequencing metrics of the samples included in this study. Cancer types are lung cancer (LC), colorectal cancer (CRC), ovarian cancer (OvC) and breast cancer (BC). Single cell suspensions were converted to single cell RNA sequencing libraries using 10X genomics kits with 3’ V1 3’ V2 or 5’ V2 chemistry; UMI: unique molecular identifier, equivalent to a unique detected transcript; sequencing saturation and total number of sequenced reads.
